# Remodeling of Trigeminal Neuron Subtypes in Chronic Orofacial Pain

**DOI:** 10.64898/2026.11.10.763722

**Authors:** Lauren F. Cook, Minghan Hu, Daniel Izquierdo, Sijung Yun, Bradford E. Hall, Kenneth M. Yamada, Ashok B. Kulkarni

**Affiliations:** Functional Genomics Section, National Institute of Dental and Craniofacial Research, National Institutes of Health, Bethesda, MD; Cell Biology Section, National Institute of Dental and Craniofacial Research, National Institutes of Health, Bethesda, MD; Genomics and Computational Biology Core, National Institutes of Health, Bethesda, MD; Predictiv Care, Inc., Mountain View, CA

**Author notes:** These authors contributed equally (Lauren F. Cook, Minghan Hu). Co-corresponding authors, **Address for Correspondence,** Ashok Kulkarni, PhD, 30 Convent Drive, Building 30, Room 130, NIDCR, NIH, Bethesda, MD 20892; Kenneth Yamada, MD, PhD, 30 Convent Drive, Building 30, Room 430, NIDCR, NIH, Bethesda, MD 20892.

**Keywords:** Ganglia, Sensory, Cranial Nerve Injuries, RNA-Seq, Single-Cell Gene Expression Analysis, Neuronal Plasticity, Mechanoreceptors

## Abstract

Chronic orofacial pain conditions, including trigeminal neuralgia, post-traumatic trigeminal neuropathic pain, and neuropathic temporomandibular disorder-related pain, remain clinically intractable. The trigeminal ganglion (TG) contains primary sensory neurons that convey sensory information from the face and oral cavity, making TG subtype-specific injury responses central to persistent craniofacial pain. However, how individual TG neuronal subtypes respond to nerve injury during chronic pain remains poorly understood. Prior single-cell injury datasets often used nerve transection models for neuropathic pain that induce strong, transient injury and regeneration-associated states. To define TG transcriptional remodeling in a model of trigeminal neuropathic pain, we performed single-nucleus RNA sequencing of mouse TG after infraorbital nerve chronic constriction injury (IoN-CCI), comparing acute (Day 7) and chronic (Day 21) stages. Analyzing 17,801 nuclei, we identified an initial injury-associated neuronal expression pattern enriched for regeneration biomarkers (*Atf3*, *Sox11*) at Day 7 that was not detected at Day 21. Unlike transection models, in which injured neurons converge on a shared injury-associated transcriptional state, IoN-CCI largely retained recognizable canonical neuronal identities while revealing delayed remodeling at Day 21. Cross-model comparisons with published TG axotomy and DRG sciatic nerve transection datasets showed limited overlap, supporting model- and ganglion-specific remodeling with trigeminal constriction injury. Acute TG responses were characterized by stress and injury-associated transcriptional programs, whereas chronic-stage transcriptional programs were enriched for synaptic/structural remodeling and ion channel reconfiguration. Chronic remodeling was most evident in canonical p-cLTMR2 mechanosensory and *Trpm8*-positive PEP2 cold-sensitive neurons, and RNAscope validated induction of *Tmem45b* and *Rgs8* in *Gfra2*-positive neurons and *Fgf13* and *Kcnmb2* in *Trpm8*-positive neurons. Our findings demonstrate that IoN-CCI drives stage- and neuronal subtype-specific transcriptional remodeling within canonical trigeminal neurons. These findings identify neuronal subtype-specific molecular changes associated with persistent orofacial neuropathic pain and highlight candidates for future therapeutic investigation.

## Introduction

Chronic orofacial pain encompasses a heterogeneous group of conditions affecting the face and oral cavity, including painful temporomandibular disorders, trigeminal neuropathic pain, and idiopathic orofacial pain conditions (International classification of orofacial pain, 1st edition (icop) 2020; Young et al. 2022). Orofacial pain affects approximately 25% of the population, with approximately 10% considered chronic (Benoliel et al. 2019). Neuropathic orofacial pain arises from lesion or disease of the somatosensory nervous system and includes trigeminal neuralgia and other trigeminal neuropathic pain conditions, such as post-traumatic trigeminal neuropathic pain (International classification of orofacial pain, 1st edition (icop) 2020; Korczeniewska et al. 2022; Young et al. 2022). Management of neuropathic orofacial pain remains challenging, and available pharmacologic options, including anticonvulsants and antidepressants, provide incomplete relief for many patients (Gambeta et al. 2020; Korczeniewska et al. 2022; Romero-Reyes et al. 2023). This limited therapeutic efficacy may partly reflect incomplete understanding of the cellular and molecular programs that contribute to persistent pain within defined trigeminal ganglion (TG) neuronal subtypes. Clinical heterogeneity further suggests that persistent orofacial pain may arise from multiple cell-type-specific mechanisms (Khan and Tao 2025; Shinoda et al. 2021).

Infraorbital nerve chronic constriction injury (IoN-CCI) models key features of human orofacial neuropathic pain and is widely used for mechanistic studies (Deseure and Hans 2024; Ding et al. 2017). Whereas dorsal root ganglia (DRG) predominantly innervate somatic territories outside the craniofacial region, the TG innervates facial and oral tissues; both contain sensory neuron populations that encode thermal, mechanical, and chemical stimuli (Nguyen et al. 2017). Major sensory neuron subtypes include neurofilament-expressing mechanosensory neurons, C-fiber low-threshold mechanoreceptors (cLTMRs), and nociceptor classes — peptidergic, nonpeptidergic, and somatostatin-positive (Megat et al. 2019; Nguyen et al. 2017). These populations are relevant to neuropathic pain because nerve injury can cause normally innocuous touch and cooling to evoke pain (MacDonald et al. 2021; Pina et al. 2024; Seal et al. 2009; Xing et al. 2007). However, their subtype-specific roles in orofacial neuropathic pain remain unclear. Although DRG ligation and transection models have defined core injury-associated transcriptional programs shared across sensory neuron classes, TG neurons differ from DRG neurons in subtype composition and gene-expression signatures, suggesting both conserved and ganglion-specific mechanisms (Bhuiyan et al. 2024; Megat et al. 2019; Nguyen et al. 2019; Nguyen et al. 2017; Renthal et al. 2020).

Bulk and targeted TG gene-expression studies identified injury-regulated genes but lacked neuronal subtype resolution (Cui et al. 2023; Korczeniewska et al. 2018). Foundational single-cell and single-nucleus RNA-sequencing (sc/snRNA-seq) injury studies have used axotomy or transection-type models, which sever axons and elicit stereotyped injury and regenerative programs in injured neurons (Nguyen et al. 2019; Renthal et al. 2020). This dominant injury program can obscure subtype-specific identity, limiting direct extrapolation to the chronic pain model CCI, where injured and canonical populations coexist. By contrast, CCI produces partial axonal damage that spares many neurons from direct injury (Obata et al. 2003), thereby enabling analysis of subtype-specific plasticity in neurons that retain canonical transcriptional identities (Zhang et al. 2022). Canonical afferents are increasingly recognized as active contributors to persistent neuropathic pain through molecular and electrophysiological remodeling (Lovatt et al. 2022; Obata et al. 2003; Tran and Crawford 2020; Zhang et al. 2022). Here, we used snRNA-seq at acute (Day 7) and chronic (Day 21) stages to identify which TG neuron types change after IoN-CCI and how transcriptional remodeling differs between these stages. Collectively, our findings reveal cell-type-specific transcriptional reprogramming within neurons during chronic orofacial neuropathic pain and highlight candidate molecular targets for therapeutic exploration.

## Methods

All experiments involving animals were approved by the NIDCR Institutional Animal Care and Use Committee (ASP 24-1161, 19-894) and conducted in accordance with NIH guidelines. C57BL/6J mice (8–10 weeks; both sexes) underwent unilateral infraorbital nerve chronic constriction injury with contralateral sham surgery. Independent cohorts were used for snRNA-seq (n=24) and RNAscope validation (n=17). For snRNA-seq, eight trigeminal ganglia were pooled per library, generating two matched control/CCI pairs at Day 7 and one at Day 21. RNAscope was analyzed at the mouse level. Transcriptomic analyses used Seurat/MAST. Detailed methods are provided in the Appendix. This study followed ARRIVE 2.0 guidelines.

## Results

### Single-nucleus profiling defines trigeminal ganglion cell-type organization and a transient injury-associated neuronal state after infraorbital nerve constriction

We performed single-nucleus RNA sequencing of trigeminal ganglia after unilateral infraorbital nerve chronic constriction injury (IoN-CCI), collecting ipsilateral IoN-CCI TGs and contralateral sham-control TGs from the same mice at Day 7 or Day 21 post-injury (Fig. 1A). After filtering, 17,801 nuclei were retained for analysis. Clustering and integration quality were assessed across multiple resolutions (Appendix Fig. 1). Unsupervised clustering resolved neuronal and non-neuronal TG populations (Fig. 1B), and we annotated the identifiable clusters into major sensory neuron, glial, vascular, and stromal classes (Nguyen et al. 2017; Yang et al. 2022). Canonical marker expression supported neuronal, glial, vascular, stromal, and sensory neuron subtype identities (Fig. 1C and D).

**Figure 1.**
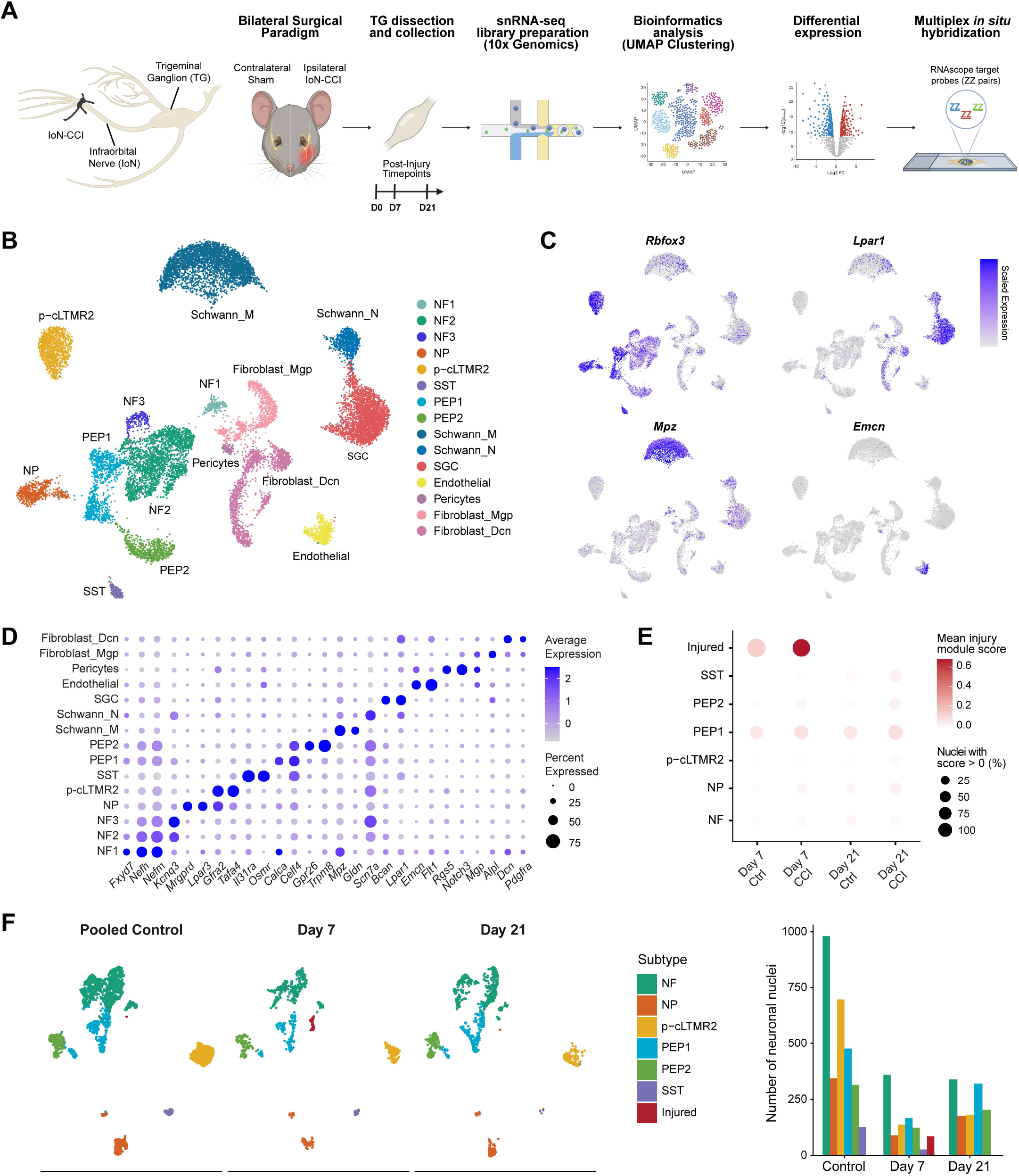
Experimental workflow and single-nucleus transcriptomic atlas of the trigeminal ganglion following IoN-CCI. **(A)** Schematic of the infraorbital nerve chronic constriction injury (IoN-CCI) model and experimental workflow. Day 0 denotes the day of unilateral IoN-CCI and contralateral sham surgery. Trigeminal ganglia were collected at postoperative Day 7 or Day 21, nuclei were isolated, and single-nucleus RNA-seq libraries were prepared using the 10x Genomics platform. Candidate transcripts were selected for validation by multiplex RNAscope in situ hybridization. **(B)** Integrated UMAP of 17,801 nuclei after SCTransform normalization, PCA, and Harmony integration. Unsupervised clustering resolved major neuronal and non-neuronal trigeminal ganglion cell classes. **(C)** Representative marker gene expression projected onto the integrated UMAP. *Rbfox3* labels neurons, *Lpar1* marks satellite glial cells, *Mpz* labels myelinating Schwann cells, and *Emcn* labels endothelial cells. **(D)** Dot plot of canonical marker gene expression across annotated classes. Dot size indicates the proportion of nuclei expressing each gene, and color intensity reflects scaled average expression. **(E)** Injury module score across neuronal subtypes and conditions. Dot size indicates the percentage of nuclei with a module score greater than 0, and color intensity indicates mean injury module score. **(F)** UMAP embedding of neuronal nuclei according to condition, with subtype quantification shown at right. Control nuclei from Day 7 and Day 21 were pooled for visualization. Neuronal nuclei resolved canonical trigeminal ganglion subtypes and a discrete injury-associated neuronal population enriched at Day 7 CCI (red).

We next focused on the neuronal population. Neuron-focused analysis identified the transcriptomically defined NF, NP, SST, PEP1, PEP2, and p-cLTMR2 subtypes, as well as a discrete injury-associated neuronal population, visualized across conditions in the pooled neuron UMAP (Fig. 1F). NF1-NF3 subclusters identified in the full atlas were combined into a single NF class for downstream timepoint-specific analyses. The injury-associated population was enriched at Day 7 CCI but was not detected at Day 21 CCI, whereas major sensory-neuron subtype identities remained recognizable. Consistent with this classification, an injury module score based on established regeneration-associated genes was highest in the Day 7 CCI injury-associated cluster, while the other neuronal subtypes showed substantially lower module activity (Fig. 1E) (Nguyen et al. 2019; Renthal et al. 2020; Seijffers et al. 2007). Additional marker analyses supported enrichment of injury-associated genes and heterogeneous canonical sensory-neuron signatures within this population (Appendix Fig. 2A–E). Quantification confirmed this pattern: injury-associated neurons were nearly absent in controls, accounted for 86 of 991 neuronal nuclei at Day 7 CCI, and were not detected at Day 21 CCI. Inferred subtype assignments of the injured neurons were analyzed across several sensory-neuron subtypes, while p-cLTMR2 and PEP2 nuclei were predominantly classified in canonical states at Day 7 (Appendix Fig. 2F,G).

**Figure 2.**
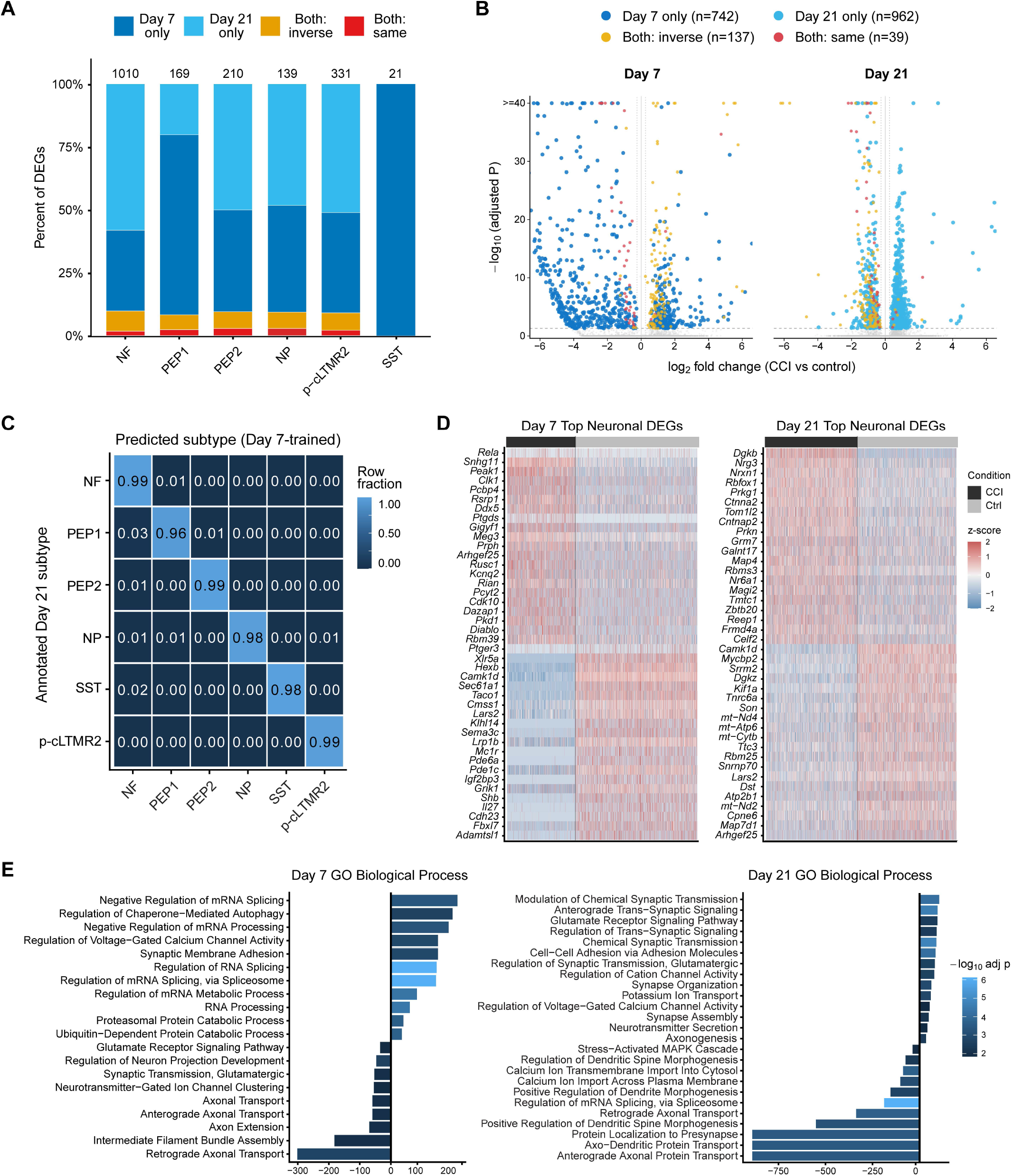
Temporal remodeling of canonical trigeminal neuron transcriptional programs after IoN-CCI. **(A)** Stacked bar plot showing the distribution of differentially expressed genes (DEGs) across canonical neuronal subtypes at Day 7 and Day 21. DEGs were classified as Day 7 only (dark blue), Day 21 only (light blue), shared with opposite-direction regulation (yellow), or shared with same-direction regulation (red). Bars indicate the percentage of DEGs in each category, and numbers above the bars represent the total number of DEG comparisons per subtype. **(B)** Paired volcano plots showing CCI-associated differential gene expression at Day 7 and Day 21. Each point represents a gene–neuronal subtype comparison, plotted according to log₂ fold change (CCI versus control) and −log₁₀ adjusted P value. Colored points correspond to significant DEGs using the same categories as in (A) and gray points represent nonsignificant comparisons. Only comparisons meeting DEG criteria at the displayed timepoint are highlighted. Dashed horizontal and dotted vertical lines indicate the significance thresholds (adjusted P < 0.05 and |log₂ fold change| ≥ 0.25). **(C)** Confusion matrix showing label transfer from Day 7 to Day 21 canonical neuronal subtypes. Values represent row-normalized fractions of Day 21 annotated subtypes assigned to each Day 7-trained predicted label. **(D)** Heatmaps of top pooled neuronal DEGs at Day 7 and Day 21. Columns represent individual nuclei grouped by condition, rows represent genes, and expression values are gene-wise z scores. **(E)** Gene Ontology biological process enrichment of pooled neuronal DEGs at Day 7 and Day 21. Bars show signed combined enrichment scores, with negative values indicating enrichment among downregulated genes and positive values indicating enrichment among upregulated genes. Color intensity represents −log10 adjusted *P* value.

### Time-dependent reorganization of neuronal transcriptional programs after nerve injury

To define how injury-associated transcriptional changes differed over time, we compared Day 7 and Day 21 CCI-regulated differentially expressed genes (DEGs) within each neuronal subtype (Fig. 2A). Gene–subtype comparisons were classified as Day 7 only, Day 21 only, shared with opposite-direction regulation, or shared with same-direction regulation. Most DEGs were identified at only one timepoint, with 742 Day 7-only and 962 Day 21-only comparisons, compared with 176 shared comparisons. PEP1 was dominated by Day 7-specific DEGs, whereas PEP2 and p-cLTMR2 exhibited larger Day 21-specific components. NP showed contributions from both timepoints, while NF neurons had the largest overall number of DEGs, with a substantial Day 21-specific component.

Paired volcano plots further highlighted the distinct transcriptional responses at Day 7 and Day 21 (Fig. 2B). Among the 176 shared gene–subtype comparisons, 137 (77.8%) showed opposite-direction regulation, predominantly exhibiting increased expression following CCI at Day 7 and decreased expression at Day 21. Same-direction genes included axonal, cytoskeletal, and injury-associated genes such as *Nefl*, *Map1a*, *Sptbn1*, *Dst*, *Syne1*, and *Znrf1* (Araki and Wakatsuki 2019; Wakatsuki et al. 2011). The pooled Day 21-specific gene set included ion-channel and sensory-signaling genes such as *Scn9a*, *Cacna2d1*, *Asic3*, *Trpm3*, and *Hcn1* (Fang et al. 2026; Ghosh et al. 2024; Kung et al. 2020; Li and Xu 2011). Cross-timepoint label transfer assessed whether neuronal subtype identity remained recognizable from Day 7 to Day 21 (Fig. 2C). Day 21 subtype labels strongly matched Day 7-based predictions, with minimal cross-subtype assignment. Thus, major neuronal subtype identities remained largely stable from Day 7 to Day 21 despite substantial temporal differences in injury-regulated transcriptional programs. Because SST neurons were sparsely recovered in the Day 21 CCI condition and yielded no Day 21 DEGs under our significance criteria, SST was excluded from subsequent IoN-CCI subtype-specific transcriptional analyses but retained in cross-model summaries.

Gene-wise z-scored heatmaps showed distinct CCI-associated expression patterns at Day 7 and Day 21 (Fig. 2D). The Day 7 signature included genes associated with intracellular trafficking, calcium-dependent signaling, axon guidance/axonal injury, and prostaglandin-related signaling, including *Sec61a1*, *Camk1d*, *Sema3c*, *Prph*, *Ptgds*, and *Ptger3*. The Day 21 signature included altered expression of genes associated with synaptic organization, neuronal connectivity, structural remodeling, RNA processing, and mitochondrial function, including *Nrxn1*, *Cntnap2*, *Grm7*, *Rbfox1*, *Ctnna2*, *Map4*, *Srrm2*, *Son*, *Rbm25*, *Kif1a*, and mitochondrial transcripts (Boccella et al. 2020; Dawes et al. 2018; He et al. 2024; Yang et al. 2025). Functional enrichment identified distinct Day 7 and Day 21 biological processes (Fig. 2E). Signed enrichment showed a shift from acute RNA processing/splicing, protein turnover, calcium signaling, and axonal remodeling to chronic synaptic and neurotransmitter signaling, ion transport, and structural remodeling. RNA-processing and splicing pathways also reversed from CCI-upregulated enrichment at Day 7 to CCI-downregulated enrichment at Day 21. Additional analyses further supported time-dependent transcriptional remodeling and showed that temporal divergence was robust to unequal nucleus recovery (Appendix Fig. 3).

**Figure 3.**
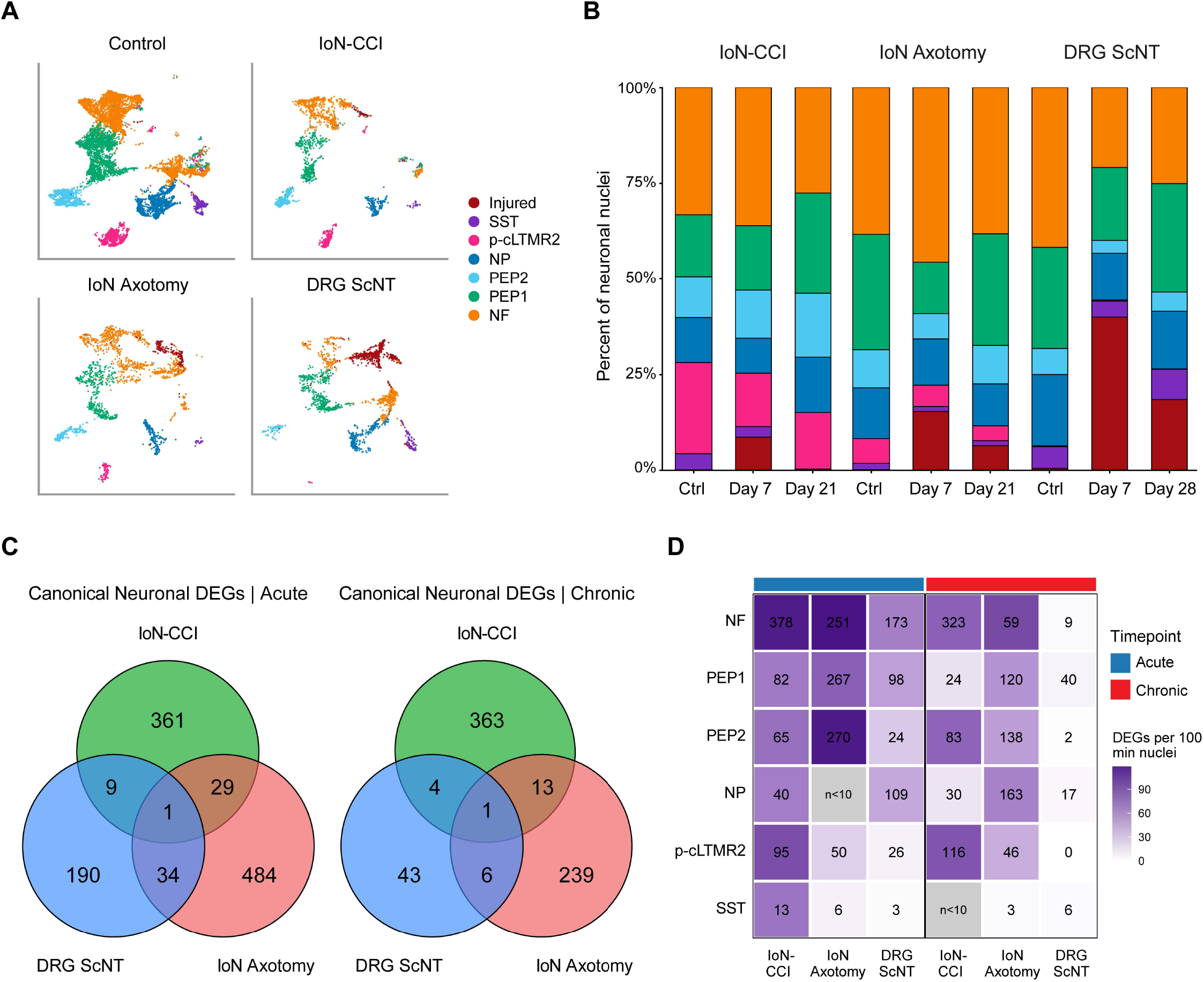
Cross-model comparison reveals distinct neuronal remodeling after constriction and transection injuries. **(A)** Integrated neuronal UMAP comparing the IoN-CCI dataset with published infraorbital nerve axotomy and dorsal root ganglion sciatic nerve transection (DRG ScNT) datasets. Neurons are colored by harmonized subtype annotations, including NF, NP, PEP1, PEP2, SST, p-cLTMR2, and injured-state neurons. **(B)** Relative neuronal subtype composition across models and timepoints. Bars show the percentage of recovered neuronal nuclei assigned to each subtype within each model and timepoint. Acute injury corresponds to Day 7 in all models; chronic injury corresponds to Day 21 for IoN-CCI and IoN axotomy and Day 28 for DRG ScNT. **(C)** Venn diagrams comparing pooled canonical-neuron differentially expressed genes (DEGs) across IoN-CCI, IoN axotomy, and DRG ScNT during acute and chronic injury phases. Canonical-neuron analyses included NF, PEP1, PEP2, NP, p-cLTMR2, and SST neurons and excluded injured-state neurons. DEGs were pooled across canonical subtypes within each model and phase. DEGs were defined as adjusted P < 0.05 and |log2 fold change| ≥ 0.5. **(D)** Heatmap of cell-number-normalized DEG burden across canonical neuronal subtypes, injury models, and injury phases. Injured-state neurons were excluded. DEGs were defined using the same threshold as in C (adjusted P < 0.05 and |log2 fold change| ≥ 0.5). Within each subtype, model, and injury phase, DEG burden was normalized to the smaller of the injury and control groups and expressed as the number of significant DEGs per 100 recovered nuclei. Tile color indicates cell-number-normalized DEG burden, and numbers within tiles indicate raw DEG counts. Comparisons with fewer than 10 recovered nuclei in either condition were excluded and are shown in gray. Acute injury corresponds to Day 7 for all models; chronic injury corresponds to Day 21 for TG IoN-CCI and TG axotomy and Day 28 for DRG ScNT.

### Cross-model comparison reveals distinct neuronal remodeling after constriction and transection injuries

We integrated IoN-CCI with published TG axotomy and DRG sciatic nerve transection datasets (Fig. 3A) (Nguyen et al. 2019; Renthal et al. 2020). Integrated analysis showed shared sensory neuron classes and injured-state neurons across datasets, with consistent subtype marker expression (Appendix Fig. 4). Subtype composition showed that IoN-CCI retained multiple canonical sensory neuron classes at Day 7 and Day 21, whereas axonal-severing models showed greater injured-state representation (Fig. 3B). Comparison of DEGs among neurons retaining canonical subtype identities showed that most differed across injury models at both acute and chronic stages, with notably limited overlap observed at the chronic stage (Fig. 3C). IoN-CCI-specific chronic DEGs included genes associated with synaptic organization, receptor signaling, axonal remodeling, and excitability, including *Nrxn1*, *Cntnap2*, *Dlg2*, *Grm7*, *Gabrb3*, *Hcn1*, *Kcnip4*, *Fgf13*, *Kcnmb2*, *Tmem45b*, and *Rgs8*. Several shared chronic DEGs also differed in increase/decrease directions across models, including *Dpp6*, *Rbfox1*, *Rbfox2*, *Prph*, and *Nefl*, indicating that overlap by gene identity did not necessarily reflect equivalent transcriptional regulation.

**Figure 4.**
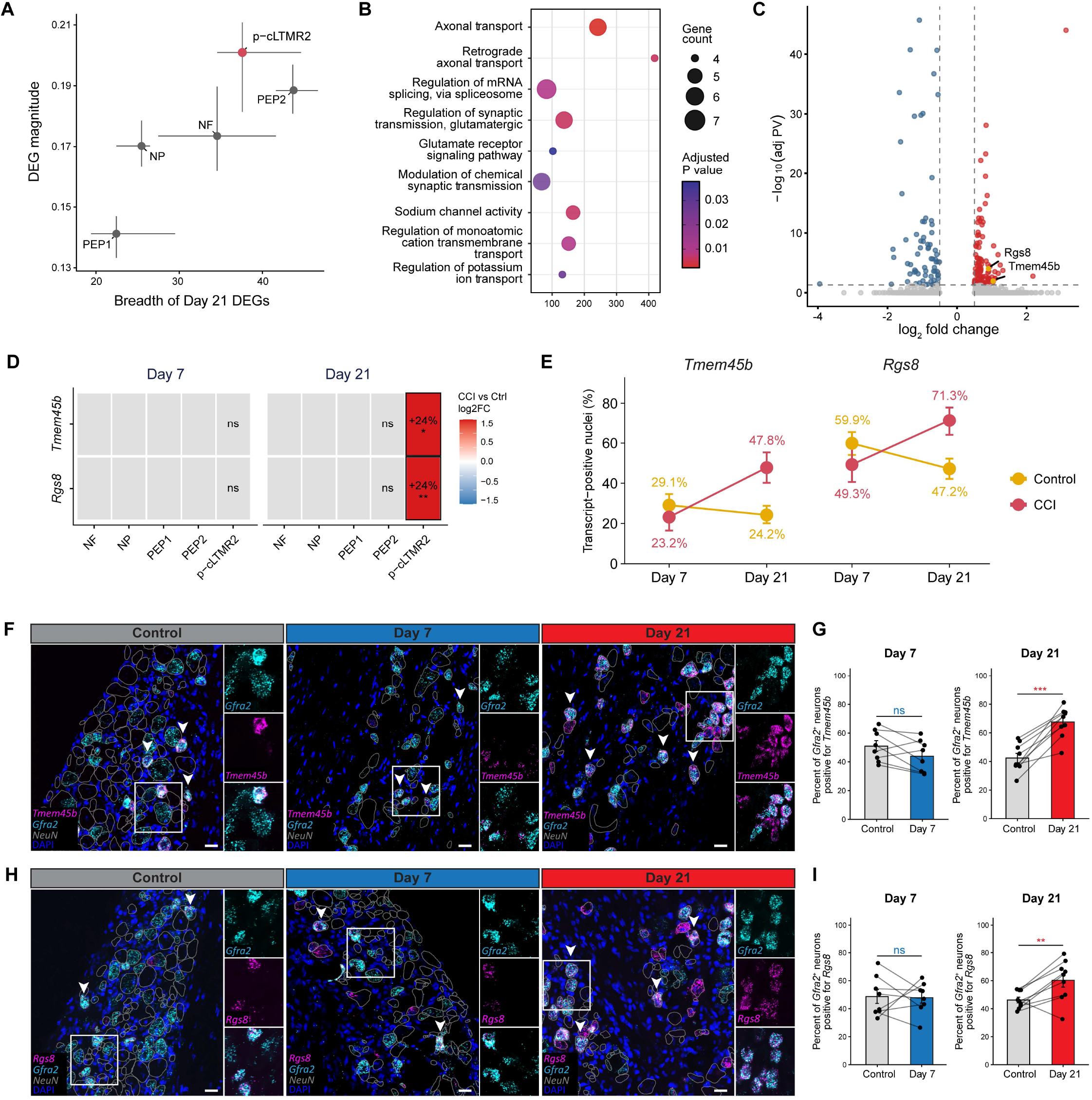
Chronic p-cLTMR2 remodeling identifies *Tmem45b* and *Rgs8* as delayed injury-induced transcripts after IoN-CCI. **(A)** Breadth and magnitude of Day 21 CCI-regulated DEGs across canonical neuronal subtypes. For each subtype, 100 nuclei per condition were randomly downsampled across 25 iterations. DEG breadth represents the number of significant DEGs, and DEG magnitude represents the significance-weighted root-mean-square log₂ fold change. Points indicate median values, with horizontal and vertical error bars showing the interquartile range across downsampling iterations. p-cLTMR2 neurons showed the highest DEG magnitude and were among the subtypes with the greatest DEG breadth. **(B)** Dot plot showing representative significant GO terms enriched among Day 21 p-cLTMR2 DEGs (adjusted P < 0.05), selected to represent major nonredundant biological processes. Dot size indicates gene count, color indicates adjusted P value, and the x-axis shows the Enrichr combined score. **(C)** Volcano plot showing Day 21 p-cLTMR2 differential expression comparing CCI and control. Red and blue points indicate significantly upregulated and downregulated genes, respectively, and highlighted candidate genes include *Tmem45b* and *Rgs8*. Dashed lines indicate the log2 fold-change and adjusted P-value thresholds. **(D)** Heatmap showing average log2 fold change of *Tmem45b* and *Rgs8* across canonical neuronal subtypes at Day 7 and Day 21. Tile color indicates CCI-versus-control log2 fold change, and values indicate the CCI–control difference in the percentage of transcript-positive nuclei. Asterisks indicate significant differential expression and gray tiles indicate nonsignificant comparisons. **(E)** Percentage of *Tmem45b*- and *Rgs8*-positive p-cLTMR2 nuclei at Day 7 and Day 21. Nuclei were classified as transcript-positive when ≥1 RNA UMI was detected. Points indicate the percentage of transcript-positive nuclei and error bars indicate exact 95% binomial confidence intervals. Condition × timepoint interactions were assessed using binomial generalized linear models (*Tmem45b*, BH-adjusted *P* = 6.22 × 10⁻⁶; *Rgs8*, BH-adjusted *P* = 4.72 × 10⁻⁷). **(F–I)** RNAscope validation of delayed transcriptional changes in *Gfra2*-positive trigeminal ganglion neurons after IoN-CCI. **(F)** Representative multiplex RNAscope images showing *Tmem45b* expression (magenta) in *Gfra2*-positive neurons (cyan) from control, Day 7 IoN-CCI, and Day 21 IoN-CCI trigeminal ganglia. *Gfra2* marks p-cLTMR2-enriched neurons. NeuN-guided cell-body outlines are shown in gray for visualization only and were not used for quantitative analysis; RNAscope quantification was performed using the automated NeuN-gated, DAPI-based ROI pipeline described in the Appendix Methods. Arrowheads indicate representative *Gfra2*-positive neurons, and boxed regions are shown as enlarged channel-separated insets. **(G)** Quantification of the percentage of *Gfra2*-positive neurons positive for *Tmem45b* at Day 7 and Day 21. *Tmem45b* expression was not significantly altered at Day 7 but was significantly increased at Day 21 after IoN-CCI. **(H)** Representative multiplex RNAscope images showing *Rgs8* expression (magenta) in *Gfra2*-positive neurons (cyan) from control, Day 7 IoN-CCI, and Day 21 IoN-CCI trigeminal ganglia. Arrowheads indicate representative *Gfra2*-positive neurons, and boxed regions are shown as enlarged channel-separated insets. **(I)** Quantification of the percentage of *Gfra2*-positive neurons positive for *Rgs8* at Day 7 and Day 21. *Rgs8* expression was not significantly altered at Day 7 but was significantly increased at Day 21 after IoN-CCI. For RNAscope quantification in G and I, n = 8 mice at Day 7 and n = 9 mice at Day 21. Each mouse contributed matched contralateral sham-control and ipsilateral IoN-CCI trigeminal ganglia and represents one biological replicate. For each mouse, one section from the contralateral sham-control TG and one section from the ipsilateral IoN-CCI TG were imaged and analyzed. The entire TG cross-section was included in the quantitative analysis without selection of specific fields or subregions, yielding one whole-TG-section value per condition for each mouse. Data are shown as mean ± SEM, with paired points representing matched control and CCI samples. Matched control and IoN-CCI values within each timepoint were compared using paired two-tailed t tests. Scale bars, 20 µm. Images were acquired at 40× magnification. ns, not significant; **P < 0.01; ***P < 0.001.

We next compared cell-number-normalized DEG burden across subtypes, models, and timepoints (Fig. 3D). SST neurons are shown for completeness but were excluded from interpretation because of sparse recovery and low DEG counts across comparisons. This analysis showed that transcriptional remodeling was not uniform across injury paradigms. During the acute phase, TG axotomy showed strong normalized DEG burden in PEP1, PEP2, and NF neurons, whereas IoN-CCI showed acute remodeling across NF, PEP1, PEP2, and p-cLTMR2 neurons, with comparatively lower burden in NP neurons. At the chronic stage, IoN-CCI showed the strongest burden in NF and p-cLTMR2 neurons, with additional remodeling in PEP2 neurons; TG axotomy showed broader remodeling across NF, NP, PEP1, and PEP2 neurons, whereas DRG ScNT showed comparatively limited remodeling. These patterns indicate that chronic IoN-CCI remodeling differs substantially from axonal-severing injury models and preferentially involves NF, p-cLTMR2, and PEP2 sensory neuron populations. Additional threshold-sensitivity, regulatory, and equal-cell subsampling analyses supported the broader model-specific remodeling patterns (Appendix Fig. 5).

**Figure 5.**
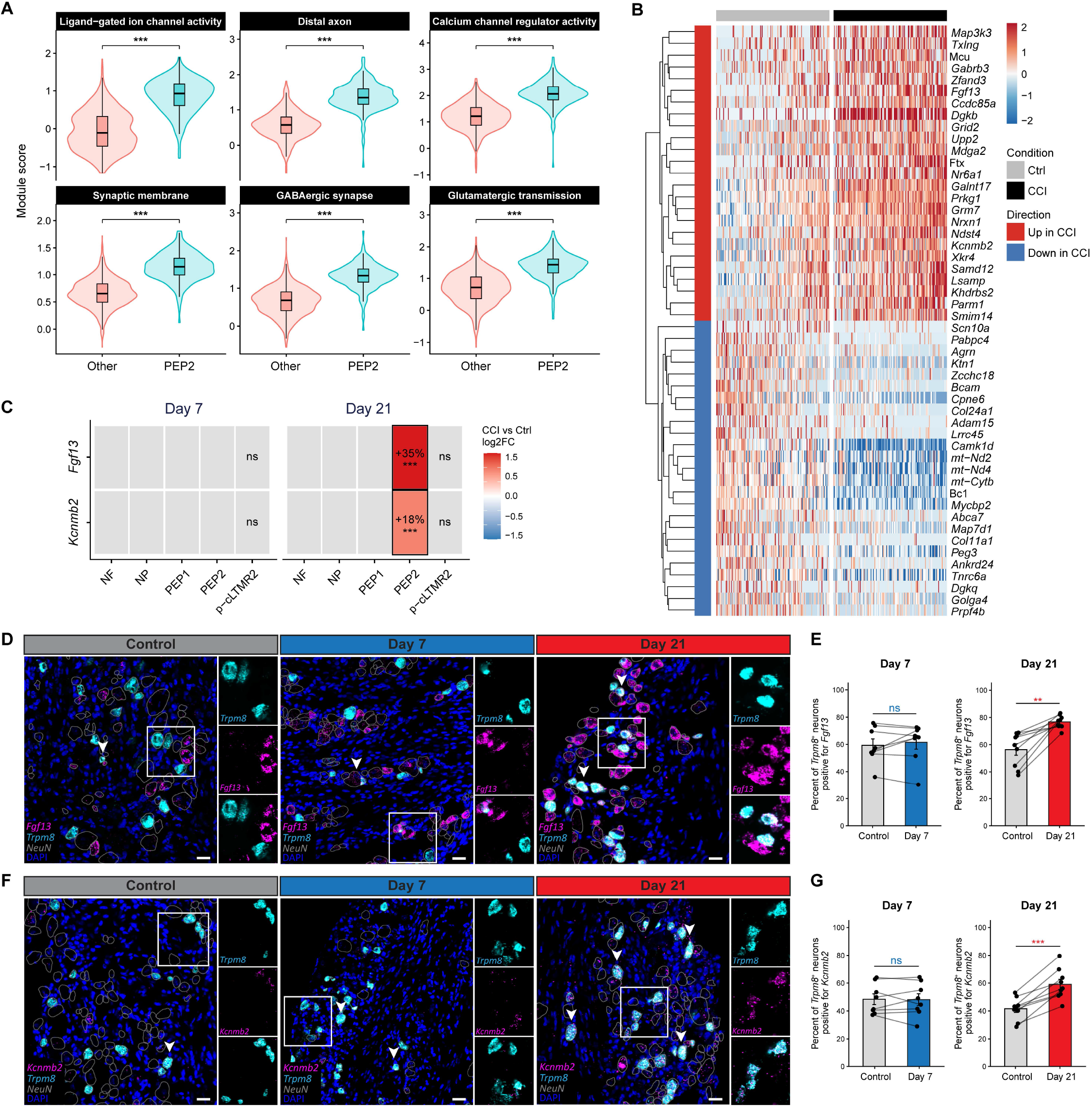
Chronic IoN-CCI induces PEP2-associated upregulation of excitability- and synaptic remodeling genes. **(A)** Violin plots showing per-nucleus module scores for selected functional gene programs in Day 21 CCI PEP2 neurons compared with pooled non-PEP2 Day 21 CCI canonical neuronal subtypes. Module scores were compared using two-sided Wilcoxon rank-sum tests with Benjamini–Hochberg correction across the six displayed terms. Boxplots indicate median and interquartile range. ***adjusted P < 0.001. **(B)** Heatmap showing scaled expression of representative differentially expressed genes in PEP2 neurons from control and CCI samples. Columns represent individual nuclei and rows represent genes. The top annotation indicates condition, and the side annotation indicates genes upregulated or downregulated in CCI. **(C)** Heatmap showing CCI-versus-control differential expression of *Fgf13* and *Kcnmb2* across canonical neuronal subtypes at Day 7 and Day 21 after IoN-CCI. Tile color indicates log2 fold change, and values indicate the CCI–control difference in the percentage of nuclei with detectable expression. Differential expression was tested within each subtype and timepoint using MAST. Significant changes were defined as adjusted P < 0.05 and |log2 fold change| > 0.25; asterisks indicate the adjusted P-value level for comparisons meeting these criteria and gray tiles indicate nonsignificant comparisons. *adjusted P < 0.05; **adjusted P < 0.01; ***adjusted P < 0.001. **(D–G)** RNAscope validation of delayed transcriptional changes in *Trpm8*-positive trigeminal ganglion neurons after IoN-CCI. **(D)** Representative RNAscope images showing Fgf13 expression (magenta) in *Trpm8*-positive neurons (cyan) from control, Day 7 IoN-CCI, and Day 21 IoN-CCI trigeminal ganglia. Trpm8 marks PEP2-enriched neurons. NeuN-guided cell-body outlines are shown in gray for visualization only and were not used for quantitative analysis; RNAscope quantification was performed using the automated NeuN-gated, DAPI-based ROI pipeline described in the Appendix Methods. Arrowheads indicate representative *Trpm8*-positive neurons, and boxed regions are shown as enlarged channel-separated insets. **(E)** Quantification of the percentage of *Trpm8*-positive neurons positive for *Fgf13* at Day 7 and Day 21. *Fgf13* expression was not significantly altered at Day 7 but was significantly increased at Day 21 after IoN-CCI. **(F)** Representative RNAscope images showing *Kcnmb2* expression (magenta) in *Trpm8*-positive neurons (cyan) from control, Day 7 IoN-CCI, and Day 21 IoN-CCI trigeminal ganglia. Arrowheads indicate representative *Trpm8*-positive neurons, and boxed regions are shown as enlarged channel-separated insets. **(G)** Quantification of the percentage of *Trpm8*-positive neurons positive for *Kcnmb2* at Day 7 and Day 21. *Kcnmb2* expression was not significantly altered at Day 7 but was significantly increased at Day 21 after IoN-CCI. For RNAscope quantification in E and G, n = 8 mice at Day 7 and n = 9 mice at Day 21. Each mouse contributed matched contralateral sham-control and ipsilateral IoN-CCI trigeminal ganglia and represents one biological replicate. For each mouse, one section from the contralateral sham-control TG and one section from the ipsilateral IoN-CCI TG were imaged and analyzed. The entire TG cross-section was included in the quantitative analysis without selection of specific fields or subregions, yielding one whole-TG-section value per condition for each mouse. Data are shown as mean ± SEM, with paired points representing matched control and CCI samples. Matched control and IoN-CCI values within each timepoint were compared using paired two-tailed t tests. Scale bars, 20 µm. Images were acquired at 40× magnification. ns, not significant; **P < 0.01; ***P < 0.001.

### Chronic p-cLTMR2 remodeling identifies Tmem45b and Rgs8 as delayed injury-induced transcripts

At Day 21, p-cLTMR2 neurons exhibited the greatest DEG magnitude and ranked among the subtypes with the highest DEG breadth, alongside PEP2 neurons (Fig. 4A). Functional enrichment of Day 21 p-cLTMR2 DEGs highlighted axonal transport, mRNA splicing, synaptic signaling, and ion-channel and ion-transport processes (Fig. 4B). Among these chronic DEGs, *Tmem45b* and *Rgs8* were prioritized for validation (Fig. 4C). Comparison across neuronal subtypes and timepoints showed that significant induction of both genes was confined to p-cLTMR2 neurons at Day 21 among the analyzed canonical subtypes (Fig. 4D,E). *Tmem45b*-positive nuclei increased from 24.2% in control to 47.8% after CCI, and *Rgs8*-positive nuclei increased from 47.2% to 71.3%, whereas neither gene showed significant induction at Day 7. Nucleus-level binomial models supported significant condition-by-timepoint interactions for *Tmem45b* (BH-adjusted *P* = 6.22 × 10⁻⁶) and *Rgs8* (BH-adjusted *P* = 4.72 × 10⁻⁷) (Fig. 4E). Equal-nucleus subsampling confirmed that the observed Day 21 increases in *Tmem45b* and *Rgs8* detection were not due to unequal nucleus recovery (Appendix Fig. 6).

Multiplex RNAscope was used to validate *Tmem45b* and *Rgs8* expression in *Gfra2*-positive p-cLTMR2-enriched neurons. RNAscope confirmed increased *Tmem45b* expression in *Gfra2*-positive neurons at Day 21 CCI, with no significant change at Day 7 (Fig. 4F,G). *Rgs8* showed the same delayed expression pattern, with increased expression at Day 21 by imaging and quantification (Fig. 4H,I). These findings support delayed remodeling of canonical mechanosensory neurons at the chronic stage after IoN-CCI.

### Chronic PEP2 remodeling identifies Fgf13 and Kcnmb2 as delayed injury-induced transcripts

Within Day 21 CCI neurons, PEP2 neurons showed higher scores for ion-channel, axonal, and synaptic gene programs than other canonical neuronal subtypes (Fig. 5A). We next visualized the top Day 21 PEP2-regulated DEGs using a z-scored heatmap (Fig. 5B). CCI-associated increases included genes involved in ion-channel regulation, synaptic signaling, and neuronal connectivity, including *Fgf13, Kcnmb2, Dgkb, Grm7, Gabrb3, Prkg1, Nrxn1, Mdga2,* and *Grid2*. Downregulated genes included transcripts associated with sodium-channel function, calcium signaling, axonal regulation, and mitochondrial function, including *Scn10a, Camk1d, Mycbp2, mt-Nd2, mt-Nd4,* and *mt-Cytb*. These changes indicated chronic remodeling of excitability-, synaptic-, and metabolism-associated gene programs in PEP2 neurons. Functional enrichment and candidate prioritization supported the chronic PEP2 transcriptional response and the selection of *Fgf13* and *Kcnmb2* for RNAscope validation (Appendix Fig. 7).

To assess temporal and subtype specificity, we compared *Fgf13* and *Kcnmb2* regulation across canonical neuronal subtypes at Day 7 and Day 21 (Fig. 5C). At Day 7, neither gene showed strong subtype-selective induction after CCI. By Day 21, however, *Fgf13* and *Kcnmb2* were significantly upregulated by nucleus-level snRNA-seq analysis and validated at the mouse level by RNAscope. Representative images showed increased *Fgf13* signal in *Trpm8*-positive neurons at Day 21 CCI compared with control and Day 7 CCI (Fig. 5D). Quantification showed a significant increase in the percentage of *Trpm8*-positive neurons expressing *Fgf13* at Day 21 (Fig. 5E). *Kcnmb2* showed a similar pattern, with increased expression in *Trpm8*-positive neurons at Day 21 CCI and significant Day 21 elevation by quantification (Fig. 5F and G). These findings show delayed remodeling in *Trpm8*-positive PEP2 neurons, including induction of excitability-associated genes *Fgf13* and *Kcnmb2*.

## Discussion

This study defines temporal and subtype-specific remodeling of trigeminal sensory neurons after IoN-CCI, a model of orofacial neuropathic pain. IoN-CCI produced an early injured-neuron response at Day 7 and delayed remodeling of canonical neurons by Day 21. The Day 7 response was enriched for injury- and regeneration-associated genes, including *Atf3* and *Sox11*, whereas the Day 21 response was marked by synaptic and neurotransmitter signaling, axonal, ion channel, and structural remodeling programs. These data indicate that major sensory neuron subtype identities remain largely recognizable while specific canonical subtypes undergo delayed transcriptional remodeling.

Axotomy and transection-type models induce shared regeneration-associated transcriptional programs across injured sensory neurons, including *Atf3*- and *Sox11*-associated responses accompanied by suppression of subtype-specific identity (Nguyen et al. 2019; Renthal et al. 2020). In contrast, IoN-CCI produced a small injury-associated neuronal population while major sensory-neuron subtype identities remained recognizable. By Day 21, these canonical populations still exhibited changes in gene expression. This pattern is consistent with partial nerve injury models, in which injured and canonical afferents coexist within the same sensory ganglion, and intact afferents can undergo plasticity that contributes to stimulus-evoked neuropathic pain (Djouhri et al. 2012; Obata et al. 2003; Tran and Crawford 2020). Accordingly, the IoN-CCI response was not a persistent version of the Day 7 injury state. Most DEGs shared between Day 7 and Day 21 changed direction over time, and many chronic-stage DEGs were Day 21-specific. These findings indicate that Day 21 represents a later remodeling phase in canonical neurons that is distinct from the acute injury response. RNA-processing and splicing pathways shifted from enrichment among CCI-upregulated genes at Day 7 to CCI-downregulated genes at Day 21, indicating temporal reorganization of these programs. Consistent with this possibility, peripheral nerve injury has been shown to alter alternative splicing in sensory neurons, including *Rbfox1*-dependent splicing programs associated with neuropathic pain (He et al. 2024). Day 21-specific transcripts included ion-channel and excitability-associated genes with established links to sensory neuron function and pain, including *Scn9a*, *Cacna2d1*, *Asic3*, *Trpm3*, and *Hcn1*. These changes suggest that chronic-stage orofacial nerve injury may involve delayed plasticity in canonical neurons.

Cross-model comparison further showed distinct patterns of sensory-neuron remodeling across injury paradigms and ganglia. IoN-CCI shared few chronic differentially expressed genes with TG axotomy and DRG sciatic nerve transection. Chronic IoN-CCI preferentially affected pathways involved in neuronal signaling and excitability. These model-specific patterns support the use of direct trigeminal constriction injury models, because DRG transection mechanisms may not fully represent orofacial neuropathic pain programs. The TG contains sensory neurons that innervate facial and oral tissues, and its injury response appears distinct from both TG axotomy and DRG transection models (Megat et al. 2019; Nguyen et al. 2017; Yang et al. 2022). Nevertheless, these cross-model comparisons should be interpreted somewhat cautiously because the datasets differed in injury modality, ganglion, sequencing preparation, study source, and chronic timepoint.

A major finding of our study is the delayed remodeling of p-cLTMR2 neurons after IoN-CCI. C-low threshold mechanoreceptors normally encode innocuous touch, but injury can convert low-threshold mechanosensory pathways into mechanical hypersensitivity (Seal et al. 2009). Day 21 p-cLTMR2 neurons showed enrichment of transcriptional programs related to axonal transport, mRNA splicing, synaptic and glutamatergic signaling, and ion-channel and ion-transport regulation. *Tmem45b* and *Rgs8* were selectively upregulated in p-cLTMR2 neurons at Day 21 and validated in *Gfra2*-positive neurons by RNAscope. *Tmem45b* contributes to mechanical hypersensitivity following inflammation or surgical incision in mice (Tanioku et al. 2022). Our findings identify *Tmem45b* as a candidate transcript associated with chronic remodeling in trigeminal p-cLTMR2 neurons after IoN-CCI. Increased *Rgs8* expression suggests a potential change in G-protein signaling regulation (Saitoh et al. 1997; Senese et al. 2020), although its functional consequences in trigeminal mechanosensory neurons remain unknown. Functional studies are needed to determine whether either transcript contributes directly to mechanical hypersensitivity after orofacial nerve injury.

PEP2 neurons also showed a delayed chronic response. These neurons are enriched for *Trpm*8, a cold-sensing channel expressed by subsets of TG neurons and implicated in cold allodynia and trigeminal pain (Gualdani et al. 2021; MacDonald et al. 2021). Recent IoN-CCI work links TRPM8-lineage neurons and Kv1.1/Kv1.2-dependent potassium currents to orofacial cold allodynia (Pina et al. 2024). At Day 21, PEP2 neurons showed increased module scores for ligand-gated ion channel activity, calcium channel regulator activity, distal axon, synaptic membrane, GABAergic synapse, and glutamatergic transmission. *Fgf13* and *Kcnmb2* were selectively upregulated in PEP2 neurons at Day 21 and validated in *Trpm8*-positive neurons by RNAscope. FGF13 has established roles in sodium channel regulation, axonal organization, and nociceptor excitability (Singh et al. 2025; Wang et al. 2021). *Kcnmb2* may link the PEP2 response to potassium-channel regulation and sensory neuron excitability (Chen et al. 2009; Zarei et al. 2007). These findings identify *Fgf13* and *Kcnmb2* as chronic PEP2-associated candidates after IoN-CCI.

In summary, IoN-CCI induces staged trigeminal neuron remodeling, progressing from an acute injured-neuron state to chronic subtype-specific remodeling in canonical sensory neurons, thereby identifying candidate molecular programs that may guide future mechanistic studies of chronic pain development after orofacial nerve injury.

## Acknowledgments

The authors thank the National Institute of Dental and Craniofacial Research/National Institutes of Health animal facility staff for technical support. The authors thank Andrew Doyle, Hwei Ling, Marit Aure, and the NIDCR Imaging Core (ZIC DE000750-01) for assistance with confocal imaging and image acquisition. The authors also thank Elina Stregevsky and George McGrady of the NIDCR Combined Technical Research Core (ZIC DE000729-09) for assistance with nuclei sorting, and Changyu Zheng of the NIDCR Genomics and Computational Biology Core (ZIC DC000086) for single-nucleus RNA-sequencing library preparation and PCR. We thank Minh Q. Nguyen for sharing the nuclei-isolation/single-nucleus RNA-sequencing protocol. We also thank Advait Limaye for technical assistance with mouse procurement, anesthesia and analgesia administration, and perioperative animal care needed for the IoN-CCI experiments. The authors also thank members of the Yamada and Kulkarni laboratories for helpful discussions and feedback during the development of this study. Figure illustrations were created in part with BioRender.com. AI was used to assist with literature searches and the identification of publications during manuscript preparation.

The contributions of the NIH authors were made as part of their official duties as NIH federal employees, are in compliance with agency policy requirements, and are considered Works of the United States Government. However, the findings and conclusions presented in this paper are those of the authors and do not necessarily reflect the views of the NIH or the U.S. Department of Health and Human Services.

## Author Contributions

L.F. Cook contributed to conception, design, methodology, data acquisition, analysis, and interpretation, drafted the manuscript, and critically revised the manuscript; M. Hu contributed to conception, design, methodology, data acquisition, drafted portions of the manuscript, and critically revised the manuscript; D. Izquierdo contributed to methodology, single-nucleus RNA-sequencing library preparation, and initial data analysis and code development; S. Yun contributed to data analysis and interpretation and critically revised the manuscript; B.E. Hall contributed to data analysis and interpretation and critically revised the manuscript; K.M. Yamada contributed to conception, design, supervision, and data interpretation and critically revised the manuscript; A.B. Kulkarni contributed to conception, design, supervision, and data interpretation and critically revised the manuscript. All authors gave their final approval and agree to be accountable for all aspects of the work.

## Statements and Declarations

### Ethical considerations

All animal procedures were approved by the National Institute of Dental and Craniofacial Research (NIDCR) Institutional Animal Care and Use Committee (Animal Study Protocols 24-1161 and 19-894) and conducted in accordance with National Institutes of Health (NIH) guidelines.

### Consent to participate

Not applicable. This study involved laboratory animals and did not include human participants.

### Consent for publication

Not applicable.

### Data and Materials Availability

Single-nucleus RNA-sequencing data generated in this study have been deposited in the Gene Expression Omnibus (GEO) and will be made publicly available upon peer-reviewed publication. Publicly available datasets reanalyzed in this study include Nguyen et al. 2019/GSE131272 and Renthal et al. 2020/GSE154659. Processed data, analysis code, and image-analysis scripts will be made available through a public repository upon peer-reviewed publication.

## References

Araki T, Wakatsuki S. 2019. Regulation of neuronal/axonal degeneration by znrf1 ubiquitin ligase. Neurosci Res. 139(21–25. https://www.ncbi.nlm.nih.gov/pubmed/30118738. doi:10.1016/j.neures.2018.07.008.

Benoliel R et al. 2019. The iasp classification of chronic pain for icd-11: Chronic secondary headache or orofacial pain. Pain. 160(1):60–68. https://www.ncbi.nlm.nih.gov/pubmed/30586072. doi:10.1097/j.pain.0000000000001435.

Bhuiyan SA et al. 2024. Harmonized cross-species cell atlases of trigeminal and dorsal root ganglia. Sci Adv. 10(25):eadj9173. https://www.ncbi.nlm.nih.gov/pubmed/38905344. doi:10.1126/sciadv.adj9173.

Boccella S et al. 2020. The modulation of pain by metabotropic glutamate receptors 7 and 8 in the dorsal striatum. Curr Neuropharmacol. 18(1):34–50. https://www.ncbi.nlm.nih.gov/pubmed/31210112. doi:10.2174/1570159X17666190618121859.

Chen SR, Cai YǪ, Pan HL. 2009. Plasticity and emerging role of bkca channels in nociceptive control in neuropathic pain. J Neurochem. 110(1):352–362. https://www.ncbi.nlm.nih.gov/pubmed/19457113. doi:10.1111/j.1471-4159.2009.06138.x.

Cui X et al. 2023. Transcriptome-wide analysis of trigeminal ganglion and subnucleus caudalis in a mouse model of chronic constriction injury-induced trigeminal neuralgia. Front Pharmacol. 14(1230633. https://www.ncbi.nlm.nih.gov/pubmed/37841912. doi:10.3389/fphar.2023.1230633.

Dawes JM et al. 2018. Immune or genetic-mediated disruption of caspr2 causes pain hypersensitivity due to enhanced primary afferent excitability. Neuron. 97(4):806–822 e810. https://www.ncbi.nlm.nih.gov/pubmed/29429934. doi:10.1016/j.neuron.2018.01.033.

Deseure KR, Hans GH. 2024. Chronic constriction injury of the distal infraorbital nerve (dion-cci) in mice to study trigeminal neuropathic pain. J Vis Exp. 205). https://www.ncbi.nlm.nih.gov/pubmed/38526120. doi:10.3791/66420.

Ding W et al. 2017. An improved rodent model of trigeminal neuropathic pain by unilateral chronic constriction injury of distal infraorbital nerve. J Pain. 18(8):899–907. https://www.ncbi.nlm.nih.gov/pubmed/28238950. doi:10.1016/j.jpain.2017.02.427.

Djouhri L, Fang X, Koutsikou S, Lawson SN. 2012. Partial nerve injury induces electrophysiological changes in conducting (uninjured) nociceptive and nonnociceptive drg neurons: Possible relationships to aspects of peripheral neuropathic pain and paresthesias. Pain. 153(9):1824–1836. https://www.ncbi.nlm.nih.gov/pubmed/22721911. doi:10.1016/j.pain.2012.04.019.

Fang Z et al. 2026. Lncrna 4930544m13rik-201 regulates cacna2d1 expression via interacting with hnrnpa2b1 to promote neuropathic pain following nerve injury. Brain Res Bull. 238(111838. https://www.ncbi.nlm.nih.gov/pubmed/41864512. doi:10.1016/j.brainresbull.2026.111838.

Gambeta E, Chichorro JG, Zamponi GW. 2020. Trigeminal neuralgia: An overview from pathophysiology to pharmacological treatments. Mol Pain. 16(1744806920901890. https://www.ncbi.nlm.nih.gov/pubmed/31908187. doi:10.1177/1744806920901890.

Ghosh K, Huang Y, Chen SR, Pan HL. 2024. Nerve injury augments cacna2d1 transcription via ck2-mediated phosphorylation of the histone deacetylase hdac2 in dorsal root ganglia. J Biol Chem. 300(11):107848. https://www.ncbi.nlm.nih.gov/pubmed/39357831. doi:10.1016/j.jbc.2024.107848.

Gualdani R et al. 2021. Trigeminal neuralgia trpm8 mutation: Enhanced activation, basal [ca(2+)](i) and menthol response. Neurol Genet. 7(1):e550. https://www.ncbi.nlm.nih.gov/pubmed/33977138. doi:10.1212/NXG.0000000000000550.

He L et al. 2024. Rbfox1 regulates alternative splicing of nrcam in primary sensory neurons to mediate peripheral nerve injury-induced neuropathic pain. Neurotherapeutics. 21(1):e00309. https://www.ncbi.nlm.nih.gov/pubmed/38241164. doi:10.1016/j.neurot.2023.e00309.

International classification of orofacial pain, 1st edition (icop). 2020. Cephalalgia. 40(2):129–221. https://www.ncbi.nlm.nih.gov/pubmed/32103673. doi:10.1177/0333102419893823.

Khan S, Tao F. 2025. Mechanisms for orofacial pain: Roles of immunomodulation, metabolic reprogramming, oxidative stress and epigenetic regulation. Biomedicines. 13(2). https://www.ncbi.nlm.nih.gov/pubmed/40002847. doi:10.3390/biomedicines13020434.

Korczeniewska OA et al. 2018. Differential gene expression in trigeminal ganglia of male and female rats following chronic constriction of the infraorbital nerve. Eur J Pain. 22(5):875–888. https://www.ncbi.nlm.nih.gov/pubmed/29350446. doi:10.1002/ejp.1174.

Korczeniewska OA, Kohli D, Benoliel R, Baddireddy SM, Eliav E. 2022. Pathophysiology of post-traumatic trigeminal neuropathic pain. Biomolecules. 12(12). https://www.ncbi.nlm.nih.gov/pubmed/36551181. doi:10.3390/biom12121753.

Kung CC et al. 2020. Deletion of acid-sensing ion channel 3 relieves the late phase of neuropathic pain by preventing neuron degeneration and promoting neuron repair. Cells. 9(11). https://www.ncbi.nlm.nih.gov/pubmed/33114619. doi:10.3390/cells9112355.

Li WG, Xu TL. 2011. Asic3 channels in multimodal sensory perception. ACS Chem Neurosci. 2(1):26–37. https://www.ncbi.nlm.nih.gov/pubmed/22778854. doi:10.1021/cn100094b.

Lovatt D et al. 2022. Scrna-seq generates a molecular map of emerging cell subtypes after sciatic nerve injury in rats. Commun Biol. 5(1):1105. https://www.ncbi.nlm.nih.gov/pubmed/36261573. doi:10.1038/s42003-022-03970-0.

MacDonald DI et al. 2021. Silent cold-sensing neurons contribute to cold allodynia in neuropathic pain. Brain. 144(6):1711–1726. https://www.ncbi.nlm.nih.gov/pubmed/33693512. doi:10.1093/brain/awab086.

Megat S et al. 2019. Differences between dorsal root and trigeminal ganglion nociceptors in mice revealed by translational profiling. J Neurosci. 39(35):6829–6847. https://www.ncbi.nlm.nih.gov/pubmed/31253755. doi:10.1523/JNEUROSCI.2663-18.2019.

Nguyen MQ, Le Pichon CE, Ryba N. 2019. Stereotyped transcriptomic transformation of somatosensory neurons in response to injury. Elife. 8(https://www.ncbi.nlm.nih.gov/pubmed/31592768. doi:10.7554/eLife.49679.

Nguyen MQ, Wu Y, Bonilla LS, von Buchholtz LJ, Ryba NJP. 2017. Diversity amongst trigeminal neurons revealed by high throughput single cell sequencing. PLoS One. 12(9):e0185543. https://www.ncbi.nlm.nih.gov/pubmed/28957441. doi:10.1371/journal.pone.0185543.

Obata K et al. 2003. Contribution of injured and uninjured dorsal root ganglion neurons to pain behavior and the changes in gene expression following chronic constriction injury of the sciatic nerve in rats. Pain. 101(1-2):65–77. https://www.ncbi.nlm.nih.gov/pubmed/12507701. doi:10.1016/s0304-3959(02)00296-8.

Pina R et al. 2024. A functional unbalance of trpm8 and kv1 channels underlies orofacial cold allodynia induced by peripheral nerve damage. Front Pharmacol. 15(1484387. https://www.ncbi.nlm.nih.gov/pubmed/39703391. doi:10.3389/fphar.2024.1484387.

Renthal W et al. 2020. Transcriptional reprogramming of distinct peripheral sensory neuron subtypes after axonal injury. Neuron. 108(1):128–144 e129. https://www.ncbi.nlm.nih.gov/pubmed/32810432. doi:10.1016/j.neuron.2020.07.026.

Romero-Reyes M et al. 2023. Pharmacological management of orofacial pain. Drugs. 83(14):1269–1292. https://www.ncbi.nlm.nih.gov/pubmed/37632671. doi:10.1007/s40265-023-01927-z.

Saitoh O, Kubo Y, Miyatani Y, Asano T, Nakata H. 1997. Rgs8 accelerates g-protein-mediated modulation of k+ currents. Nature. 390(6659):525–529. https://www.ncbi.nlm.nih.gov/pubmed/9394004. doi:10.1038/37385.

Seal RP et al. 2009. Injury-induced mechanical hypersensitivity requires c-low threshold mechanoreceptors. Nature. 462(7273):651–655. https://www.ncbi.nlm.nih.gov/pubmed/19915548. doi:10.1038/nature08505.

Seijffers R, Mills CD, Woolf CJ. 2007. Atf3 increases the intrinsic growth state of drg neurons to enhance peripheral nerve regeneration. J Neurosci. 27(30):7911–7920. https://www.ncbi.nlm.nih.gov/pubmed/17652582. doi:10.1523/JNEUROSCI.5313-06.2007.

Senese NB, Kandasamy R, Kochan KE, Traynor JR. 2020. Regulator of g-protein signaling (rgs) protein modulation of opioid receptor signaling as a potential target for pain management. Front Mol Neurosci. 13(5. https://www.ncbi.nlm.nih.gov/pubmed/32038168. doi:10.3389/fnmol.2020.00005.

Shinoda M et al. 2021. Orofacial neuropathic pain-basic research and their clinical relevancies. Front Mol Neurosci. 14(691396. https://www.ncbi.nlm.nih.gov/pubmed/34295221. doi:10.3389/fnmol.2021.691396.

Singh AK et al. 2025. Sensory neuron-expressed fgf13 controls nociceptive signaling in diabetic neuropathy models. J Clin Invest. 135(14). https://www.ncbi.nlm.nih.gov/pubmed/40662354. doi:10.1172/JCI183749.

Tanioku T et al. 2022. Tmem45b is essential for inflammation- and tissue injury-induced mechanical pain hypersensitivity. Proc Natl Acad Sci U S A. 119(45):e2121989119. https://www.ncbi.nlm.nih.gov/pubmed/36322717. doi:10.1073/pnas.2121989119.

Tran EL, Crawford LK. 2020. Revisiting pns plasticity: How uninjured sensory afferents promote neuropathic pain. Front Cell Neurosci. 14(612982. https://www.ncbi.nlm.nih.gov/pubmed/33362476. doi:10.3389/fncel.2020.612982.

Wakatsuki S, Saitoh F, Araki T. 2011. Znrf1 promotes wallerian degeneration by degrading akt to induce gsk3b-dependent crmp2 phosphorylation. Nat Cell Biol. 13(12):1415–1423. https://www.ncbi.nlm.nih.gov/pubmed/22057101. doi:10.1038/ncb2373.

Wang Q, et al. 2021. Fibroblast growth factor 13 stabilizes microtubules to promote na(+) channel function in nociceptive drg neurons and modulates inflammatory pain. J Adv Res. 31(97–111. https://www.ncbi.nlm.nih.gov/pubmed/34194835. doi:10.1016/j.jare.2020.12.009.

Xing H, Chen M, Ling J, Tan W, Gu JG. 2007. Trpm8 mechanism of cold allodynia after chronic nerve injury. J Neurosci. 27(50):13680–13690. https://www.ncbi.nlm.nih.gov/pubmed/18077679. doi:10.1523/JNEUROSCI.2203-07.2007.

Yang J et al. 2025. Restoration of mitochondrial function alleviates trigeminal neuropathic pain in mice. Free Radic Biol Med. 226(185–198. https://www.ncbi.nlm.nih.gov/pubmed/39528053. doi:10.1016/j.freeradbiomed.2024.11.011.

Yang L et al. 2022. Human and mouse trigeminal ganglia cell atlas implicates multiple cell types in migraine. Neuron. 110(11):1806–1821 e1808. https://www.ncbi.nlm.nih.gov/pubmed/35349784. doi:10.1016/j.neuron.2022.03.003.

Young A, Sahabi L, Noma N, Kalladka M, Yan Z. 2022. Temporomandibular disorders, neuropathic and idiopathic orofacial pain, and headaches: A literature review. Frontiers of Oral and Maxillofacial Medicine. 5(https://fomm.amegroups.org/article/view/62672.

Zarei MM et al. 2007. Endocytic trafficking signals in kcnmb2 regulate surface expression of a large conductance voltage and ca(2+)-activated k+ channel. Neuroscience. 147(1):80–89. https://www.ncbi.nlm.nih.gov/pubmed/17521822. doi:10.1016/j.neuroscience.2007.04.019.

Zhang C et al. 2022. Scrna-sequencing reveals subtype-specific transcriptomic perturbations in drg neurons of pirt(egfpf) mice in neuropathic pain condition. Elife. 11(https://www.ncbi.nlm.nih.gov/pubmed/36264609. doi:10.7554/eLife.76063.

