## Supplementary Appendix: Methods, Figures, and Tables for "Remodeling of Trigeminal Neuron Subtypes in Chronic Orofacial Pain"

### Appendix Materials and Methods

#### Mice

All animal procedures (ASP 24-1161, 19-894) were conducted in accordance with NIH guidelines and approved by the NIDCR Institutional Animal Care and Use Committee. This study is reported in accordance with ARRIVE 2.0 guidelines. Experiments were performed using C57BL/6J mice (The Jackson Laboratory, stock #000664), 8–10 weeks old and of both sexes. Mice were acclimated for at least 72 h before surgery and group-housed with up to five mice per cage under a 14 h light/10 h dark cycle at 22 ± 2 °C with enrichment and ad libitum access to food and water. A total of 42 mice entered the study: 24 for snRNA-seq and 18 for RNAscope validation. One mouse in the RNAscope cohort died following an injection and was excluded from analysis. No other animals were excluded. The snRNA-seq cohort included 16 mice at Day 7 (8 male, 8 female) and 8 at Day 21 (4 male, 4 female); the final RNAscope cohort included 8 mice at Day 7 (3 male, 5 female) and 9 at Day 21 (4 male, 5 female). RNAscope control and CCI comparisons were paired within individual mice. A minimum of eight mice per timepoint was targeted, although no formal a priori power calculation was performed. Mice were not randomized to timepoint, cohort, or surgical side; surgically treated mice received left-sided IoN-CCI and contralateral right-sided sham surgery. Investigators were not blinded to experimental condition, and the study protocol and analysis plan were not preregistered.

#### Infraorbital Chronic Constriction Injury (CCI) and Sham Surgery

IoN-CCI was performed under intraperitoneal ketamine (100 mg/kg) and xylazine (10 mg/kg) anesthesia, with anesthetic depth confirmed by absence of pedal withdrawal and palpebral reflexes. Ketoprofen (5 mg/kg, subcutaneous) was administered preoperatively and at approximately 24, 48, and 72 h thereafter. Mice were maintained on a warming surface set to 37 °C, and ophthalmic ointment was applied to both eyes. After preparation of the whisker-pad surgical field, an approximately 3-mm incision was made dorsal to the caudal whisker pad, and the main infraorbital nerve trunk was exposed by blunt dissection and isolated under magnification. The left infraorbital nerve underwent IoN-CCI and the contralateral right nerve underwent sham surgery. For IoN-CCI, a single circumferential 7-0 silk ligature was placed around the intact nerve and tightened until mild indentation was visible without blanching, complete occlusion, or transection; the ligature was left in place. Sham surgery involved identical nerve exposure and isolation without ligation. Wounds were closed with Vetbond tissue adhesive. Surgeries were performed aseptically, with instruments sterilized between animals according to approved institutional procedures. Mice recovered in prewarmed cages and were monitored until ambulatory and then daily for at least 10 days. Hydrogel and soft diet were provided for 7 days. Humane endpoints included poor body condition, emaciation, ruffled coat, or other signs of severe distress requiring veterinary evaluation; persistent body condition score ≤2/5 for 48 h was designated an endpoint.

**Experimental Design and Trigeminal Ganglia Collection for Single-Nucleus RNA-seq**

Mice were euthanized by gradual-fill $CO_2$ inhalation followed by cervical dislocation in accordance with approved protocols. Each mouse contributed an ipsilateral CCI TG and matched contralateral sham-control TG. For snRNA-seq, eight TG from the same eight mice were pooled per condition/library. Day 7 included two matched control/CCI library pairs (16 mice), whereas Day 21 included one matched pair (8 mice). TG were dissected bilaterally, rinsed in ice-cold PBS, frozen on dry ice, and stored at −80 °C until nuclei isolation. Nuclei isolation, FANS, and library preparation were performed on the same day for each batch. The final dataset comprised 3 control, 2 Day 7 CCI, and 1 Day 21 CCI libraries. Control libraries were pooled for atlas construction and selected visualizations, whereas timepoint-specific differential-expression analyses used the corresponding matched controls. Because TG were pooled before 10x capture, individual mouse and sex identities were not retained; snRNA-seq was therefore used for subtype-resolved discovery and candidate prioritization, with selected chronic-stage findings validated independently by paired mouse-level RNAscope.

**Nuclei Isolation and snRNA-seq Library Preparation:** Before nuclei isolation, reagents, homogenization equipment, and centrifuges were cooled to 4 °C. Nuclei isolation was adapted from the spinal cord protocol described by Alkaslasi et al. (2021), with modifications for TG tissue processing. Lysis and low-sucrose buffers were prepared fresh on the day of isolation. TG samples pooled for each library were homogenized in 1 mL of ice-cold lysis buffer (320 mM sucrose, 10 mM HEPES, pH 8.0, 5 mM $CaCl_2$, 3 mM Mg-acetate, 0.1 mM EDTA, 1 mM DTT, and 0.1% NP-40) using a prechilled Dounce homogenizer (Fisher Scientific, Cat. #357538). Homogenization consisted of five strokes with the loose pestle followed by ten strokes with the tight pestle. The TGs were processed in batches of four per Dounce homogenization. The resulting nuclei suspensions from the same experimental condition were then combined before the subsequent centrifugation and sorting steps. Homogenized samples were transferred through a 40 μm strainer (Thermo Fisher, Cat. #08-771-1) into low-bind microcentrifuge tubes (Sorenson BioScience, Cat. #11700). Nuclei were subsequently pelleted by centrifugation at 300 × g for 5 min at 4 °C. Following centrifugation, nuclei were washed in 1 mL of low-sucrose buffer lacking NP-40. The pellet was recovered by a second centrifugation and resuspended in 500 μL of 1× PBS containing 1% BSA and 0.2 U/μL SUPERase•In RNase inhibitor (Thermo Fisher, Cat. #AM2696). For neuronal nuclei enrichment, samples were incubated with anti-NeuN antibody (Millipore, Cat. #ABN78, 1:2000–1:5000) for 30 min at 4 °C with gentle rotation, followed by GFP secondary antibody (1:200, 15 min, 4 °C). Nuclei were washed twice with PBS + 1% BSA + SUPERase•In buffer. The nuclear pellet was resuspended in 500 μL FACS buffer (PBS + 1% BSA + 0.2 M sucrose + SUPERase•In), filtered through a 30 μm strainer, and stained with 2.5 μL DRAQ5 (Thermo Fisher, Cat. #62251) for sorting. Fluorescence-activated nuclei sorting (FANS) was performed on a Sony SH800 cell sorter using a 100 µm sorting chip. Nuclei were gated based on scatter properties, followed by singlet discrimination and selection of $DRAQ5^+/GFP^+$ nuclei. Sorted $DRAQ5^+/GFP^+$ nuclei were collected into low-bind tubes and immediately processed using the 10x Genomics Chromium platform. cDNA was generated according to the manufacturer's instructions, quantified before library preparation, and final libraries were submitted for Illumina sequencing.

**Bioinformatics and Data Processing**

Raw snRNA-seq data were processed with Cell Ranger v7.2.0 against the GRCm39/mm39 mouse reference genome. Filtered feature-barcode matrices were imported into R v4.4.3 and analyzed with Seurat v5.0. Seurat objects were created with min.cells = 3 and min.features = 200. Nuclei with <200 detected genes or ≥5% mitochondrial transcripts were excluded, and putative doublets were identified with scDblFinder. Key subtype-specific differential-expression analyses were repeated after doublet exclusion and showed highly concordant effect sizes (Pearson r = 0.997–1.000), with all original DEGs retaining the same direction of effect. Filtered datasets were normalized with SCTransform using glmGamPoi while regressing mitochondrial percentage, followed by PCA, integration where indicated, and shared-nearest-neighbor clustering.

**Cluster Annotation:** Clusters were annotated using canonical TG/DRG markers: NF (*Nefh, Nefm*), NP (*Synpr, Lpar3*), SST (*Osmr, Il31ra*), PEP1 (*Calca, Celf4*), PEP2 (*Gpr26, Trpm8*), and p-cLTMR2 (*Gfra2, Tafa4*). Non-neuronal populations included satellite glia (*Bcan, Lpar1*), Schwann cells (*Mpz, Scn7a*), endothelial cells (*Emcn, Flt1*), pericytes (*Notch3, Rgs5*), and fibroblasts (*Mgp, Dcn*). For the integrated TG atlas, QC-filtered nuclei were merged, normalized with SCTransform/glmGamPoi, and integrated with Harmony by sample identity. UMAP and clustering used the first 30 Harmony dimensions; clustering was evaluated across resolutions 0.1–0.8, with neuronal subtype assignments based on resolutions 0.4 and 0.3 for Day 7 and Day 21, respectively. Canonical marker dot plots used log-normalized RNA expression. Injury module scores were calculated in neuronal nuclei with AddModuleScore using *Atf3, Sprr1a, Sox11, Gap43, Spp1, Gal, Npy,* and *Bdnf* and summarized by subtype, condition, and timepoint. For neuronal visualization, Day 7 and Day 21 annotated neuronal nuclei were merged and UMAP was recalculated using 30 principal components; control nuclei were pooled for visualization, whereas Day 7 and Day 21 CCI nuclei were displayed separately.

**Independent Analyses of Day 7 and Day 21 Cohorts**

Day 7 and Day 21 datasets were analyzed separately by merging control and CCI nuclei within each timepoint, followed by RNA-layer joining, log normalization, variable-feature selection, scaling, PCA, nearest-neighbor analysis, UMAP, and clustering using 30 principal components. Neuronal clusters were annotated as NF, NP, SST, PEP1, PEP2, p-cLTMR2, or Injured based on canonical markers. NF, NP, SST, PEP1, PEP2, and p-cLTMR2 neurons lacking the injury-associated state are referred to as canonical-subtype neurons; this designation reflects transcriptional state rather than anatomical absence of injury. Injured-state neurons were excluded from canonical-subtype analyses unless otherwise indicated. Annotated neuronal nuclei from Day 7 and Day 21 datasets were also merged for cross-timepoint visualization and comparison of neuronal subtype structure.

**Injury-State Canonical Subtype Inference:** Canonical subtype-marker expression was visualized across Day 7 CCI neuronal populations using log-normalized RNA expression. Putative canonical identities of Day 7 CCI injury-state nuclei were inferred by comparison with Day 7 control canonical neurons. The 3,000 most variable control-reference genes were identified, excluding established injury-associated and mitochondrial genes. For each canonical subtype, a mean-expression reference centroid was calculated, and each injury-state nucleus was assigned to the subtype with the highest Spearman correlation. Inferred identities were combined with directly annotated Day 7 CCI canonical neurons to calculate canonical versus injury-associated state proportions within each subtype.

**SST Sensitivity Analysis:** SST representation was summarized as the percentage of recovered neuronal nuclei assigned to the SST subtype within each condition and timepoint. To test whether SST-like CCI nuclei were assigned to other neuronal populations, timepoint-matched control canonical neurons were used as references for Seurat label transfer. Reference subtypes were balanced to the smallest population size (70 nuclei per subtype at Day 7 and 57 at Day 21), injury-associated and mitochondrial genes were excluded from transfer features, and consensus labels were assigned across 20 balanced-reference iterations. Classification sensitivity was tested by withholding 20% of Day 21 control nuclei from each canonical subtype and predicting their identities with the same workflow. Accuracy and prediction scores were summarized descriptively and were not used for biological-replicate inference. Only five Day 21 CCI nuclei were classified as SST-like, while all held-out Day 21 control SST prediction events were correctly classified, indicating that sparse Day 21 SST recovery was not explained by classification failure.

**Differential Expression and Pathway Analysis:** Differential expression between CCI and control nuclei was assessed separately within each neuronal subtype and timepoint using Seurat FindMarkers with MAST, with nCount_RNA and percent.mt included as latent variables. For Figure 2 temporal analyses, DEGs were defined as adjusted $p < 0.05$ and $|\log_2 \text{ fold change}| \geq 0.25$; candidate genes for validation required $\log_2$ fold change $> 0.5$ and adjusted $p < 0.05$. Because sex identities were lost after pooling, sex-linked genes were excluded from candidate prioritization and selected DEG summaries but retained in unfiltered DE tables; their exclusion did not materially alter major results. Within each canonical subtype, significant Day 7 and Day 21 DEGs were classified as Day 7-specific, Day 21-specific, shared in the same direction, or shared in opposite directions. Heatmaps displayed selected top neuronal DEGs after excluding poorly annotated, sex-linked, and likely contaminating genes. Preservation of subtype identity across timepoints was assessed by applying Day 7-trained subtype labels to Day 21 nuclei and summarizing predictions in a row-normalized confusion matrix. GO Biological Process enrichment was performed on significant DEGs using clusterProfiler v4.10 and org.Mm.eg.db with Benjamini–Hochberg correction; enriched terms were visualized using signed combined enrichment scores indicating the direction of enrichment.

**Additional Regulatory and Network Analyses:** For transcription factor gene analysis, pooled canonical neuronal nuclei were analyzed separately at day 7 and day 21 after excluding the injured neuronal cluster. CCI versus control differential expression was performed using Seurat FindMarkers with MAST, with nCount_RNA and percent.mt included as latent variables. Transcription factor and transcription-regulator genes were identified using mouse Gene Ontology annotations for DNA-binding transcription factor activity and transcription regulator activity. Genes were displayed if they were significant in at least one timepoint, using adjusted P < 0.05 and |log2 fold change| ≥ 0.25. Heatmap values represent log2 fold change for CCI versus control, and black points indicate significant differential expression at the corresponding timepoint. For sodium-channel expression analysis, Day 7 and Day 21 neuronal Seurat objects were joined at the RNA assay level, and labeled neuronal nuclei were retained after excluding injured-state neurons. Control nuclei from Day 7 and Day 21 were pooled into a single control group, while CCI nuclei were grouped as Day 7 CCI or Day 21 CCI.

**Subtype-level temporal PCA:** To visualize temporal changes in subtype-specific transcriptional responses, CCI-versus-control differential-expression profiles were compared across NF, NP, PEP1, PEP2, and p-cLTMR2 neurons at Day 7 and Day 21; SST neurons were excluded because of sparse Day 21 recovery. For each subtype and timepoint, genes with positive CCI-versus-control log2 fold change were ranked by effect size, and up to the top 300 genes were retained. Statistical significance was not used as an additional filter for this exploratory PCA. A subtype/timepoint-by-gene matrix was constructed using the union of retained genes across all subtype/timepoint combinations, with genes absent from a given profile assigned a value of 0. Genes with near-zero variance across profiles were removed. Each subtype/timepoint profile was L2-normalized, and principal component analysis was then performed with gene features centered and scaled. Day 7 and Day 21 coordinates for each neuronal subtype were connected to visualize temporal displacement in transcriptional-response space.

**Equal-n temporal robustness analysis:** To evaluate whether temporal differences in CCI-associated transcriptional responses were sensitive to unequal nucleus recovery, NF, NP, PEP1, PEP2, and p-cLTMR2 neurons were analyzed by repeated equal-n downsampling; SST neurons were excluded because only five Day 21 CCI nuclei were recovered. The gene set for each subtype was defined from the union of Day 7 and Day 21 DEGs identified in the full-data Figure 2 analysis using adjusted P < 0.05 and |log2 fold change| ≥ 0.25. For each subtype, the downsampling target was set to the smallest nucleus count among the four Day 7 control, Day 7 CCI, Day 21 control, and Day 21 CCI groups. In each of 200 iterations, nuclei were sampled without replacement from each of the four groups to this common size. CCI-versus-control log2 fold changes for the predefined union gene set were recalculated from the downsampled data, and Spearman correlation between Day 7 and Day 21 effect sizes was calculated within each subtype. The full-data Spearman correlation was calculated using the same union gene set and is shown as a reference.

#### Cross-Dataset Integration and Comparative Analysis

The TG IoN-CCI dataset was compared with Nguyen et al. 2019 (GSE131272; TG infraorbital nerve axotomy, control, Day 7, and Day 21) and Renthal et al. 2020 (GSE154659; DRG sciatic nerve transection, control, Day 7, and Day 28). Raw count matrices were processed in Seurat v5 using SCTransform/glmGamPoi with mitochondrial percentage regressed, followed by PCA, UMAP, clustering, and marker-based annotation using 30 principal components. Unified neuronal labels included NF, PEP1, PEP2, NP, p-cLTMR2, SST, and injured neurons, with injury states identified by markers including *Atf3, Sox11, Gfra1,* and *Gal*. Independently annotated datasets were merged and integrated with Harmony using study and library-preparation type as covariates for visualization only; model-specific subtype annotations were retained for downstream analyses. Timepoints were harmonized as control, acute (Day 7), or chronic (Day 21 for TG datasets; Day 28 for DRG ScNT), and neuronal composition was calculated as the percentage of recovered neurons assigned to each subtype.

To compare DEG burden and account for differences in nuclei recovery, significant DEGs were counted by subtype, injury model, and phase using adjusted $P < 0.05$ and $|\log_2$ fold change$| \geq 0.5$. DEG burden was expressed as DEGs per 100 recovered nuclei: (number of significant DEGs / smaller of the injury and control nuclei counts) × 100. Comparisons with <10 nuclei in either condition were excluded. Heatmap color represents normalized DEG burden, whereas tile values indicate raw DEG counts; injured-state neurons were excluded.

**Additional Cross-Model Visualization and Robustness Analyses:** For Appendix Figure 4, model-specific datasets were merged, normalized using SCTransform/glmGamPoi, and integrated with Harmony using study as the batch covariate. UMAP visualization and clustering were performed using the first 30 Harmony dimensions at resolution 0.3. Local cross-study mixing was assessed using the inverse Simpson diversity index calculated from study identities among each nucleus's 30 nearest neighbors in Harmony space, with higher scores indicating greater mixing. Scores were also calculated in PCA space before integration for comparison. UMAPs were colored by mixing score, injury model, injury phase, and Harmony cluster. Canonical marker expression was assessed using Seurat DotPlot, with dot size representing the percentage of nuclei expressing each marker and color indicating scaled average expression. Neuronal subtype identities were further characterized using a heatmap of scaled average canonical marker expression.

For Appendix Figure 5A, DEG threshold sensitivity was assessed using the cross-model differential-expression table. Genes were filtered to remove sex-linked genes and poorly annotated predicted genes, including Gm* and *Rik genes. DEG counts were summarized across two adjusted P-value thresholds, adjusted P < 0.05 and adjusted P < 0.01. Within each injury model and injury phase, genes were counted as pooled Canonical-neuron DEGs if they met the DEG threshold in at least one canonical neuronal subtype. For Appendix Figure 5B, Gene Ontology over-representation analysis was performed to test whether cross-model enrichment patterns were robust to DEG threshold choice. For each model and injury phase, canonical-neuron DEGs were pooled across subtypes and analyzed separately using DEG thresholds of adjusted P < 0.05 or adjusted P < 0.01. GO Biological Process, Cellular Component, and Molecular Function enrichment analyses were performed using clusterProfiler and org.Mm.eg.db, with all genes tested in the corresponding model and phase used as the background universe. Enrichment P values were corrected using the Benjamini–Hochberg method. Terms from biologically relevant categories, including synaptic organization, excitability, axonal transport, injury and wound responses, metabolism, and neuropeptide signaling, were selected for visualization. Dot size represents gene overlap, and color indicates −log10 false-discovery rate.

For Appendix Figure 5C, transcription factor activity was inferred from subtype-specific injury-associated $\log_2$ fold-change profiles using DoRothEA mouse regulons and the decoupleR univariate linear model. Genes absent from the regulon target set were excluded, and regulons with confidence levels A–D and ≥5 represented target genes were retained. Activity scores were row-scaled across model, phase, and subtype combinations, clipped to the plotted range, and visualized as z scores. For Appendix Figure 5D, subtype-resolved remodeling across IoN-CCI,

TG axotomy, and DRG ScNT was compared by repeated subsampling of 40 injury and 40 control nuclei per subtype, model, and phase. After SCTransform/glmGamPoi normalization with mitochondrial percentage regression, 20 independent subsamples were drawn without replacement and analyzed with MAST. Significant DEGs were defined as adjusted $P < 0.05$ and $|\log_2 \text{fold change}| \geq 0.5$; sex-linked and poorly annotated predicted genes matching Gm* or *Rik were excluded. DEG burden was summarized as the median number of significant DEGs across successful iterations, and comparisons with <40 nuclei in either condition were not tested. Acute comparisons used Day 7 for all models; chronic comparisons used Day 21 for IoN-CCI and TG axotomy and Day 28 for DRG ScNT. Equal-cell subsampling controls for differences in cell number but does not create biological replication or eliminate study-, platform-, or ganglion-specific differences.

**p-cLTMR2 Candidate Prioritization and Robustness Analyses**

Day 21 CCI-versus-control differential-expression responses were compared across NF, PEP1, PEP2, NP, and p-cLTMR2 neurons, excluding injured-state neurons. For each subtype, 100 nuclei per condition were randomly downsampled across 25 iterations; DEG breadth was defined as the number of genes with adjusted $P < 0.05$, and DEG magnitude as the significance-weighted root-mean-square $\log_2$ fold change. Values are shown as median and interquartile range across downsampling iterations. Subtype-specific CCI-versus-control differential expression was performed after RNA-layer joining, SCTransform/glmGamPoi normalization with mitochondrial percentage regression, PrepSCTFindMarkers, and MAST testing. Day 21 p-cLTMR2 candidate genes were prioritized using adjusted $P < 0.05$ and $\log_2$ fold change > 0.5 and considered subtype-selective if not significantly upregulated in other canonical subtypes. Sex-linked and poorly annotated predicted genes, including Gm* and *Rik genes, were excluded from candidate ranking. For functional enrichment, Day 21 p-cLTMR2 DEGs with adjusted $P < 0.05$ and $|\log_2 \text{fold change}| > 0.25$ were analyzed using Enrichr through the enrichR package. Significant terms (Enrichr-adjusted $P < 0.05$) were identified across GO Biological Process, GO Molecular Function, and additional pathway and disease databases. For Figure 4B, representative nonredundant GO terms related to axonal transport, RNA processing, synaptic signaling, and ion transport were selected.

Day 21 p-cLTMR2 DEGs were visualized by average $\log_2$ fold change and adjusted $P$ value, with *Tmem45b* and *Rgs8* highlighted for RNAscope validation. For percent-positive analyses, p-cLTMR2 nuclei were subsetted by timepoint, and nuclei with ≥1 RNA UMI for *Tmem45b* or *Rgs8* were classified as transcript-positive. Percent-positive values and exact binomial 95% confidence intervals were calculated from positive and total nucleus counts. For Figure 4E, condition-by-timepoint interactions were tested using binomial generalized linear models with positive and negative nuclei as the response. Interaction $P$ values were obtained by likelihood-ratio testing against a reduced additive model and adjusted across candidate genes using the Benjamini–Hochberg method. Equal-n downsampling was used to assess whether *Tmem45b* and *Rgs8* induction was robust to unequal p-cLTMR2 recovery. For each gene and timepoint, control and CCI nuclei were downsampled without replacement to the smaller group size, and the CCI-control difference in percent-positive nuclei was recalculated across 5,000 iterations. For chronic-shift analysis, all four condition/timepoint groups were downsampled to the smallest group size, and the chronic shift was calculated as [(Day 21 CCI − Day 21 control)

− (Day 7 CCI − Day 7 control)]. Cross-model comparisons used model-specific controls, with significance defined as adjusted $P < 0.05$ and $|\log_2$ fold change$| \geq 0.5$.

**PEP2 Candidate Prioritization**

Day 21 PEP2 nuclei were analyzed from the neuronal Seurat object after RNA-layer joining and SCTransform/glmGamPoi normalization with mitochondrial transcript percentage regression, followed by PrepSCTFindMarkers. CCI-versus-control differential expression was tested using Seurat FindMarkers with MAST (min.pct = 0.01, logfc.threshold = 0). Candidate PEP2 injury-regulated genes were defined as significantly upregulated with adjusted $P < 0.05$ and $\log_2$ fold change > 0.5. To assess subtype and temporal specificity, CCI-versus-control differential expression was compared across canonical neuronal subtypes at Day 7 and Day 21. Genes were considered PEP2-selective if significantly upregulated in Day 21 PEP2 neurons but not in other canonical subtypes, and Day 21-specific if significant at Day 21 but not Day 7. Functional enrichment of Day 21 PEP2 DEGs (adjusted $P < 0.05$, $|\log_2$ fold change$| > 0.25$) was performed using clusterProfiler and org.Mm.eg.db, with all genes tested in the Day 21 PEP2 analysis as the background. GO Biological Process, Molecular Function, and Cellular Component terms, KEGG, Reactome, and Disease Ontology were tested with Benjamini–Hochberg correction.

For Figure 5A, module scores derived from enriched gene sets were compared between Day 21 CCI PEP2 nuclei and pooled non-PEP2 canonical neurons (NF, NP, PEP1, and p-cLTMR2). The most strongly PEP2-enriched terms were selected for visualization. Significance was assessed by Wilcoxon rank-sum tests with Benjamini–Hochberg correction. For Figure 5B, significant Day 21 PEP2 DEGs were filtered to remove sex-linked, poorly annotated predicted, and likely contaminating genes. Top upregulated and downregulated genes were visualized across individual PEP2 nuclei using control-centered row z-scores, with condition downsampling when needed. For Figure 5C, *Fgf13* and *Kcnmb2* regulation was summarized across canonical neuronal subtypes at Day 7 and Day 21 using CCI-versus-control $\log_2$ fold changes and differences in the percentage of nuclei with detectable expression.

**Additional PEP2 Analyses:** For Appendix Figure 7A–B, robustness of the six Figure 5A functional programs was assessed by repeated equal-n subsampling of Day 21 CCI canonical neurons. For Figure 7A, PEP2, NF, NP, PEP1, and p-cLTMR2 nuclei were subsampled to 148 nuclei per subtype across 1,000 iterations, and PEP2 median module scores were compared with the equal-weighted mean of the four comparator-subtype medians. Figure 7B similarly compared PEP2 with each comparator subtype using pairwise equal-n subsampling. Median differences and 2.5th–97.5th percentile ranges were summarized across iterations. For Figure 7C, GO over-representation analysis was performed on Day 21 PEP2-upregulated genes (adjusted $P < 0.05$, $\log_2$ fold change $\geq 0.25$) using clusterProfiler and org.Mm.eg.db with Benjamini–Hochberg correction; the most significant terms from each ontology and additional terms corresponding to Figure 5A programs were visualized.

For Figure 7D, candidate genes were defined as Day 21 PEP2-upregulated transcripts with adjusted $P < 0.05$ and $\log_2$ fold change > 0.5 and ranked by fold change and the CCI–control difference in transcript-positive nuclei; *Fgf13* and *Kcnmb2* were selected for validation. Sex-linked and poorly annotated transcripts were excluded from candidate prioritization. For

Figure 7E, *Fgf13* and *Kcnmb2* transcript detection was evaluated by 5,000 equal-n subsampling iterations within each timepoint, with CCI–control differences summarized by their mean and 2.5th–97.5th percentile ranges. For Figure 7F, PEP2 differential expression of *Fgf13* and *Kcnmb2* was compared across TG IoN-CCI, TG axotomy, and DRG sciatic nerve transection using harmonized subtype annotations and model-specific controls; significance was defined as adjusted $P < 0.05$ and $|\log_2 \text{fold change}| \geq 0.5$.

**RNAscope Multiplex Fluorescent *In Situ* Hybridization with Immunofluorescence**

RNAscope in situ hybridization was performed using the RNAscope Multiplex Fluorescent v2 assay (Advanced Cell Diagnostics, ACD) following the manufacturer's instructions with modifications from Shiers et al. (2020) adapted for mouse TG. Mice were euthanized by $CO_2$ inhalation followed by decapitation, and TG were dissected bilaterally, embedded in OCT, and stored at −80 °C. TG were cryosectioned at 20 µm onto SuperFrost Plus slides. Sections were fixed in 4% paraformaldehyde/PBS for 15 min at room temperature, dehydrated through 50%, 70%, and 100% ethanol, treated with hydrogen peroxide for 10 min, and incubated with Protease III for 5 s. Target probes were hybridized for 2 h at 40 °C in an ACD HybEZ oven, followed by washing and overnight incubation in 5× SSC at room temperature before amplification.

The following RNAscope probes (Advanced Cell Diagnostics, Newark, California) were used: Channel 1: *Dgkb* (Cat. #1826041-C1), *Tmem45b* (Cat. #420461), *Trpm8* (Cat. #420451); Channel 2: *Nefm* (Cat. #315611-C2), *Fgf13* (Cat. #1303051-C2), *Gfra2* (Cat. #441481-C2); Channel 3: *Trpv1* (Cat. #313331-C3), *Rgs8* (Cat. #515041-C3), and *Kcnmb2* (Cat. #539871-C3). The Mm-Ubc-C3 probe (Cat. #310771-C3) was used as a positive control and the RNAscope 3-plex Negative Control Probe (Cat. #320871) was used as the negative control. RNAscope signal detection and background were assessed using Ubc positive-control and dapB negative-control probes, respectively, with an additional secondary-antibody-only immunofluorescence control (Appendix Fig. 8). Signal development was performed using TSA Vivid™ fluorophores (Thermo Fisher Scientific): TSA Vivid 570 (Cat. #323272) and TSA Vivid 650 (Cat. #323273). AMP1–3 amplification and sequential HRP-mediated C1–C3 signal development were performed according to the manufacturer's protocol. Sections were then immediately processed for immunofluorescence to permit co-detection of protein markers.

**Immunofluorescence Staining:** Sections were washed three times in 1× PBS (5 min each) at room temperature, then blocked and permeabilized in 10% normal goat serum (NGS; Sigma-Aldrich, G9023) with 0.3% Triton X-100 (BioUltra, 93443) in 0.1 M of phosphate buffer (PB; pH 7.4) for 1 h at room temperature. Primary antibody incubation was performed in blocking solution in a humidified chamber overnight at 4 °C. For neuronal cell body labeling, sections were incubated with rabbit monoclonal anti-NeuN antibody (ab177487, Abcam; 1:400) overnight at 4 °C. Control sections received blocking solution alone. Following three washes in PBS (5 min each), sections were incubated for 1 h at room temperature with Alexa Fluor 488-conjugated secondary antibody diluted 1:2000 in blocking solution and counterstained with DAPI (1:5000) to label nuclei. After a final wash in PBS, slides were coverslipped with ProLong Gold antifade mountant (Fisher Scientific, P36930) and cured for 24 h at room temperature in the dark before sealing with nail polish. Slides were stored at 4 °C for short-term use or at −20 °C/−80 °C for long-term storage.

**Confocal Imaging**

Confocal images were acquired using a Nikon A1RHD laser scanning confocal microscope with a 40× oil immersion objective (CFI Plan Apo λ, NA 1.3) and NIS-Elements software. Large TG sections were imaged using the advanced large-image tiling workflow. Tissue boundaries were mapped by low-resolution preview imaging, and the tile acquisition area was defined in JOBS Explorer using the "high mag tiling via boundary marking" function. Images were acquired by ND acquisition using XY tile positions and Z-stacks with resonant bidirectional scanning at 1024 × 1024-pixel resolution and 1.5× optical zoom. Four fluorescence channels were acquired: 405 nm for DAPI, 488 nm for Alexa Fluor 488/NeuN, 555 nm for TSA Vivid 570, and 647 nm for TSA Vivid 650. Z-stacks were acquired over a 20-μm range with 11 optical sections at approximately 2-μm spacing. After acquisition, images were processed in NIS-Elements using Denoise.ai, stitched using Stitch Multipoint to Large Image, and converted to maximum-intensity projections. Stitched projection images were exported for downstream Fiji/ImageJ quantification. The same imaging and processing workflow was applied across paired control and IoN-CCI tissues within each probe set. For figure presentation, neuronal cell bodies were manually outlined using the NeuN immunofluorescence channel, and the outlines were overlaid on merged RNAscope images for visualization only; these outlines were not used for quantitative image analysis (Appendix Fig. 12). Whole-section RNAscope images of the trigeminal ganglion for *Tmem45b* and *Rgs8* in *Gfra2*-positive neurons and *Fgf13* and *Kcnmb2* in *Trpm8*-positive neurons are provided in Appendix Figures 10 and 11.

**Image Analysis**

For each mouse, one section from the ipsilateral IoN-CCI TG and one section from the matched contralateral sham-control TG were analyzed. The entire TG cross-section was imaged by tiled confocal acquisition and quantified without selective field or subregion sampling, yielding one whole-section measurement per condition per mouse. Fluorescence images were analyzed in ImageJ/Fiji using a custom macro that automated channel separation, maximum-intensity projection, background subtraction, thresholding, binary mask generation, ROI selection, and measurement export. Four-channel image stacks were separated into DAPI, NeuN, RNA1, and RNA2 channels. NeuN images were processed by rolling-ball background subtraction, unsharp masking, median filtering, and RenyiEntropy thresholding to generate a binary neuronal mask. This mask was used to restrict nuclear ROI detection to NeuN-positive regions and was not interpreted as complete neuronal soma segmentation. DAPI images were processed independently using background subtraction, unsharp masking, median filtering, maximum-intensity projection, contrast enhancement, Otsu thresholding, dilation, and watershed separation. DAPI-positive nuclei within the NeuN-positive mask were retained as neuronal nuclear ROIs and enlarged by 10 pixels to capture perinuclear cytoplasmic RNAscope signal; this expansion was used only for RNA quantification and not as full cell-body segmentation.

RNAscope channels were maximum-projected and processed using background subtraction, unsharp masking, convolution filtering, contrast enhancement, and MaxEntropy thresholding. Binary puncta masks were generated using predefined particle-size thresholds applied consistently within each probe set. Transcript-positive cells were classified by overlap between the expanded neuronal ROI and the corresponding RNAscope binary mask; ROIs with a maximum binary-mask value of 255 were classified as positive and those with a value of 0 as negative. CSV files exported from Fiji were merged using a custom Excel VBA workflow. Count summaries were used to quantify neuronal ROIs, RNA1- and RNA2-positive ROIs, double-positive ROIs, and single-positive subsets. Percent-positive values were calculated for each section and averaged at the mouse level before statistical comparison. Quantification was not performed blinded to experimental condition; however, all images were analyzed using the same automated pipeline, with predefined thresholding, ROI generation, neuronal gating, transcript-positive classification, and count-summary procedures applied consistently across paired control and IoN-CCI tissues.

**Statistical Analysis**

All statistical analyses were performed in R v4.4.3. Single-nucleus RNA-seq data were analyzed using Seurat v5, with differential expression tested using MAST. Genes with adjusted $P < 0.05$ and $|\log_2 FC| \geq 0.5$ were considered differentially expressed unless otherwise specified, with multiple-testing correction performed by the corresponding differential-expression or enrichment workflow. RNAscope quantification was performed from exported image-analysis count tables, with the primary outcome defined as the percentage of marker-defined neurons positive for each candidate transcript. Percentages were calculated for each section and averaged at the mouse level for each condition. Paired control and IoN-CCI comparisons were analyzed using paired *t* tests, with paired Wilcoxon signed-rank tests as nonparametric sensitivity analyses. Count-based validation used binomial mixed-effects logistic regression with transcript-positive and transcript-negative neurons as the response, condition as a fixed effect, and mouse identity as a random intercept. Biological replicates were individual mice; sections or images from the same mouse were technical replicates. Nucleus-level downsampling and resampling were used only as sensitivity analyses and not for biological-replicate inference. Sample sizes are reported in figure legends, and data are shown as mean ± SEM unless otherwise indicated. Statistical significance was defined as $P < 0.05$ or adjusted $P < 0.05$, as appropriate.

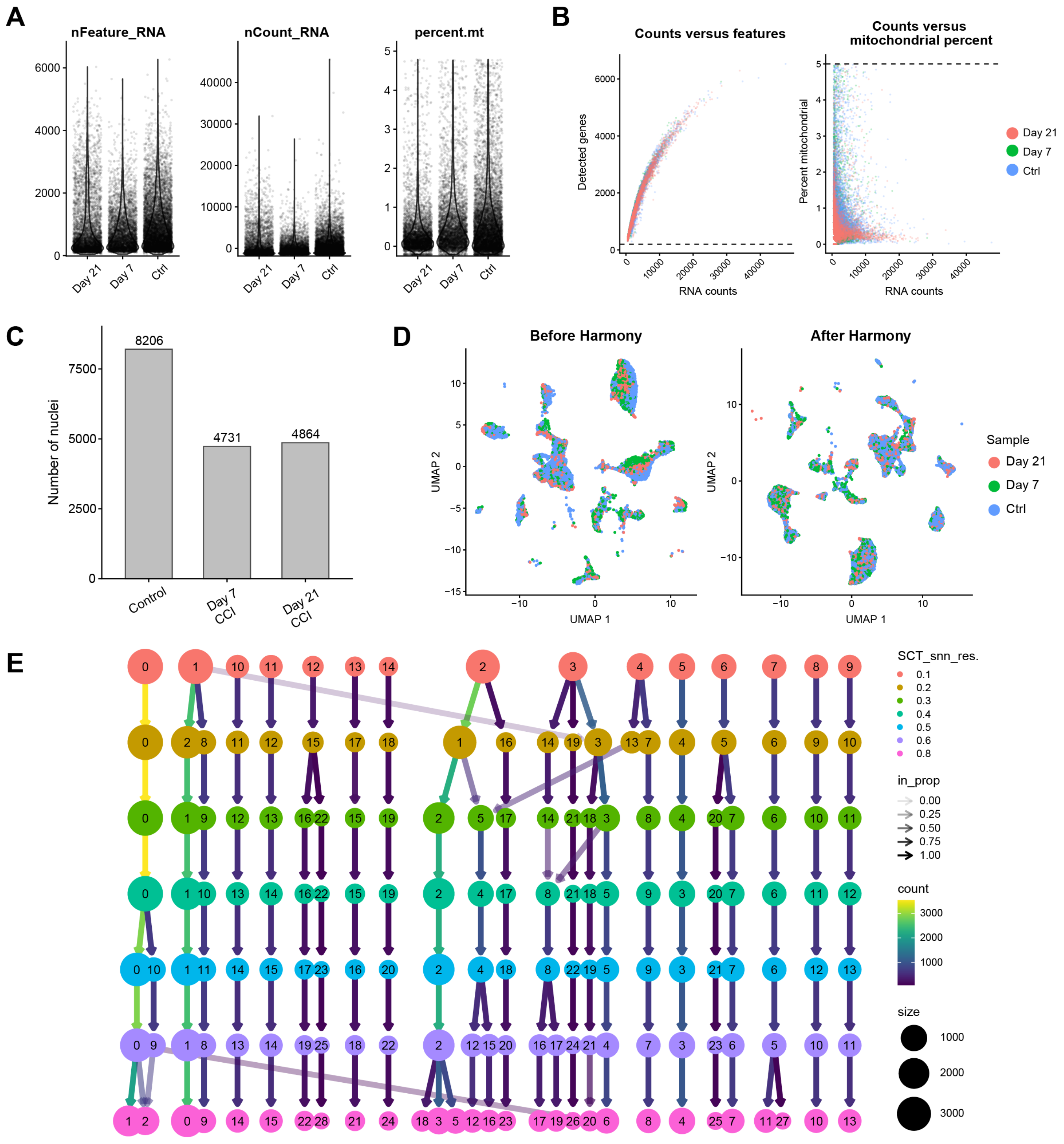
A
nFeature_RNA
nCount_RNA
percent.mt
Day 21
Day 7
Ctrl
B
Counts versus features
Counts versus mitochondrial percent
Detected genes
Percent mitochondrial
RNA counts
Day 21
Day 7
Ctrl
C
Number of nuclei
8206
4731
4864
Control
Day 7 CCI
Day 21 CCI
D
Before Harmony
After Harmony
UMAP 1
UMAP 2
Sample
Day 21
Day 7
Ctrl
E
SCT_snn_res.
0.1
0.2
0.3
0.4
0.5
0.6
0.8
in_prop
0.00
0.25
0.50
0.75
1.00
count
3000
2000
1000
size
1000
2000
3000

**Appendix Figure 1. Quality control and integration assessment for the integrated trigeminal ganglion snRNA-seq atlas (related to Figure 1). (A)** Violin plots showing the distribution of detected genes, total RNA counts, and mitochondrial transcript percentage across nuclei grouped by experimental timepoint/condition. Each point represents an individual nucleus. **(B)** QC scatter plots showing the relationship between total RNA counts and detected genes, and between total RNA counts and mitochondrial transcript percentage. Dashed horizontal lines indicate filtering thresholds used to exclude low-quality nuclei or nuclei with high mitochondrial content. **(C)** Bar plot showing the number of nuclei retained for analysis in each experimental group after quality control filtering. Numbers above bars indicate the total nuclei recovered per group. **(D)** UMAP visualization of selected annotated neuronal and non-neuronal populations before and after Harmony integration, colored by sample group. The post-Harmony embedding shows improved mixing across control, day 7 CCI, and day 21 CCI samples while preserving cell population structure. **(E)** Clustree analysis showing cluster relationships across increasing Seurat clustering resolutions. Nodes represent clusters at each resolution, with node size corresponding to the number of nuclei and edges indicating the proportion of nuclei shared between clusters across resolutions. The analysis was used to guide selection of the final clustering resolution for cell-type annotation.

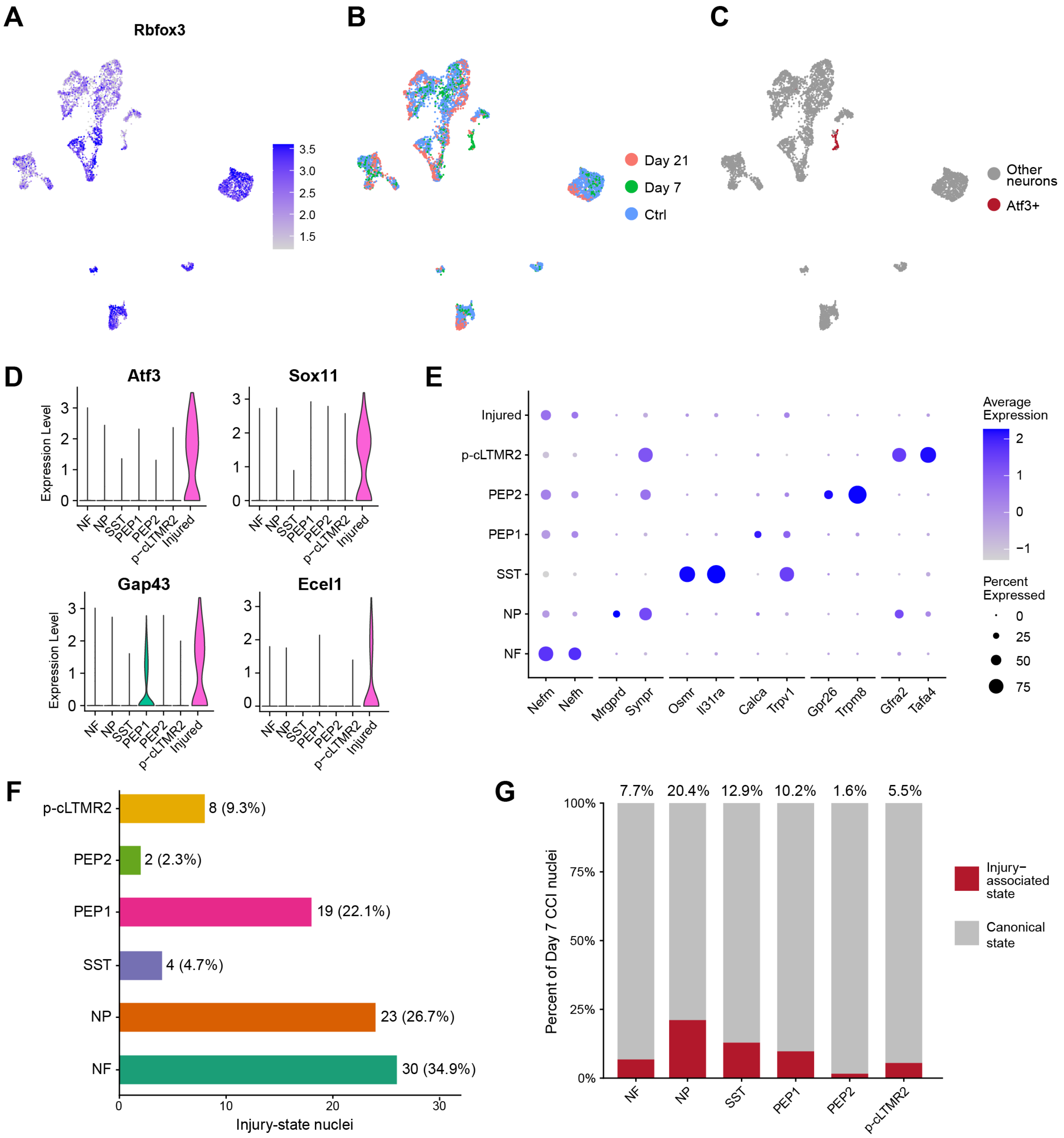
A
Rbfox3
3.5
3.0
2.5
2.0
1.5
B
Day 21
Day 7
Ctrl
C
Other neurons
Atf3+
D
Atf3
Sox11
Gap43
Ecel1
Expression Level
NF
NP
SST
PEP1
PEP2
p-cLTMR2
Injured
E
Injured
p-cLTMR2
PEP2
PEP1
SST
NP
NF
Nefm
Nefh
Mrgprd
Synpr
Osmr
Il31ra
Calca
Trpv1
Gpr26
Trpm8
Gfra2
Tafa4
Average Expression
Percent Expressed
F
8 (9.3%)
2 (2.3%)
19 (22.1%)
4 (4.7%)
23 (26.7%)
30 (34.9%)
Injury-state nuclei
G
7.7%
20.4%
12.9%
10.2%
1.6%
5.5%
Percent of Day 7 CCI nuclei
Injury-associated state
Canonical state

**Appendix Figure 2. Injury-associated neurons retain heterogeneous canonical sensory-neuron signatures after IoN-CCI (related to Figure 1). (A)** Feature plot showing expression of the pan-neuronal marker *Rbfox3* across the neuronal subset UMAP. **(B)** UMAP visualization of the neuronal subset colored by experimental group, including pooled control, Day 7 CCI, and Day 21 CCI nuclei. **(C)** UMAP highlighting *Atf3*-positive nuclei within the neuronal subset. *Atf3*-positive nuclei are shown in red and all other neuronal nuclei in gray. **(D)** Violin plots showing expression of the injury-associated genes *Atf3, Sox11, Gap43,* and *Ecel1* across neuronal subtypes. Expression of these genes was concentrated in the injury-associated neuronal cluster, with comparatively limited expression across canonical sensory-neuron subtypes. **(E)** Dot plot showing canonical sensory-neuron marker expression across Day 7 CCI neuronal populations, including the injury-associated cluster. Marker sets included *Nefm/Nefh* (NF), *Mrgprd/Synpr* (NP), *Osmr/Il31ra* (SST), *Calca/Trpv1* (PEP1), *Gpr26/Trpm8* (PEP2), and *Gfra2/Tafa4* (p-cLTMR2). Dot size indicates the percentage of nuclei expressing each gene and color indicates scaled average expression. **(F)** Putative canonical subtype identities of the 86 Day 7 CCI injury-state nuclei inferred by transcriptomic similarity to canonical Day 7 control neurons. For each canonical subtype, a control reference centroid was calculated from normalized expression of variable genes after exclusion of injury-associated and mitochondrial genes, and each injury-state nucleus was assigned to the subtype with the highest Spearman correlation. Bars indicate the number of injury-state nuclei assigned to each putative canonical subtype, with labels showing number and percentage of the injury-state population. The inferred assignments comprised 30 NF-like (34.9%), 23 NP-like (26.7%), 4 SST-like (4.7%), 19 PEP1-like (22.1%), 2 PEP2-like (2.3%), and 8 p-cLTMR2-like (9.3%) nuclei. **(G)** Relative proportions of recovered Day 7 CCI nuclei assigned to canonical versus injury-associated transcriptional states within each inferred sensory-neuron subtype. Canonical-state nuclei retain their directly annotated canonical subtype, whereas injury-state nuclei are grouped according to the inferred canonical identity shown in F. Injury-associated nuclei represented 7.7% of inferred NF, 20.4% of NP, 12.9% of SST, 10.2% of PEP1, 1.6% of PEP2, and 5.5% of p-cLTMR2 recovered nuclei.

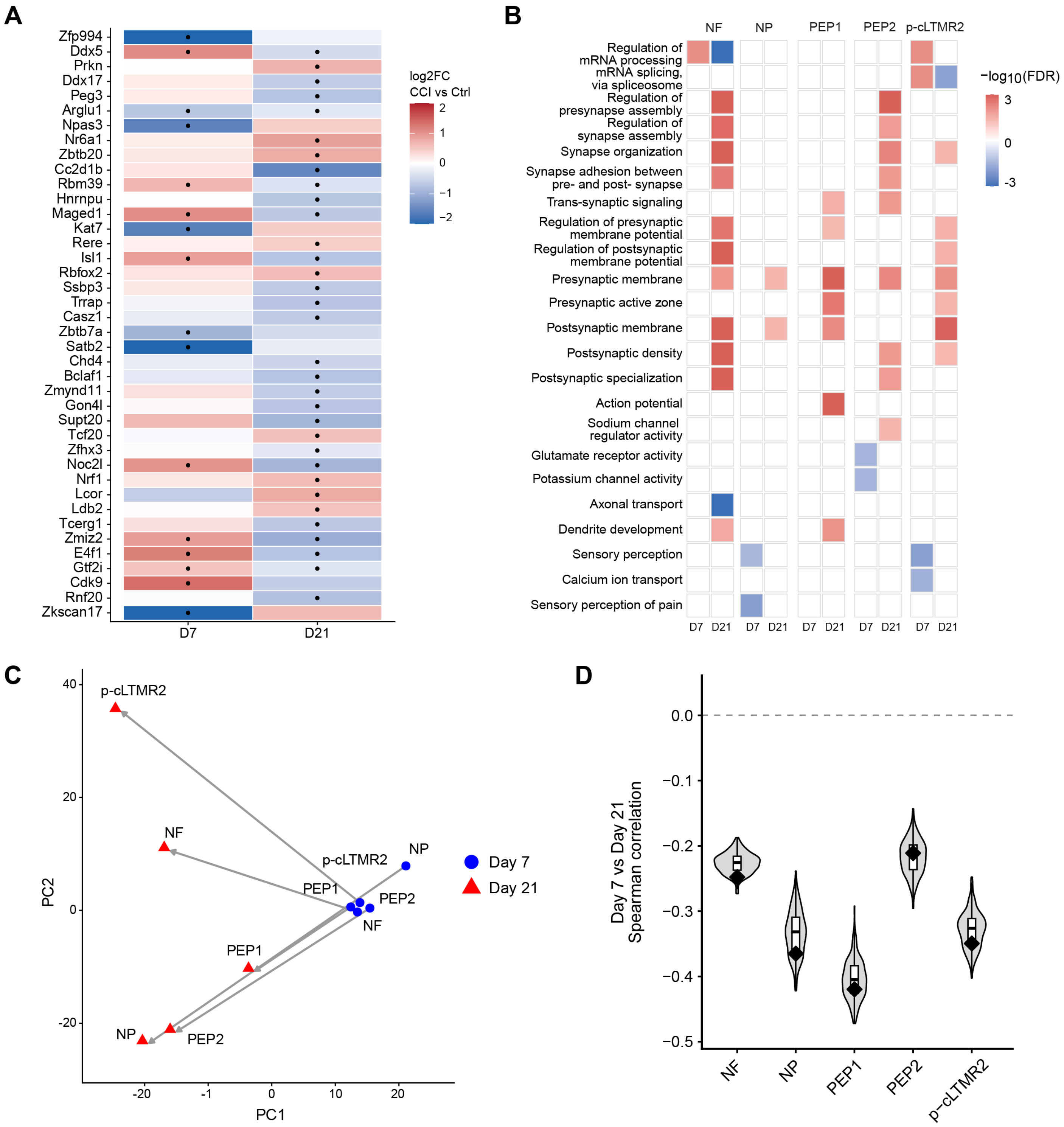
A
Zfp994
Ddx5
Prkn
Ddx17
Peg3
Arglu1
Npas3
Nr6a1
Zbtb20
Cc2d1b
Rbm39
Hnrnpu
Maged1
Kat7
Rere
Isl1
Rbfox2
Ssbp3
Trrap
Casz1
Zbtb7a
Satb2
Chd4
Bclaf1
Zmynd11
Gon4l
Supt20
Tcf20
Zfhx3
Noc2l
Nrf1
Lcor
Ldb2
Tcerg1
Zmiz2
E4f1
Gtf2i
Cdk9
Rnf20
Zkscan17
D7
D21
log2FC
CCI vs Ctrl
2
1
0
−1
−2
B
NF
NP
PEP1
PEP2
p-cLTMR2
Regulation of mRNA processing
mRNA splicing, via spliceosome
Regulation of presynapse assembly
Regulation of synapse assembly
Synapse organization
Synapse adhesion between pre- and post- synapse
Trans-synaptic signaling
Regulation of presynaptic membrane potential
Regulation of postsynaptic membrane potential
Presynaptic membrane
Presynaptic active zone
Postsynaptic membrane
Postsynaptic density
Postsynaptic specialization
Action potential
Sodium channel regulator activity
Glutamate receptor activity
Potassium channel activity
Axonal transport
Dendrite development
Sensory perception
Calcium ion transport
Sensory perception of pain
D7
D21
−log10(FDR)
3
0
-3
C
p-cLTMR2
NF
NP
PEP1
PEP2
Day 7
Day 21
PC2
PC1
40
20
0
-20
-1
10
D
Day 7 vs Day 21
Spearman correlation
0.0
−0.1
−0.2
−0.3
−0.4
−0.5
NF
NP
PEP1
PEP2
p-cLTMR2

**Appendix Figure 3. Complementary analyses supporting temporal transcriptional and signaling remodeling after IoN-CCI (related to Figure 2). (A)** Heatmap showing significant differentially expressed transcription factor genes in pooled canonical subtype neurons at day 7 and day 21 after CCI. Tile color indicates log2 fold change in CCI relative to control, with red indicating upregulation and blue indicating downregulation. Black dots mark genes meeting the significance threshold at the corresponding timepoint. **(B)** Subtype-resolved Gene Ontology (GO) Biological Process enrichment of CCI-regulated genes at Day 7 and Day 21. Upregulated and downregulated DEGs were analyzed separately using the corresponding tested genes as background. Sex-associated and poorly characterized genes were excluded. Tiles show signed −log10(FDR), with red indicating enrichment among upregulated DEGs, blue indicating enrichment among downregulated DEGs, and white indicating no significant enrichment. Representative terms highlight acute RNA-processing programs and chronic synaptic, excitability, axonal transport, and structural remodeling**. (C)** Principal component analysis of subtype-level differential-expression profiles in canonical sensory neuronal subtypes at acute and chronic timepoints. The PCA space was defined using NF, NP, PEP1, PEP2, and p-cLTMR2 profiles. Points represent neuronal subtypes, with blue circles indicating Day 7 and red triangles indicating Day 21. Arrows connect the same neuronal subtype across timepoints, illustrating temporal shifts in injury-associated transcriptional remodeling. **(D)** Robustness of temporal transcriptional divergence to unequal nucleus recovery. CCI and control nuclei were randomly downsampled to equal numbers within each analyzed neuronal subtype and timepoint over 200 iterations. Violin plots show the distribution of Spearman correlations between Day 7 and Day 21 CCI-versus-control log2FC values across the union of temporal DEGs; internal boxplots indicate the median and interquartile range, and black diamonds denote correlations from the complete dataset.

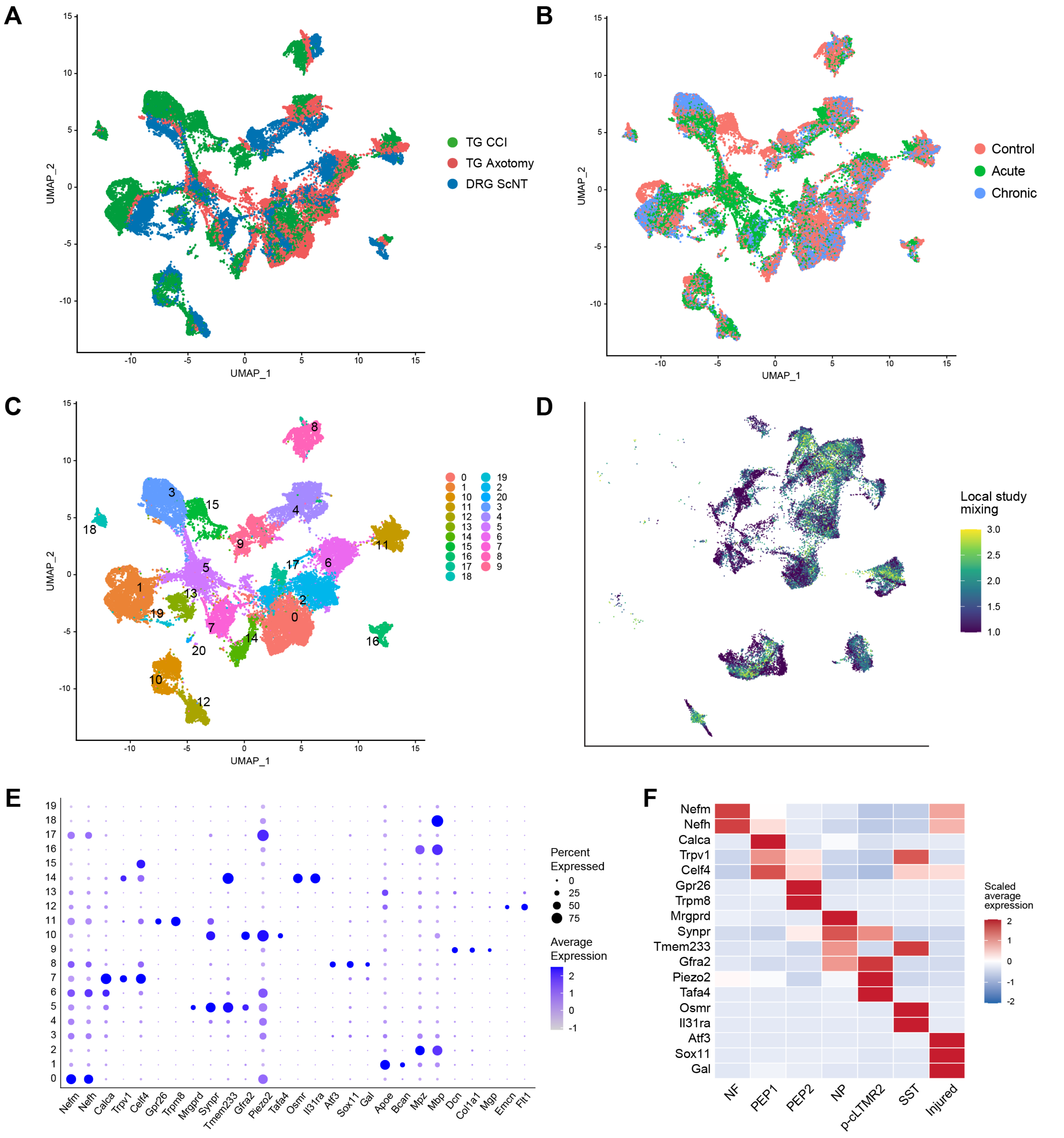
A
TG CCI
TG Axotomy
DRG ScNT
UMAP_1
UMAP_2
B
Control
Acute
Chronic
UMAP_1
UMAP_2
C
UMAP_1
UMAP_2
D
Local study mixing
3.0
2.5
2.0
1.5
1.0
E
Percent Expressed
0
25
50
75
Average Expression
2
1
0
-1
Nefm
Nefh
Calca
Trpv1
Celf4
Gpr26
Trpm8
Mrgprd
Synpr
Tmem233
Gfra2
Piezo2
Tafa4
Osmr
Il31ra
Atf3
Sox11
Gal
Apoe
Bcan
Mpz
Mbp
Dcn
Col1a1
Mgp
Emcn
Flt1
F
Scaled average expression
NF
PEP1
PEP2
NP
p-cLTMR2
SST
Injured

**Appendix Figure 4. Combined atlas integration and neuronal subtype annotation across injury models (related to Figure 3). (A)** All-cell Harmony-integrated UMAP colored by injury model, including TG IoN-CCI, TG IoN axotomy, and DRG ScNT datasets. **(B)** All-cell Harmony-integrated UMAP colored by injury phase, including control, acute (day 7), and chronic (day 21 and 28) timepoints. **(C)** UMAP visualization of all-cell Harmony clusters at resolution 0.3. Harmony clusters were used for cross-dataset visualization, while neuronal subtype identities were retained from the independently annotated datasets. **(D)** Harmony-integrated UMAP restricted to neuronal nuclei retained for cross-model comparison, colored by local cross-study mixing. Mixing was quantified using the inverse Simpson diversity index of study identities among each nucleus's 30 nearest neuronal neighbors in the 30-dimensional Harmony embedding. **(E)** Dot plot showing canonical neuronal and non-neuronal marker expression across Harmony clusters. Dot size represents the percentage of nuclei expressing each marker, and color indicates average scaled expression. **(F)** Heatmap showing scaled average expression of canonical sensory neuron and injury-state markers across annotated neuronal subtypes retained for cross-model comparison.

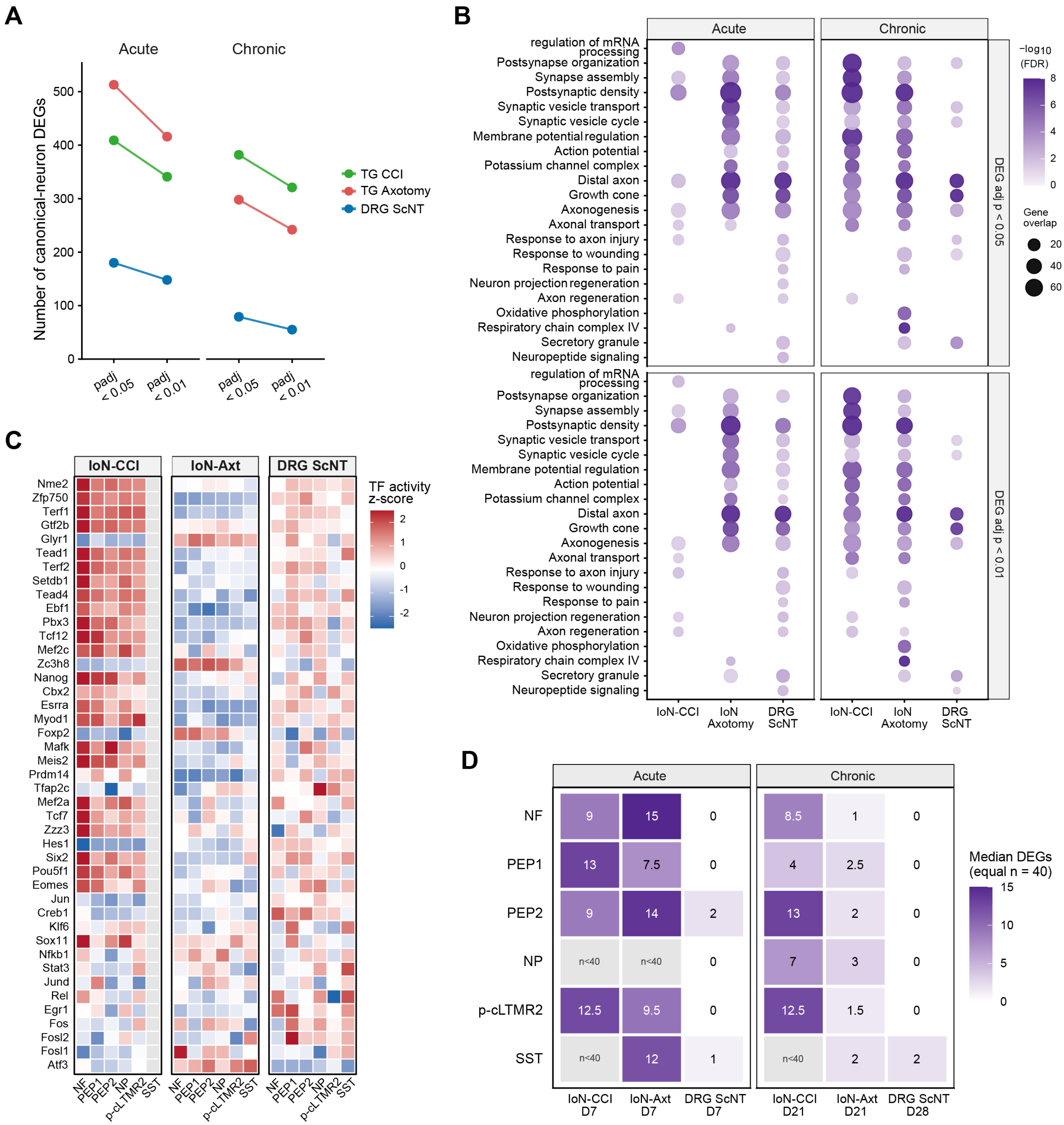


| | IoN-CCI D7 | IoN-Axt D7 | DRG ScNT D7 | IoN-CCI D21 | IoN-Axt D21 | DRG ScNT D28 |
|---|---|---|---|---|---|---|
| NF | 9 | 15 | 0 | 8.5 | 1 | 0 |
| PEP1 | 13 | 7.5 | 0 | 4 | 2.5 | 0 |
| PEP2 | 9 | 14 | 2 | 13 | 2 | 0 |
| NP | n<40 | n<40 | 0 | 7 | 3 | 0 |
| p-cLTMR2 | 12.5 | 9.5 | 0 | 12.5 | 1.5 | 0 |
| SST | n<40 | 12 | 1 | n<40 | 2 | 2 |

**Appendix Figure 5. Robustness and regulatory features of canonical-neuron transcriptional remodeling across injury models (related to Figure 3). (A)** Pooled canonical-neuron DEG counts across adjusted p-value thresholds. DEGs were pooled across NF, PEP1, PEP2, NP, p-cLTMR2, and SST neurons for each injury model and phase. Counts are shown using DEG adjusted $p < 0.05$ and a more stringent DEG adjusted $p < 0.01$ threshold. Line plots show that the relative model-specific DEG patterns are preserved across thresholds for acute and chronic comparisons. **(B)** GO enrichment threshold-sensitivity analysis for pooled canonical-neuron DEGs. GO terms were selected from biologically relevant categories including synaptic organization, excitability, axonal transport, injury/wounding/pain responses, metabolism, and neuropeptide signaling. Enrichment was performed separately using DEGs defined at adjusted $p < 0.05$ or adjusted $p < 0.01$. Dot size represents gene overlap, and color indicates −log10(FDR). **(C)** Inferred transcription factor activity across chronic canonical-neuron subtypes in IoN-CCI, IoN axotomy, and DRG ScNT models. TF activity was inferred from subtype-specific injury log2 fold-change profiles using DoRothEA/decoupleR regulons. Rows show selected transcription factors, columns show canonical neuronal subtypes, and color indicates row-scaled TF activity z-score. **(D)** Subtype-resolved differential-expression burden across injury models after equal-cell subsampling. Canonical neuronal subtypes were compared across infraorbital nerve chronic constriction injury (IoN-CCI), infraorbital nerve axotomy (IoN axotomy), and dorsal root ganglion sciatic nerve transection (DRG ScNT) datasets at acute and chronic injury stages. For each neuronal subtype, model, and injury phase, injury and model-specific control nuclei were repeatedly subsampled to equal numbers ($n = 40$ per condition) and differential expression was assessed using MAST. Significant differentially expressed genes (DEGs) were defined as adjusted $P < 0.05$ and $|\log2 \text{ fold change}| \geq 0.5$. Tile color and numbers indicate the median number of significant DEGs across 20 repeated equal-*n* subsampling iterations. Comparisons with fewer than 40 nuclei in either condition were not tested and are shown in gray. Acute injury corresponds to Day 7 for all three models; chronic injury corresponds to Day 21 for IoN-CCI and IoN axotomy and Day 28 for DRG ScNT. Injured-state neurons were excluded from the analysis.

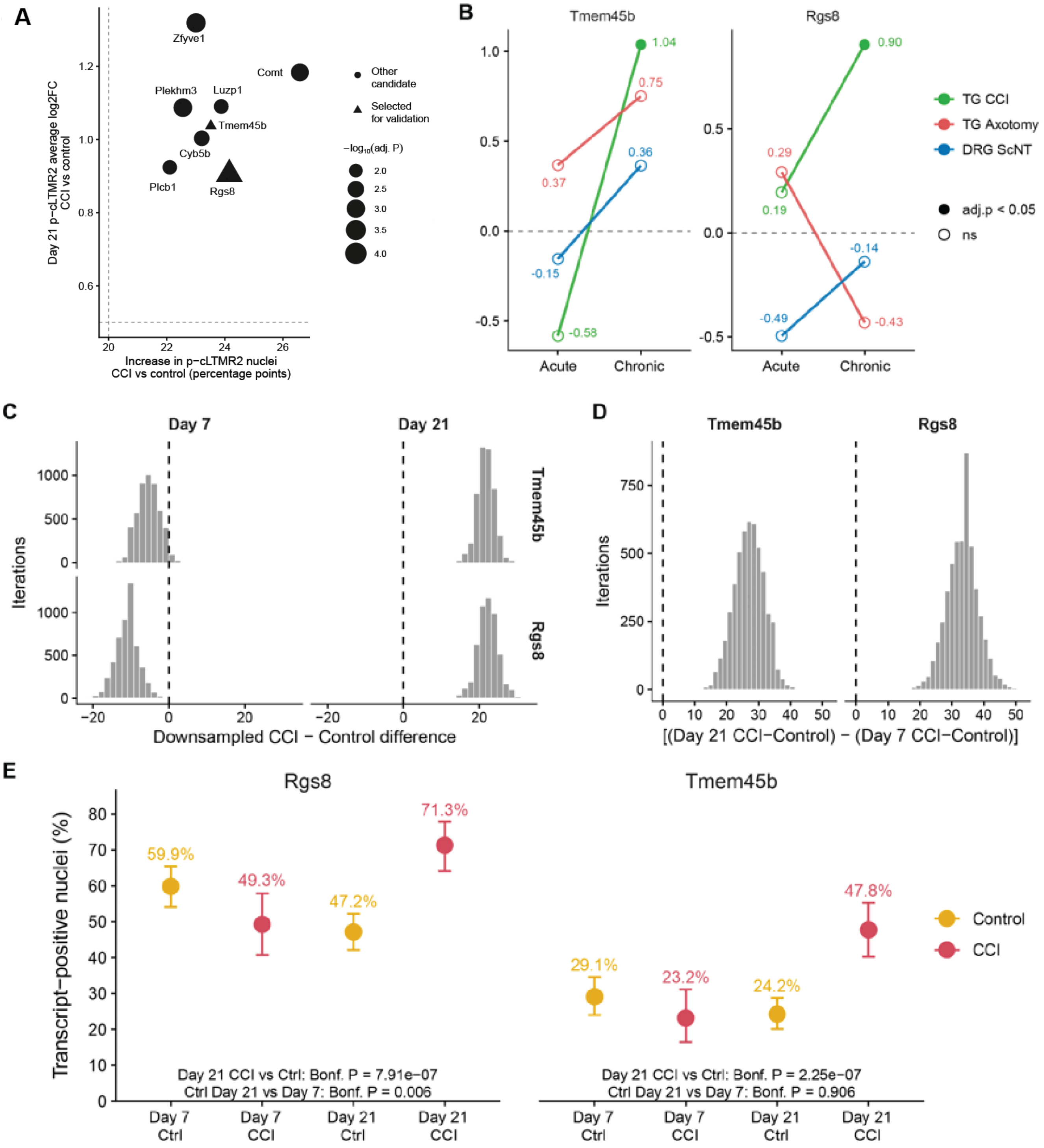
A
Zfyve1
Comt
Plekhm3
Luzp1
Tmem45b
Cyb5b
Plcb1
Rgs8
Day 21 p−cLTMR2 average log2FC
CCI vs control
Increase in p−cLTMR2 nuclei
CCI vs control (percentage points)
Other candidate
Selected for validation
−log10(adj. P)
B
Tmem45b
Rgs8
TG CCI
TG Axotomy
DRG ScNT
adj.p < 0.05
ns
Acute
Chronic
C
Day 7
Day 21
Iterations
Downsampled CCI − Control difference
D
Tmem45b
Rgs8
Iterations
[(Day 21 CCI−Control) − (Day 7 CCI−Control)]
E
Rgs8
Tmem45b
Transcript−positive nuclei (%)
59.9%
49.3%
47.2%
71.3%
29.1%
23.2%
24.2%
47.8%
Control
CCI
Day 21 CCI vs Ctrl: Bonf. P = 7.91e−07
Ctrl Day 21 vs Day 7: Bonf. P = 0.006
Day 21 CCI vs Ctrl: Bonf. P = 2.25e−07
Ctrl Day 21 vs Day 7: Bonf. P = 0.906
Day 7 Ctrl
Day 7 CCI
Day 21 Ctrl
Day 21 CCI

**Appendix Figure 6. Candidate prioritization and robustness of chronic p-cLTMR2 transcriptional remodeling after IoN-CCI (related to Figure 4). (A)** Candidate-gene prioritization plot for Day 21 p-cLTMR2 neurons. Genes are plotted by the increase in the percentage of expressing p-cLTMR2 neurons in CCI relative to control and by average log2 fold change in CCI versus control. Dashed lines indicate candidate-selection thresholds. Point size represents −log10 adjusted p-value. Triangles indicate genes selected for RNAscope validation, and circles indicate secondary candidate genes. **(B)** Cross-model targeted differential-expression comparison of Tmem45b and Rgs8 in p-cLTMR2 neurons. Line plots show log2 fold change relative to model-specific controls at acute and chronic timepoints for TG CCI, TG axotomy, and DRG ScNT models. Filled points indicate adjusted $p < 0.05$ and $|\log2FC| \geq 0.5$, and open points indicate non-significant comparisons. IoN-CCI values were calculated from the original p-cLTMR2 cluster-level analyses, whereas TG axotomy and DRG ScNT values were calculated using targeted differential-expression testing in the integrated p-cLTMR2 annotations. **(C)** Equal-n downsampling analysis of CCI-Control percent-positive differences for *Tmem45b* and *Rgs8* within each timepoint. For each gene and timepoint, Control and CCI p-cLTMR2 nuclei were repeatedly downsampled to equal group sizes, and the CCI-Control difference in percent-positive nuclei was recalculated across iterations. Dashed vertical lines indicate zero difference. Day 21 distributions were shifted above zero for both *Tmem45b* and *Rgs8*. **(D)** Equal-n chronic-shift analysis for Tmem45b and Rgs8. Chronic shift was calculated as [(Day 21 CCI − Day 21 Control) − (Day 7 CCI − Day 7 Control)] after repeated equal-n downsampling across all four groups. Chronic-shift distributions were positive for both genes. **(E)** Four-group percent-positive validation plot for Rgs8 and Tmem45b in p-cLTMR2 nuclei. Points indicate the percentage of p-cLTMR2 nuclei with detectable transcript (RNA UMI count > 0), and error bars indicate exact 95% binomial confidence intervals. Bonferroni-adjusted pairwise comparisons among the four groups were performed using count-level binomial generalized linear models. Both Rgs8 and Tmem45b showed significant Day 21 CCI-associated increases relative to Day 21 Control. Rgs8 also differed significantly between Day 7 and Day 21 Control, whereas Tmem45b showed no significant Control difference between timepoints.

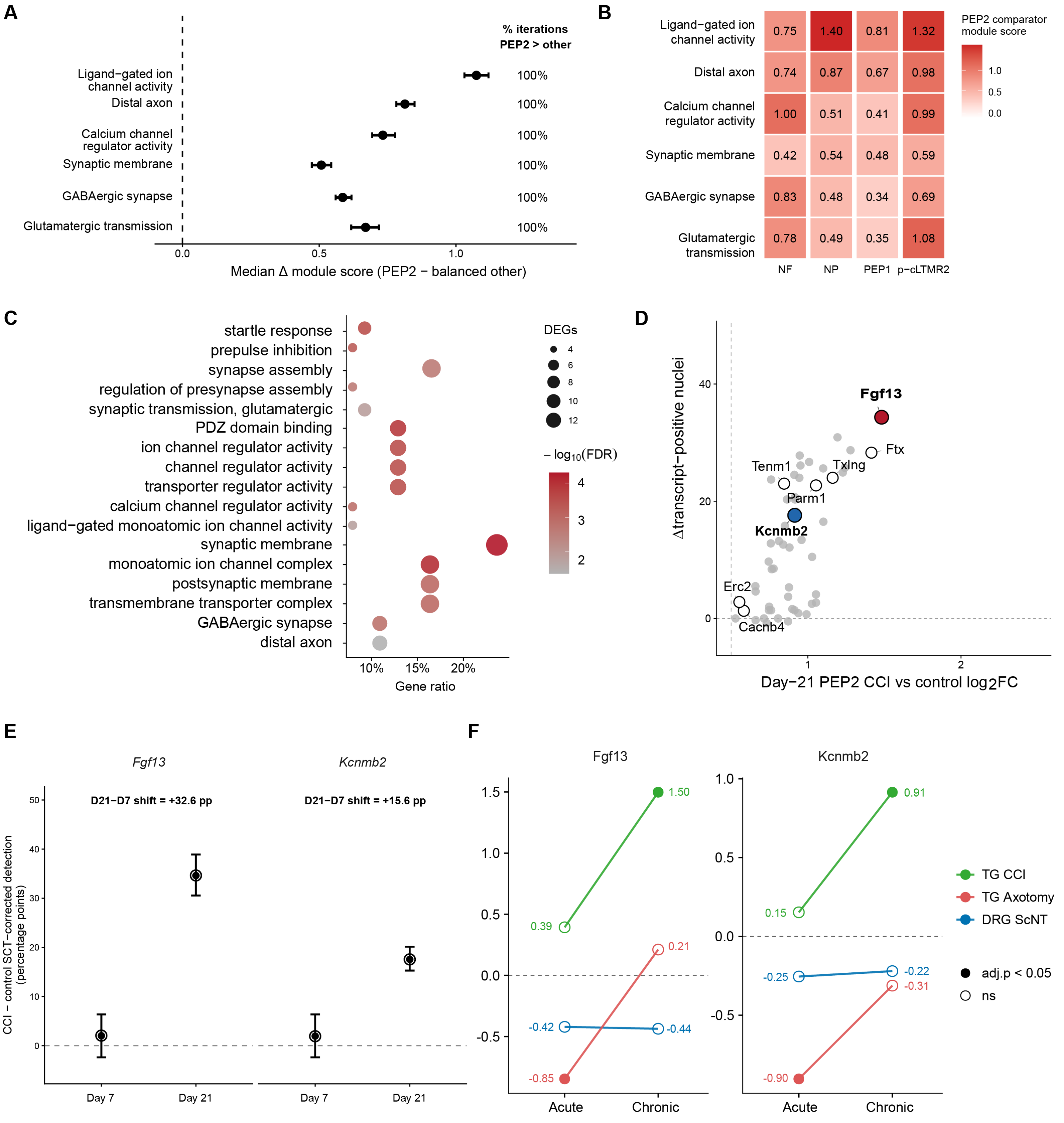
A
% iterations
PEP2 > other
Ligand−gated ion
channel activity
Distal axon
Calcium channel
regulator activity
Synaptic membrane
GABAergic synapse
Glutamatergic transmission
100%
100%
100%
100%
100%
100%
0.0
0.5
1.0
Median Δ module score (PEP2 − balanced other)
B
Ligand−gated ion channel activity 0.75 1.40 0.81 1.32
Distal axon 0.74 0.87 0.67 0.98
Calcium channel regulator activity 1.00 0.51 0.41 0.99
Synaptic membrane 0.42 0.54 0.48 0.59
GABAergic synapse 0.83 0.48 0.34 0.69
Glutamatergic transmission 0.78 0.49 0.35 1.08
NF
NP
PEP1
p−cLTMR2
PEP2 comparator
module score
1.0
0.5
0.0
C
startle response
prepulse inhibition
synapse assembly
regulation of presynapse assembly
synaptic transmission, glutamatergic
PDZ domain binding
ion channel regulator activity
channel regulator activity
transporter regulator activity
calcium channel regulator activity
ligand−gated monoatomic ion channel activity
synaptic membrane
monoatomic ion channel complex
postsynaptic membrane
transmembrane transporter complex
GABAergic synapse
distal axon
10%
15%
20%
Gene ratio
DEGs
4
6
8
10
12
− log10(FDR)
4
3
2
D
Δtranscript−positive nuclei
40
20
0
Fgf13
Ftx
Tenm1
Txlng
Parm1
Kcnmb2
Erc2
Cacnb4
1
2
Day−21 PEP2 CCI vs control log2FC
E
Fgf13
Kcnmb2
D21−D7 shift = +32.6 pp
D21−D7 shift = +15.6 pp
CCI − control SCT−corrected detection
(percentage points)
50
40
30
20
10
0
Day 7
Day 21
Day 7
Day 21
F
Fgf13
Kcnmb2
1.5
1.0
0.5
0.0
-0.5
1.50
0.39
0.21
-0.42
-0.44
-0.85
Acute
Chronic
1.0
0.5
0.0
-0.5
0.91
0.15
-0.25
-0.22
-0.31
-0.90
Acute
Chronic
TG CCI
TG Axotomy
DRG ScNT
adj.p < 0.05
ns

**Appendix Figure 7. Functional enrichment, candidate prioritization, and robustness of chronic PEP2 transcriptional remodeling after IoN-CCI (related to Figure 5). (A)** Equal-subtype sensitivity analysis of the six functional programs shown in Figure 5A. Day 21 CCI nuclei from PEP2, NF, NP, PEP1, and p-cLTMR2 neurons were repeatedly downsampled to equal subtype sizes (n = 148 nuclei per subtype; 1,000 iterations). Points indicate the median difference between the PEP2 median module score and the equal-weighted mean of the four comparator-subtype medians, and whiskers indicate the 2.5th–97.5th percentile range across subsampling iterations. Values at right indicate the percentage of iterations in which the PEP2 score exceeded the balanced comparator. PEP2 showed higher scores for all six programs in 100% of iterations. **(B)** Pairwise equal-n sensitivity analysis of PEP2 functional-program enrichment relative to individual canonical neuronal subtypes. PEP2 nuclei and each comparator subtype were repeatedly downsampled to equal group sizes across 1,000 iterations. Tiles show the median difference in module score between PEP2 and the indicated comparator subtype, with positive values indicating higher scores in PEP2; tile color reflects the magnitude of this difference. PEP2 exceeded NF, NP, PEP1, and p-cLTMR2 neurons for all six functional programs in 100% of pairwise subsampling iterations. **(C)** Unbiased functional enrichment of genes significantly upregulated in Day 21 PEP2 neurons after IoN-CCI. Gene Ontology terms are shown across biological process, molecular function, and cellular component categories. Gene ratio represents the proportion of input DEGs associated with each term, point size indicates the number of contributing DEGs, and color denotes −log10 false-discovery rate (FDR). Enriched terms included synaptic signaling, ion-channel regulation, calcium-channel activity, and axonal processes. **(D)** Candidate-eligible upregulated genes (adjusted $P < 0.05$ and log2FC > 0.5) are plotted according to average log2 fold change in CCI relative to control and the corresponding change in the percentage of transcript-positive nuclei. *Fgf13* and *Kcnmb2*, selected for independent RNAscope validation, are highlighted in red and blue, respectively. Open circles identify genes for which no other analyzed canonical neuronal subtype met the same differential-expression threshold. **(E)** Equal-n subsampling analysis of *Fgf13* and *Kcnmb2* transcript detection in PEP2 neurons at Day 7 and Day 21. Points indicate the mean CCI–control difference in the percentage of transcript-positive nuclei across 5,000 equal-n subsampling iterations, and whiskers indicate the 2.5th–97.5th percentile sampling range. Day 7 effects were small and their sampling ranges overlapped zero, whereas Day 21 effects remained positive for both genes after equalizing recovered nucleus numbers. Annotated values indicate the observed Day 21-minus-Day 7 difference in CCI–control transcript detection. **(F)** Cross-model targeted differential-expression comparison of *Fgf13* and *Kcnmb2* in PEP2 neurons across trigeminal ganglion CCI, trigeminal ganglion axotomy, and dorsal root ganglion sciatic nerve transection models. Line plots show log2 fold change relative to model-specific controls at acute and chronic injury phases. Filled points indicate adjusted $p < 0.05$ and $|\log2FC| \geq 0.5$, and open points indicate non-significant comparisons.

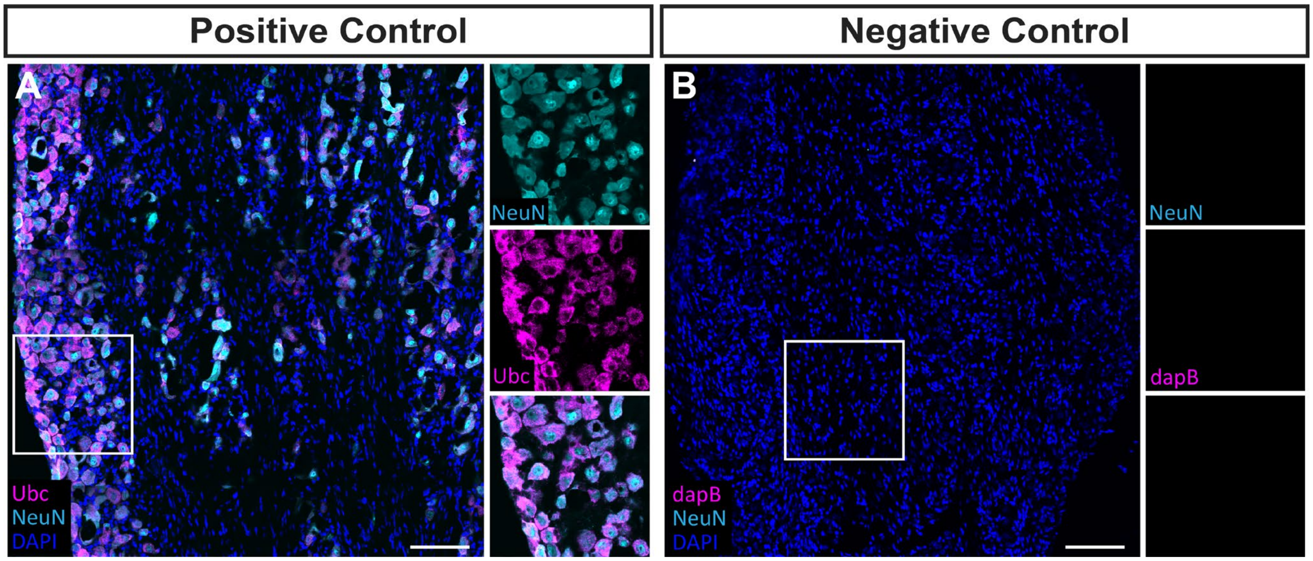


**Appendix Figure 8. RNAscope positive and negative control validation in mouse trigeminal ganglion tissue. (A)** Representative positive-control image showing detection of the housekeeping transcript Ubc using the RNAscope positive-control probe in channel 3 with TSA Vivid Fluorophore 650. Neuronal nuclei/cell bodies were labeled with rabbit anti-NeuN followed by goat anti-rabbit 488, and nuclei were counterstained with DAPI. The boxed region is shown at higher magnification in single-channel and merged insets to demonstrate robust Ubc signal in NeuN-positive cells. **(B)** Representative negative-control image showing minimal background signal following hybridization with the bacterial dapB negative-control probe in channel 1 with TSA Vivid Fluorophore 650. NeuN immunostaining was processed in parallel using the secondary antibody control condition, in which goat anti-rabbit 488 was applied without rabbit anti-NeuN primary antibody. The boxed region is shown at higher magnification in single-channel insets, demonstrating minimal dapB and secondary-only background signal. DAPI marks nuclei. Scale bars, 100 μm.

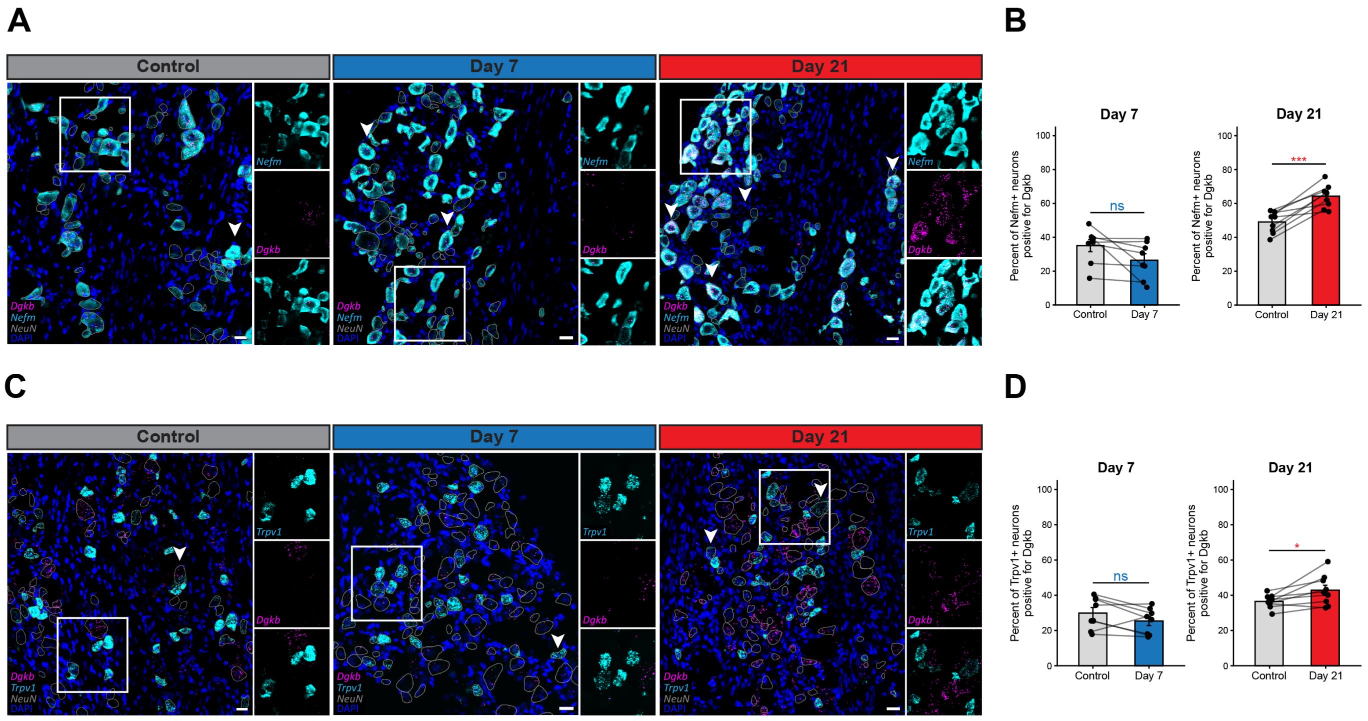


**Appendix Figure 9. RNAscope validation of broadly regulated Dgkb expression in distinct trigeminal sensory neuron populations after IoN-CCI.** *Dgkb* was selected for validation because its expression increased across multiple neuronal subtypes at Day 21 in the snRNA-seq analysis. **(A)** Representative multiplex RNAscope images showing *Dgkb* expression (magenta) in *Nefm*-positive neurons (cyan) from control, Day 7 IoN-CCI, and Day 21 IoN-CCI trigeminal ganglia. *Nefm* marks NF-associated neurons. Arrowheads indicate representative Nefm-positive neurons, and boxed regions are shown as enlarged channel-separated insets. **(B)** Percentage of *Nefm*-positive neurons positive for *Dgkb* at Day 7 and Day 21. No significant CCI-associated difference in *Dgkb* expression was detected at Day 7, whereas expression was significantly increased at Day 21. **(C)** Representative multiplex RNAscope images showing *Dgkb* expression (magenta) in *Trpv1*-positive neurons (cyan) from control, Day 7 IoN-CCI, and Day 21 IoN-CCI trigeminal ganglia. *Trpv1* marks nociceptor-associated neurons. Arrowheads indicate representative *Trpv1*-positive neurons, and boxed regions are shown as enlarged channel-separated insets. **(D)** Percentage of *Trpv1*-positive neurons positive for *Dgkb* at Day 7 and Day 21. *Dgkb* was unchanged at Day 7 but modestly increased at Day 21 after IoN-CCI. NeuN immunolabeling was used to define neuronal somata and is shown as cell-body outlines to preserve RNAscope puncta. For B and D, n = 8 mice at Day 7 and n = 9 mice at Day 21; one whole-section value per condition was analyzed for each mouse. Data are mean ± SEM, with paired points indicating matched samples. Matched control and IoN-CCI values within each timepoint were compared using paired two-tailed t tests.  Scale bars, 20 µm; images were acquired at 40× magnification. ns, not significant; $*P < 0.05$; $***P < 0.001$.

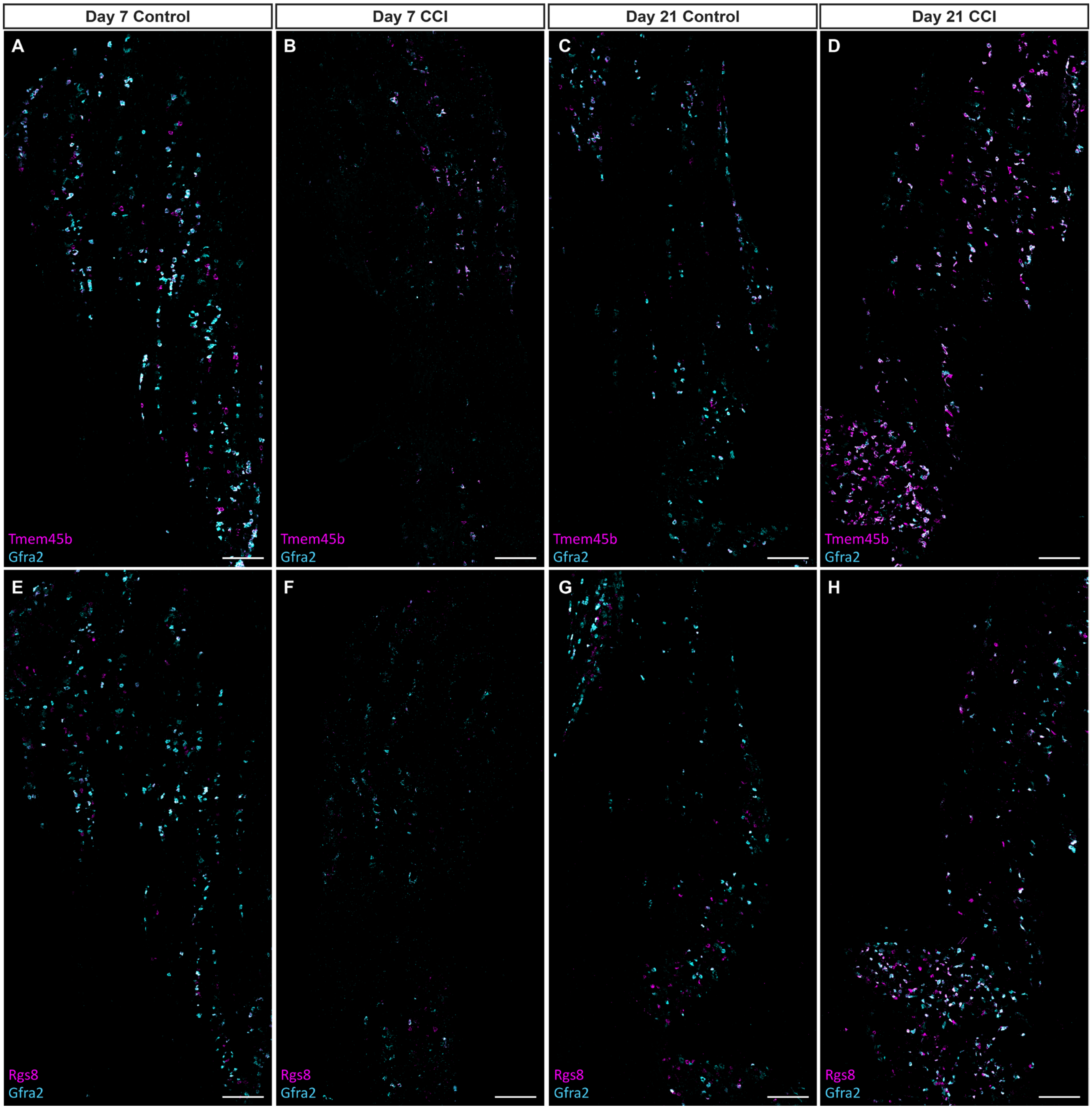


**Appendix Figure 10. Full trigeminal ganglion RNAscope images for p-cLTMR2 (related to Figure 4).** Representative tiled multiplex RNAscope images of trigeminal ganglion (TG) sections from Day 7 contralateral control, Day 7 IoN-CCI, Day 21 contralateral control, and Day 21 IoN-CCI conditions. **(A–D)** *Tmem45b* (magenta) and *Gfra2* (cyan) expression across the TG at Day 7 and Day 21. **(E–H)** *Rgs8* (magenta) and *Gfra2* (cyan) expression across the TG at Day 7 and Day 21. *Gfra2* marks p-cLTMR2-enriched neurons. Scale bars, 200 μm.

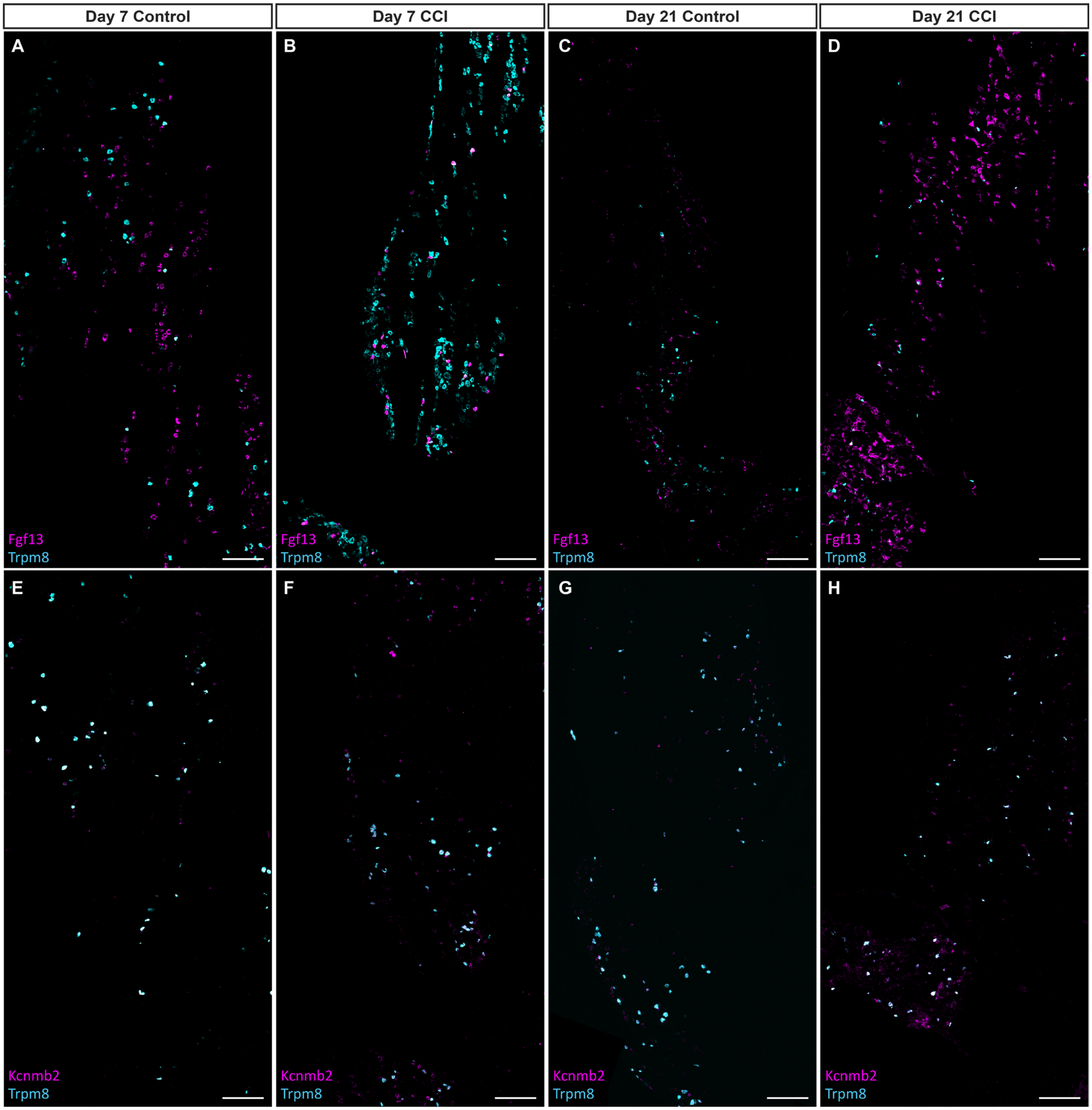


**Appendix Figure 11. Full trigeminal ganglion RNAscope images for PEP2 (related to figure 5).** Representative tiled multiplex RNAscope images of trigeminal ganglion (TG) sections from Day 7 contralateral control, Day 7 IoN-CCI, Day 21 contralateral control, and Day 21 IoN-CCI conditions. **(A–D)** *Fgf13* (magenta) and *Trpm8* (cyan) expression across the TG at Day 7 and Day 21. **(E–H)** *Kcnmb2* (magenta) and *Trpm8* (cyan) expression across the TG at Day 7 and Day 21. *Trpm8* marks PEP2-enriched neurons. Scale bars, 200 µm.

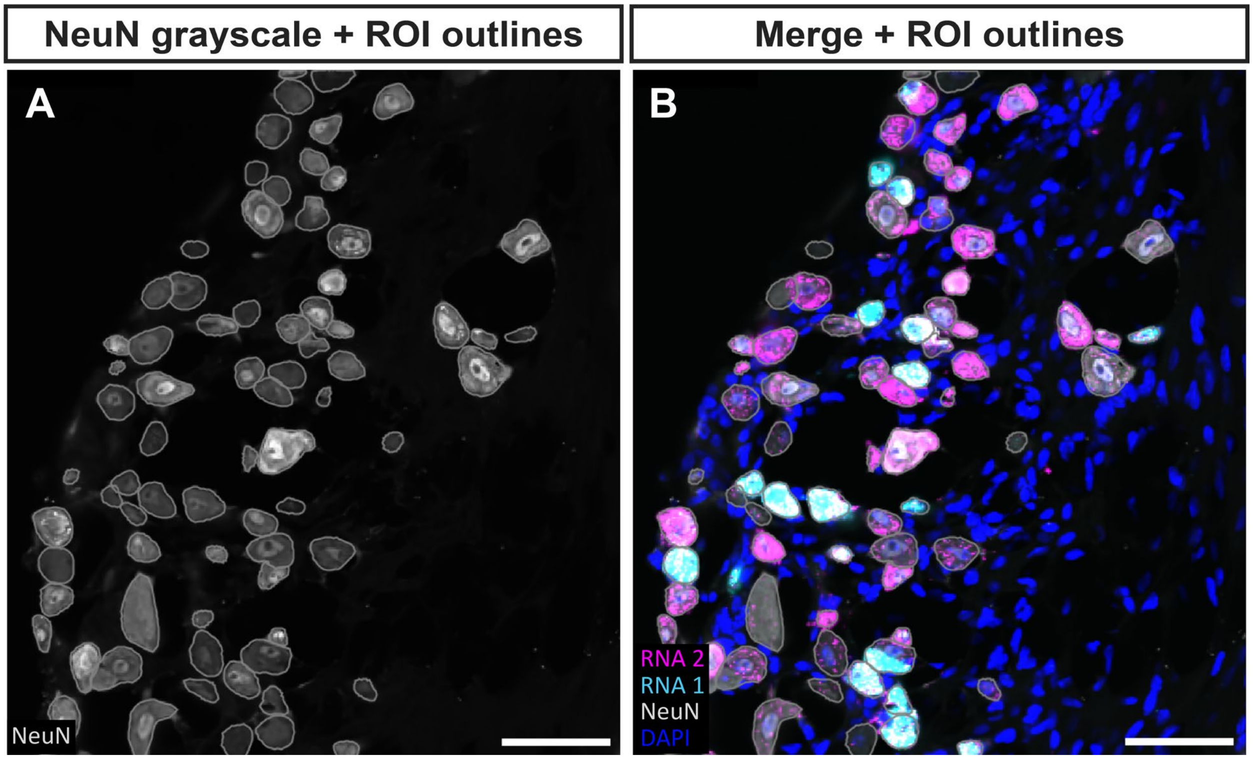


**Appendix Figure 12. NeuN-guided neuronal outlines used for RNAscope image presentation. (A)** NeuN immunofluorescence channel used to identify and manually trace neuronal cell bodies. **(B)** Corresponding merged image showing DAPI, RNA1, and RNA2 signals with the NeuN-guided neuronal outlines overlaid. The NeuN signal was displayed as an outline rather than as an additional fluorescence channel to avoid obscuring or introducing a competing color into the RNAscope probe signals. These outlines were used for figure visualization only and were not used for quantitative image analysis. Scale bars, 50 μm.

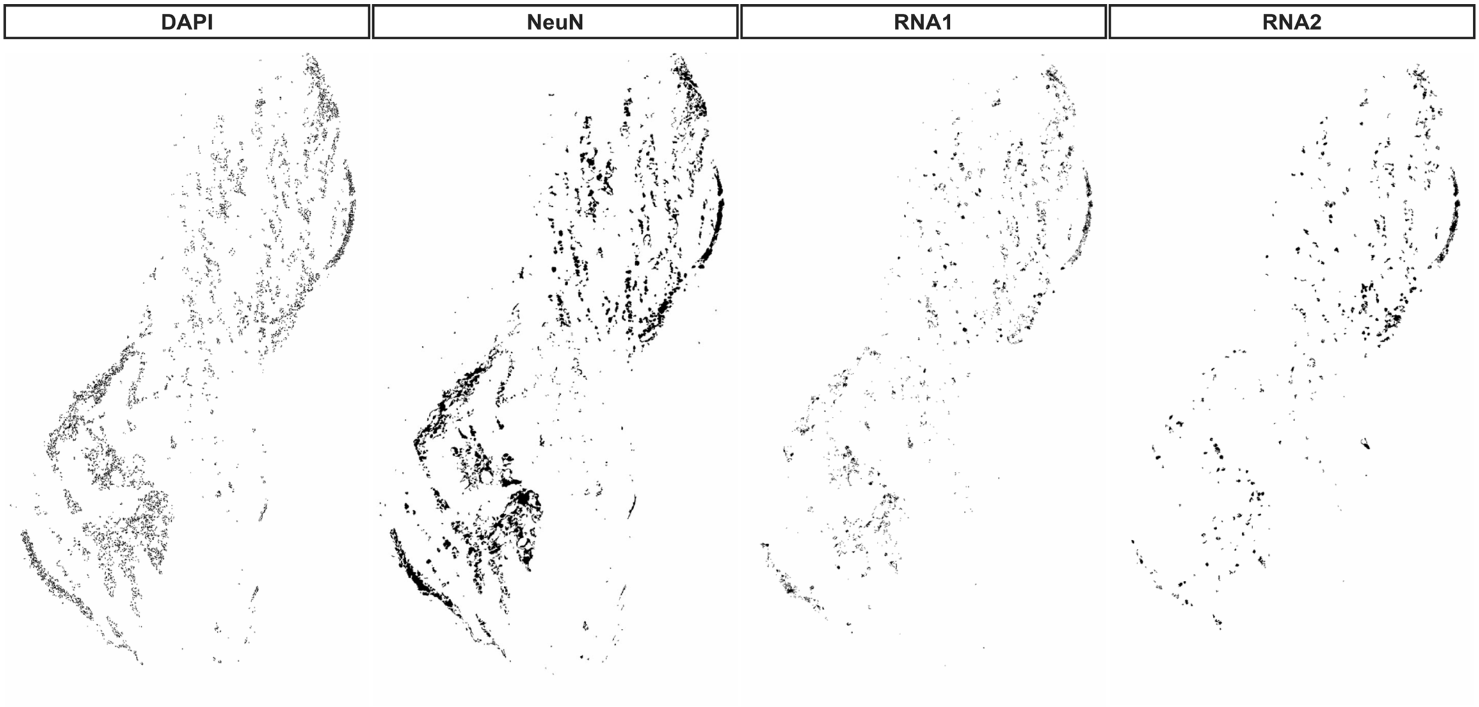


**Appendix Figure 13. Representative binary masks generated during RNAscope image analysis of trigeminal ganglion sections. (A)** DAPI mask used to identify nuclei. **(B)** NeuN mask used to restrict DAPI-derived nuclear ROI selection to neuronal regions. **(C)** RNA1 mask corresponding to the neuronal subtype-marker probe. **(D)** RNA2 mask corresponding to the candidate-gene probe. The same image-processing and thresholding workflow was applied across experimental conditions and timepoints for RNAscope quantification.

**Appendix**
**Table 1**
Materials

| Category | Item / Material | Details (Catalog #, Manufacturer, Version) | RRIDs |
|---|---|---|---|
| **Animals** | C57BL/6J wild-type mice | 8–10 weeks old, both sexes | RRID: IMSR_JAX:000664 |
| **Anesthesia & Analgesia** | Ketamine/xylazine cocktail | For surgical anesthesia | — |
| | Ophthalmic ointment | To prevent corneal drying | — |
| | Ketoprofen (5 mg/kg) | Pre- and post-operative analgesia | — |
| **Surgical Materials** | Surgical microscope | For infraorbital nerve exposure | — |
| | 7-0 silk suture | Nerve ligation | — |
| | Surgical glue | For skin closure | — |
| | Glass bead sterilizer | 250 °C, 15 s between animals | — |
| | Sterile drapes, gloves, gauze, cotton swabs | Standard surgical consumables | — |
| | Heating pad | To maintain body temperature during surgery | — |
| **Animal Care Post-surgery** | Hydrogel packs | ClearH2O HydroGel™ | — |
| | Soft dough diet | Provided on cage floor for 7 days | — |
| **Tissue Handling & Processing** | Sucrose (Molecular Biology Grade) | Sigma-Aldrich, Cat. #S0389 | — |
| | HEPES Buffer (1 M, pH 8.0) | Thermo Fisher Scientific, Cat. #15630080 | — |
| | Calcium Chloride ($CaCl_2$) | Sigma-Aldrich, Cat. #C5670 | — |
| | Magnesium Acetate | Sigma-Aldrich, Cat. #M5661 | — |
| | EDTA (0.5 M, pH 8.0) | Thermo Fisher Scientific, Cat. #15575020 | — |
| | Dithiothreitol (DTT) | Thermo Fisher Scientific, Cat. #R0861 | — |
| | NP-40 Substitute (IGEPAL CA-630) | Sigma-Aldrich, Cat. #I8896 | — |
| | Bovine Serum Albumin (BSA), molecular biology grade | Sigma-Aldrich, Cat. #A9647 | — |
| | SUPERase•In RNase Inhibitor | Thermo Fisher Scientific, Cat. #AM2696 | — |
| | 40 µm Cell Strainer | Thermo Fisher Scientific, Cat. #08-771-1 | — |
| | 30 µm Cell Strainer | Miltenyi Biotec, Cat. #130-041-407 | — |
| | Dounce Homogenizer | Fisher Scientific, Cat. #357538 | — |
| | Sucrose Cushion Solution (1.8 M) | Sigma-Aldrich, Cat. #NUC-201 | — |
| | DRAQ5 Nuclear Stain | Thermo Fisher Scientific, Cat. #62251 | — |
| | SuperFrost Plus slides | Thermo Fisher, Cat. #12-550-15 | — |
| | 4% Paraformaldehyde (PFA) in PBS | Fixative | — |
| | Ethanol (50%, 70%, 100%) | For dehydration | — |
| | 5× SSC buffer | For RNAscope probe preservation | — |
| | Hydrophobic barrier pen | ImmEdge, Vector Labs | — |
| **Single-nucleus RNA-seq** | 10x Genomics Chromium Controller | Single-nucleus library preparation | RRID:SCR_019326 |
| | 10x Genomics Chromium Next GEM Single Cell 3' Reagent Kit v3.1 | 10x Genomics | — |
| | 10x Genomics Cell Ranger | Pipeline, count function | RRID:SCR_017344 |
| | STAR aligner | Read alignment | RRID:SCR_004463 |
| | Illumina NovaSeq 6000 | Illumina | RRID:SCR_016387 |
| | Qubit 4 Fluorometer | Thermo Fisher Scientific, Cat. #Q33226 | RRID:SCR_018095 |
| | Agilent Bioanalyzer | Agilent Technologies, Cat. #G2939BA | RRID:SCR_018043 |
| | Sony SH800 Sorter | BD Biosciences / Sony Biotechnology | RRID:SCR_018066 |
| | Low-bind Microcentrifuge Tubes | Sorenson BioScience, Cat. #11700 | — |
| | Refrigerated Microcentrifuge | Eppendorf 5418R | RRID:SCR_018050 |
| | R v4.4.3 | Data analysis | RRID:SCR_001905 |
| | Seurat v5 | Data analysis | RRID:SCR_016341 |
| | clusterProfiler | GO/KEGG enrichment | RRID:SCR_016884 |
| | Harmony (R package) | Version 0.1 or later, https://github.com/immunogenomics/harmony | RRID:SCR_022206 |
| **RNAscope ISH** | Multiplex Fluorescent v2 Kit | Advanced Cell Diagnostics (ACD) | — |

| | | | |
|---|---|---|---|
| | HybEZ Oven | Cat. #321711, ACD | — |
| | HybEZ Humidity Control Tray | Cat. #310012, ACD | — |
| | EZ-Batch™ Slide Holder | Cat. #321716, ACD | — |
| | Multiplex Fluorescent Detection Reagents v2 | Cat. #323110, ACD | — |
| | Protease III Reagent | Cat. #322381, ACD | — |
| | Hydrogen Peroxide | Cat. #322381, ACD | — |
| | Dgkb probe | Cat. #1826041-C1, ACD | — |
| | Tmem45b probe | Cat. #420461, ACD | — |
| | Trpm8 probe | Cat. #420451, ACD | — |
| | Trpv1 probe | Cat. #313331-C3, ACD | — |
| | Nefm probe | Cat. #315611-C2, ACD | — |
| | Fgf13 probe | Cat. #1303051-C2, ACD | — |
| | Gfra2 probe | Cat. #441481-C2, ACD | — |
| | Rgs8 probe | Cat. #515041-C3, ACD | — |
| | Kcnmb2 probe | Cat. #539871-C3, ACD | — |
| | Ubc probe (positive control) | Cat. #310771-C3, ACD | — |
| | 3-plex Negative Control probe | Cat. #320871, ACD | — |
| | TSA Vivid™ 570 | Cat. #323272, ACD | — |
| | TSA Vivid™ 650 | Cat. #323273, ACD | — |
| | Akoya Diluent | For TSA fluorophore dilution | — |
| **Immunofluorescence (Antibodies)** | NeuN | Rabbit monoclonal IgG, Abcam (ab177487), 1:400 | RRID: AB_2532109 |
| | Goat anti-Rabbit IgG (H+L) Alexa Fluor 488 | Invitrogen, conjugated secondary | RRID: AB_143165 |
| | Goat anti-Rabbit IgG (H+L) Alexa Fluor 555 | Invitrogen, conjugated secondary | RRID: AB_2535849 |
| | Goat anti-Rabbit IgG (H+L) Alexa Fluor 647 | Invitrogen, conjugated secondary | RRID: AB_2535812 |
| | Anti-NeuN (Rabbit polyclonal) | Millipore, Cat. #ABN78, 1:2000–1:5000 | RRID: AB_10807945 |
| | Goat anti-Rabbit IgG (FITC or Cy3 conjugated) | Jackson ImmunoResearch, Cat. #111-095-003 / #111-165-003 | RRID: AB_2337972 |
| **Buffers & Reagents** | 1× PBS | Crystalgen Ultrapure, Ref. 221-132-10 | — |
| | Normal Goat Serum (NGS) | Sigma-Aldrich, G9023 | — |
| | Triton X-100 | BioUltra, 93443 | — |
| | DAPI solution | Thermo Fisher, Cat. #62248 | — |
| **Microscopy** | Nikon A1R HD Confocal | — | RRID:SCR_020317 |
| | Immersion Oil (Type F) | Nikon | — |
| | Coverslips (No. 1.5H) | Thickness-matched | — |
| | Nikon Elements (JOBS Explorer, Tiling, Stitching, MIP) | Image acquisition & analysis | RRID:SCR_014329 |
| **Image Analysis & Statistics** | Fiji | Custom macro for segmentation and quantification | RRID:SCR_002285 |
| | Excel (VBA macro: ImportCSVAddCols) | Data merging and organization | — |
| | R v4.4.3 | Statistical analysis and visualization | RRID:SCR_001905 |

**Appendix Table 2:** IoN-CCI Day 7 top significant neuronal DEGs. Top 100 per subtype; adjusted p-value < 0.05; |log2FC| > 0.5. Injured Day 7 comparison is cluster 20 vs canonical neurons within CCI.

NF

| Gene Name | adjusted p-value | log2FoldChange | Gene Name | adjusted p-value | log2FoldChange |
|---|---|---|---|---|---|
| Camk1d | 5.58e-246 | -2.324 | Gm36264 | 7.58e-23 | 6.860 |
| Bc1 | 1.24e-172 | 5.593 | Gm765 | 1.47e-22 | -2.660 |
| Hexb | 2.23e-152 | -4.166 | Arglu1 | 1.62e-22 | -0.698 |
| Xlr5a | 2.25e-109 | -9.157 | Dst | 2.60e-22 | -0.552 |
| Sec61a1 | 1.36e-97 | -5.256 | Clk1 | 6.94e-22 | 1.452 |
| Taco1 | 2.98e-96 | -3.484 | Rsrp1 | 3.31e-21 | 0.863 |
| Snhg11 | 1.12e-79 | 0.975 | Ptgds | 3.94e-21 | 3.767 |
| Cmss1 | 8.26e-77 | -1.323 | Pkd1 | 6.72e-20 | 0.940 |
| Gm19951 | 6.17e-76 | 2.089 | Slx1b | 8.39e-20 | -2.829 |
| Lars2 | 1.93e-71 | -1.620 | Polr1b | 1.06e-19 | -2.623 |
| 5830416I19Rik | 2.94e-70 | -8.277 | Rusc1 | 2.06e-19 | 1.321 |
| Pde1c | 2.45e-69 | -1.843 | Ddx5 | 2.52e-19 | 1.093 |
| Grik1 | 2.02e-67 | -2.311 | Ptprs | 3.23e-19 | 0.819 |
| Peak1 | 9.05e-67 | 2.032 | Pcdh15 | 3.34e-19 | -1.677 |
| AY036118 | 2.60e-64 | 1.641 | Gm11337 | 3.79e-18 | -6.140 |
| Meg3 | 4.53e-64 | 0.718 | Acoxl | 5.70e-18 | -4.715 |
| Gm12216 | 4.95e-59 | -4.616 | Ano2 | 8.93e-18 | -4.320 |
| Klhl14 | 8.19e-52 | -7.679 | Dgkz | 1.05e-17 | 0.868 |
| 1700054A03Rik | 1.92e-50 | -7.983 | Arhgef25 | 1.63e-17 | 1.492 |
| Fbxl7 | 8.04e-45 | -2.841 | Map1a | 2.58e-17 | -0.982 |
| Sema3c | 3.58e-44 | -3.434 | Sil1 | 2.98e-17 | -1.694 |
| Mc1r | 2.87e-42 | -6.359 | Syne1 | 3.71e-17 | -0.679 |
| Kcnq2 | 2.99e-39 | 1.217 | mt-Co3 | 1.04e-16 | 0.737 |
| Gm50306 | 6.75e-39 | -7.168 | Wdr72 | 1.06e-16 | -3.055 |
| 8430422H06Rik | 9.14e-39 | -5.700 | 5031439G07Rik | 1.19e-16 | 1.440 |
| Pde6a | 3.81e-38 | -5.616 | Cdk10 | 1.38e-16 | 1.492 |
| Gm42418 | 7.79e-38 | -0.929 | Ms4a1 | 3.55e-16 | -5.094 |
| Gm10062 | 6.87e-37 | -7.047 | Prph | 4.24e-16 | 1.047 |
| Igf2bp3 | 2.09e-34 | -6.106 | Ninj2 | 4.75e-16 | -4.643 |
| Gm26917 | 6.12e-34 | -1.395 | Vit | 1.73e-15 | -2.572 |
| Lrp1b | 2.86e-33 | -0.843 | Cmpk1 | 1.47e-14 | -2.411 |
| Adamtsl1 | 8.65e-33 | -3.808 | Sh3glb2 | 3.50e-14 | 1.496 |
| 1700126G02Rik | 1.99e-32 | -5.861 | Gm14326 | 3.93e-14 | 2.734 |
| Il27 | 9.25e-32 | -5.468 | Atp6v0a1 | 4.12e-14 | 0.703 |
| Gh | 1.66e-31 | 5.235 | Rgs11 | 4.25e-14 | 1.377 |

| Gene Name | adjusted p-value | log2FoldChange | Gene Name | adjusted p-value | log2FoldChange |
|---|---|---|---|---|---|
| Shb | 2.13e-31 | -3.550 | Mylip | 8.57e-14 | -4.106 |
| Gm15564 | 1.12e-30 | -4.707 | Tnrc6a | 8.99e-14 | 0.618 |
| Chl1 | 2.15e-30 | -2.640 | Rbm39 | 1.18e-13 | 0.755 |
| Cdk8 | 7.62e-30 | -1.019 | Npas3 | 2.55e-13 | -1.819 |
| Nckap5 | 4.65e-29 | -3.165 | Fyb2 | 2.56e-13 | -3.462 |
| Gm28307 | 3.58e-28 | -6.640 | 4930441J16Rik | 3.77e-13 | -5.600 |
| Naaladl2 | 5.53e-28 | -1.439 | Gm17529 | 3.77e-13 | -5.695 |
| Gm48099 | 1.98e-26 | -2.619 | 4930511M06Rik | 5.42e-13 | -3.725 |
| Cdh23 | 3.31e-26 | -4.134 | 4930558K02Rik | 6.17e-13 | -5.583 |
| Pcbp4 | 2.47e-25 | 2.028 | Ctcflos | 9.24e-13 | -4.854 |
| Ntm | 3.08e-25 | -1.357 | Fsip1 | 1.01e-12 | -5.566 |
| Zfp994 | 6.33e-25 | -4.784 | Gm47371 | 1.65e-12 | -5.648 |
| Gigyf1 | 1.16e-23 | 1.520 | Hba-a1 | 2.06e-12 | 4.504 |
| Asxl3 | 4.11e-23 | -2.425 | Rian | 2.59e-12 | 0.619 |
| Prl | 7.58e-23 | 7.475 | Lmtk3 | 2.78e-12 | 1.505 |

NP

| Gene Name | adjusted p-value | log2FoldChange | Gene Name | adjusted p-value | log2FoldChange |
|---|---|---|---|---|---|
| Camk1d | 1.20e-57 | -2.140 | Pcdh15 | 3.28e-03 | -2.313 |
| Bc1 | 7.53e-34 | 5.769 | Pcbp4 | 3.79e-03 | 1.673 |
| Hexb | 1.40e-28 | -3.281 | Trim2 | 3.91e-03 | -1.340 |
| Gm19951 | 1.34e-24 | 2.202 | Adamtsl1 | 5.34e-03 | -2.966 |
| Xlr5a | 2.03e-24 | -5.500 | Zcchc18 | 5.70e-03 | 2.103 |
| Sec61a1 | 4.31e-20 | -4.991 | Gm11337 | 6.52e-03 | -4.108 |
| Snhg11 | 3.95e-19 | 1.086 | Arhgap33 | 6.87e-03 | 2.133 |
| Taco1 | 5.02e-18 | -3.140 | Arglu1 | 7.47e-03 | -0.871 |
| 5830416I19Rik | 2.12e-17 | -6.298 | mt-Co2 | 7.55e-03 | 0.596 |
| AY036118 | 3.37e-16 | 1.586 | Stub1 | 9.66e-03 | 2.785 |
| Lrp1b | 4.11e-14 | -1.545 | Ms4a1 | 1.09e-02 | -4.196 |
| 1700054A03Rik | 2.45e-12 | -5.809 | Gm15564 | 1.20e-02 | -3.331 |
| Klhl14 | 8.80e-12 | -5.693 | Tmem108 | 1.55e-02 | -3.600 |
| Gm12216 | 1.10e-11 | -5.097 | AC149090.1 | 2.16e-02 | -0.693 |
| Cmss1 | 7.29e-11 | -0.940 | Wdyhv1 | 2.28e-02 | -3.108 |
| Peak1 | 9.67e-11 | 2.178 | Ddx5 | 2.28e-02 | 1.001 |

| Gene Name | adjusted p-value | log2FoldChange | Gene Name | adjusted p-value | log2FoldChange |
|---|---|---|---|---|---|
| Lars2 | 1.17e-10 | -1.263 | Pcyt2 | 2.44e-02 | 1.486 |
| Prph | 1.22e-09 | 1.390 | 6030443J06Rik | 2.87e-02 | -2.008 |
| 8430422H06Rik | 7.30e-09 | -5.085 | Vwc2l | 3.00e-02 | -4.062 |
| Mc1r | 2.35e-08 | -5.196 | Grina | 3.41e-02 | 1.987 |
| Dgkz | 2.64e-08 | 0.894 | Dst | 3.99e-02 | -0.521 |
| Pde1c | 7.46e-08 | -1.526 | Naaladl2 | 4.14e-02 | -1.114 |
| Gm10062 | 7.48e-08 | -4.991 | Fam189b | 4.27e-02 | 1.576 |
| Gm50306 | 7.48e-08 | -5.298 | | | |
| Cdh23 | 4.13e-07 | -4.966 | | | |
| Ntm | 7.63e-07 | -2.196 | | | |
| Pde6a | 1.06e-06 | -4.356 | | | |
| Gm26917 | 1.41e-06 | -1.295 | | | |
| Sema3c | 2.13e-06 | -2.916 | | | |
| Wdr72 | 2.21e-06 | -4.890 | | | |
| Shb | 2.51e-06 | -4.031 | | | |
| Zfp994 | 1.15e-05 | -4.991 | | | |
| 1700126G02Rik | 1.68e-05 | -4.337 | | | |
| Il27 | 1.95e-05 | -4.196 | | | |
| Gm42418 | 3.24e-05 | -0.706 | | | |
| Rsrp1 | 3.25e-05 | 1.023 | | | |
| Acoxl | 3.41e-05 | -4.693 | | | |
| Epha6 | 9.47e-05 | -1.881 | | | |
| Slx1b | 1.31e-04 | -3.015 | | | |
| Csmd3 | 1.59e-04 | -2.874 | | | |
| Ano2 | 1.69e-04 | -4.394 | | | |
| Gm28307 | 2.88e-04 | -4.466 | | | |
| Igf2bp3 | 4.78e-04 | -3.809 | | | |
| 4930511M06Rik | 4.87e-04 | -4.500 | | | |
| Gh | 5.87e-04 | 3.657 | | | |
| Gm48099 | 7.57e-04 | -2.956 | | | |
| Hspa14 | 8.11e-04 | -3.752 | | | |
| D430042O09Rik | 8.58e-04 | -2.510 | | | |
| Clk1 | 1.33e-03 | 1.460 | | | |
| Kcnt1 | 1.54e-03 | 0.913 | | | |

SST

| Gene Name | adjusted p-value | log2FoldChange | Gene Name | adjusted p-value | log2FoldChange |
|---|---|---|---|---|---|
| Camk1d | 4.67e-20 | -2.189 | | | |
| Bc1 | 3.49e-17 | 4.605 | | | |
| Hexb | 1.69e-10 | -4.077 | | | |
| Cmss1 | 1.84e-09 | -1.455 | | | |
| Gm36264 | 2.37e-08 | 6.182 | | | |
| Taco1 | 1.23e-07 | -4.096 | | | |
| Lrp1b | 1.60e-06 | -3.070 | | | |
| Xlr5a | 2.14e-06 | -5.240 | | | |
| Sec61a1 | 2.56e-05 | -3.874 | | | |
| Gm19951 | 3.28e-05 | 2.144 | | | |
| Grik1 | 7.79e-05 | -3.520 | | | |
| Sema3c | 2.22e-04 | -3.874 | | | |
| Gm26917 | 2.84e-04 | -1.270 | | | |
| 5830416I19Rik | 4.00e-04 | -4.775 | | | |
| Csmd3 | 2.26e-03 | -3.983 | | | |
| Gm42418 | 4.27e-03 | -0.842 | | | |
| Ddx5 | 4.79e-03 | 1.169 | | | |
| 1700054A03Rik | 6.77e-03 | -4.458 | | | |
| Ivns1abp | 6.86e-03 | 1.338 | | | |
| Snhg11 | 7.28e-03 | 0.589 | | | |

PEP1

| Gene Name | adjusted p-value | log2FoldChange | Gene Name | adjusted p-value | log2FoldChange |
|---|---|---|---|---|---|
| Camk1d | 8.31e-98 | -2.436 | Fbxl7 | 9.09e-09 | -1.506 |
| Bc1 | 4.02e-69 | 5.574 | Gm36264 | 1.09e-08 | 4.804 |
| Sec61a1 | 5.45e-54 | -6.177 | Prph | 1.94e-08 | 1.020 |
| Hexb | 1.25e-51 | -4.183 | Ptgds | 3.78e-08 | 3.882 |
| Xlr5a | 3.28e-51 | -8.332 | Zfp994 | 4.78e-08 | -3.792 |
| Taco1 | 5.73e-42 | -4.454 | Dnah11 | 4.89e-08 | -1.214 |
| 5830416I19Rik | 4.33e-36 | -7.536 | Asxl3 | 9.35e-07 | -2.023 |
| Snhg11 | 3.52e-35 | 1.001 | Gm47371 | 1.06e-06 | -5.147 |
| Cmss1 | 1.23e-31 | -1.626 | Sil1 | 1.26e-06 | -1.749 |
| 1700054A03Rik | 4.26e-27 | -7.010 | Ano2 | 1.35e-06 | -1.688 |
| Grik1 | 7.20e-24 | -2.557 | mt-Co1 | 1.37e-06 | 1.082 |

| Gene Name | adjusted p-value | log2FoldChange | Gene Name | adjusted p-value | log2FoldChange |
|---|---|---|---|---|---|
| Lars2 | 8.03e-24 | -1.818 | Gm14326 | 2.05e-06 | 3.249 |
| Gm12216 | 1.35e-23 | -4.826 | Adamtsl1 | 3.14e-06 | -2.688 |
| Pde1c | 1.73e-22 | -1.472 | H2-K1 | 3.17e-06 | 2.234 |
| Lrp1b | 8.44e-22 | -1.239 | Gm11337 | 3.33e-06 | -4.890 |
| Meg3 | 2.90e-21 | 0.729 | A930016O22Rik | 3.86e-06 | -1.536 |
| Klhl14 | 2.06e-20 | -5.936 | Dazap1 | 6.95e-06 | 1.583 |
| Sema3c | 1.83e-19 | -3.951 | Chsy3 | 9.20e-06 | -2.599 |
| Shb | 2.44e-18 | -4.809 | Cmpk1 | 1.21e-05 | -2.589 |
| Gm19951 | 3.95e-18 | 1.695 | Chl1 | 1.58e-05 | -1.320 |
| Ntm | 4.79e-17 | -2.514 | Col5a3 | 2.10e-05 | 1.400 |
| Il27 | 1.21e-16 | -4.900 | Rubcnl | 3.23e-05 | -4.576 |
| Mc1r | 6.12e-16 | -6.224 | Dctd | 3.23e-05 | -4.688 |
| Gm42418 | 2.70e-15 | -1.181 | Rab29 | 3.23e-05 | -4.921 |
| Gm26917 | 1.26e-14 | -1.798 | Rusc1 | 5.23e-05 | 1.282 |
| mt-Co3 | 1.85e-14 | 0.995 | 4930558K02Rik | 5.68e-05 | -4.792 |
| Nckap5 | 2.46e-14 | -3.807 | Prl | 8.55e-05 | 5.219 |
| Ddx5 | 8.78e-14 | 1.248 | AC149090.1 | 1.06e-04 | -0.767 |
| Gm50306 | 9.92e-14 | -6.039 | Ninj2 | 1.19e-04 | -4.224 |
| AY036118 | 1.34e-13 | 1.335 | Epha6 | 1.32e-04 | -1.264 |
| 8430422H06Rik | 2.41e-13 | -4.770 | Slc9b1 | 1.75e-04 | -4.411 |
| 1700126G02Rik | 6.44e-13 | -5.905 | Rian | 1.80e-04 | 0.725 |
| Cdh23 | 1.14e-12 | -5.134 | Csmd3 | 1.85e-04 | -1.619 |
| Gm10062 | 1.20e-12 | -5.936 | Kcnd2 | 2.31e-04 | -3.556 |
| Igf2bp3 | 2.22e-12 | -6.108 | Gm15564 | 2.37e-04 | -2.516 |
| Pde6a | 2.50e-12 | -4.161 | Atp9b | 2.47e-04 | -1.231 |
| Rsrp1 | 2.54e-12 | 1.080 | Tmem108 | 2.52e-04 | -1.745 |
| Slx1b | 2.40e-11 | -3.614 | Arhgef25 | 2.63e-04 | 1.439 |
| Gm28307 | 2.57e-11 | -5.595 | Gh | 3.05e-04 | 4.049 |
| Polr1b | 6.10e-11 | -3.799 | Agrn | 3.61e-04 | 1.360 |
| Pcbp4 | 8.35e-11 | 2.178 | Selenow | 4.19e-04 | 1.401 |
| mt-Atp6 | 1.99e-10 | 0.993 | Adam15 | 4.80e-04 | 1.708 |
| Wdr72 | 2.61e-10 | -4.874 | Syne1 | 5.04e-04 | -0.914 |
| Gm48099 | 3.22e-10 | -3.432 | Vit | 5.07e-04 | -2.504 |
| Acoxl | 9.49e-10 | -5.475 | Gm14582 | 5.35e-04 | -4.652 |
| Naaladl2 | 1.18e-09 | -1.406 | Gm38190 | 6.07e-04 | -2.951 |
| Cdk8 | 1.56e-09 | -1.198 | Rbm39 | 7.15e-04 | 0.630 |

| Gene Name | adjusted p-value | log2FoldChange | Gene Name | adjusted p-value | log2FoldChange |
|---|---|---|---|---|---|
| 4930511M06Rik | 3.11e-09 | -5.536 | Hbb-bs | 8.84e-04 | 3.804 |
| Peak1 | 5.29e-09 | 1.555 | Gm765 | 8.87e-04 | -1.394 |
| Clk1 | 6.47e-09 | 1.368 | Scn10a | 9.10e-04 | 0.795 |

PEP2

| Gene Name | adjusted p-value | log2FoldChange | Gene Name | adjusted p-value | log2FoldChange |
|---|---|---|---|---|---|
| Camk1d | 8.47e-91 | -2.455 | Acoxl | 1.52e-05 | -4.760 |
| Bc1 | 4.14e-55 | 4.904 | Cdh23 | 1.83e-05 | -4.136 |
| Xlr5a | 2.27e-36 | -7.780 | Adamtsl1 | 2.33e-05 | -2.852 |
| Taco1 | 1.57e-35 | -3.696 | Arhgef25 | 2.36e-05 | 1.699 |
| Grik1 | 2.57e-35 | -5.047 | Paxbp1 | 2.69e-05 | -1.726 |
| Snhg11 | 6.88e-34 | 0.976 | Nsmf | 3.80e-05 | 3.196 |
| Hexb | 1.10e-32 | -3.655 | Slx1b | 5.28e-05 | -2.914 |
| Lars2 | 2.95e-25 | -1.706 | Gigyf1 | 5.70e-05 | 1.594 |
| Sec61a1 | 4.86e-23 | -3.707 | Ddx5 | 6.39e-05 | 1.068 |
| Hjurp | 8.33e-23 | 1.466 | Fbxl7 | 7.39e-05 | -1.690 |
| Gm42418 | 1.10e-22 | -1.141 | Cmpk1 | 7.68e-05 | -3.010 |
| 5830416I19Rik | 2.76e-22 | -6.528 | Gm14326 | 8.40e-05 | 3.536 |
| Cmss1 | 3.87e-21 | -1.207 | Pcbp4 | 8.43e-05 | 1.885 |
| Gm26917 | 1.20e-18 | -1.540 | AC149090.1 | 1.04e-04 | -0.824 |
| Lrp1b | 1.55e-18 | -1.465 | Dgkz | 2.82e-04 | 0.774 |
| Meg3 | 1.65e-17 | 0.577 | Gm48099 | 4.58e-04 | -2.810 |
| Pde6a | 3.75e-17 | -6.043 | Diablo | 5.08e-04 | 1.646 |
| 1700054A03Rik | 1.96e-15 | -6.345 | Prl | 6.44e-04 | 4.908 |
| Klhl14 | 1.96e-15 | -6.251 | Ctcflos | 7.26e-04 | -4.551 |
| Gm19951 | 2.71e-15 | 1.962 | Nell1 | 7.74e-04 | -3.909 |
| Mc1r | 4.26e-14 | -5.873 | Gnaz | 8.42e-04 | -2.003 |
| Peak1 | 1.04e-13 | 1.795 | Syne1 | 9.41e-04 | -0.672 |
| Igf2bp3 | 1.93e-13 | -5.799 | Vit | 1.11e-03 | -2.482 |
| Il27 | 4.05e-13 | -5.195 | Dctd | 1.37e-03 | -4.505 |
| 1700126G02Rik | 8.60e-13 | -5.855 | Asxl3 | 1.51e-03 | -1.976 |
| Gm28307 | 3.33e-11 | -5.505 | Cdk8 | 1.62e-03 | -0.886 |
| Polr1b | 1.08e-10 | -3.977 | Gh | 1.77e-03 | 5.121 |
| Gm10062 | 1.40e-10 | -5.458 | Arglu1 | 2.37e-03 | -0.743 |

| Gene Name | adjusted p-value | log2FoldChange | Gene Name | adjusted p-value | log2FoldChange |
|---|---|---|---|---|---|
| AY036118 | 1.82e-10 | 1.152 | Gm47371 | 2.58e-03 | -4.505 |
| Sema3c | 2.13e-10 | -3.777 | Ptgds | 3.19e-03 | 3.081 |
| Wdr72 | 5.77e-10 | -5.434 | Gm15564 | 4.44e-03 | -3.324 |
| Gm36264 | 1.42e-09 | 5.807 | mt-Co2 | 6.77e-03 | 0.948 |
| Gm50306 | 4.71e-09 | -5.384 | 4930511M06Rik | 8.51e-03 | -3.214 |
| Pde1c | 9.89e-09 | -0.857 | Ms4a1 | 8.62e-03 | -3.799 |
| mt-Co3 | 2.39e-08 | 1.257 | Gm42726 | 8.82e-03 | -3.760 |
| Naaladl2 | 3.84e-08 | -1.427 | Gm17529 | 9.07e-03 | -4.358 |
| Gm12216 | 6.03e-08 | -2.311 | Rbm39 | 9.32e-03 | 0.716 |
| Csmd3 | 1.08e-07 | -2.229 | Rusc1 | 1.13e-02 | 1.070 |
| Chl1 | 1.55e-07 | -2.764 | Ttc39b | 1.37e-02 | -2.902 |
| Nckap5 | 2.17e-07 | -3.136 | 8430422H06Rik | 1.52e-02 | -1.860 |
| Zfp994 | 2.84e-07 | -5.279 | Gm14582 | 1.70e-02 | -4.075 |
| mt-Atp6 | 5.42e-07 | 1.154 | Zcchc18 | 1.95e-02 | 1.285 |
| Ano2 | 5.55e-07 | -5.043 | Epha6 | 1.99e-02 | -0.896 |
| Shb | 1.06e-06 | -3.569 | Prph | 2.11e-02 | 0.706 |
| Gm11337 | 1.08e-06 | -4.836 | Sil1 | 3.10e-02 | -1.353 |
| 6030443J06Rik | 1.37e-06 | -2.786 | Gm28578 | 3.16e-02 | -4.195 |
| Clk1 | 2.75e-06 | 1.533 | Map1a | 3.44e-02 | -1.088 |
| Ninj2 | 5.07e-06 | -4.195 | Rbm5 | 3.44e-02 | 0.950 |
| Pkd1 | 5.24e-06 | 1.079 | Tnrc6a | 3.70e-02 | 0.641 |
| mt-Co1 | 8.93e-06 | 1.502 | Aatk | 4.33e-02 | 1.109 |

p-cLTMR2

| Gene Name | adjusted p-value | log2FoldChange | Gene Name | adjusted p-value | log2FoldChange |
|---|---|---|---|---|---|
| Camk1d | 7.39e-93 | -2.290 | Wdr72 | 6.88e-08 | -5.238 |
| Hexb | 2.33e-43 | -3.415 | mt-Atp6 | 9.30e-08 | 0.940 |
| Xlr5a | 1.01e-41 | -6.147 | Adam11 | 1.09e-07 | 0.782 |
| Taco1 | 3.78e-40 | -3.642 | Ttc39b | 2.23e-07 | -4.290 |
| Snhg11 | 5.53e-40 | 0.870 | Gm42726 | 2.71e-07 | -4.835 |
| Bc1 | 1.21e-39 | 5.109 | Chl1 | 3.17e-07 | -2.050 |
| Sec61a1 | 1.27e-39 | -6.141 | Gabbr1 | 4.57e-07 | 1.266 |
| Lrp1b | 7.58e-38 | -2.733 | Il27 | 7.21e-07 | -3.238 |

| Gene Name | adjusted p-value | log2FoldChange | Gene Name | adjusted p-value | log2FoldChange |
|---|---|---|---|---|---|
| 4930567K12Rik | 1.70e-35 | 4.728 | Syne1 | 7.41e-07 | -0.736 |
| Gm19951 | 1.10e-29 | 2.373 | Arhgef25 | 1.03e-06 | 1.302 |
| 5830416I19Rik | 7.62e-27 | -6.787 | Epha6 | 1.09e-06 | -1.925 |
| Peak1 | 6.87e-23 | 2.190 | Rian | 1.14e-06 | 0.690 |
| Cmss1 | 1.98e-20 | -0.959 | 1700126G02Rik | 1.25e-06 | -4.404 |
| Sema3c | 2.50e-19 | -4.701 | Atp2b2 | 1.28e-06 | -0.794 |
| AY036118 | 8.43e-19 | 1.451 | Pcyt2 | 1.46e-06 | 1.387 |
| Klhl14 | 1.70e-18 | -6.010 | Pcdh15 | 1.63e-06 | -2.511 |
| Gm12216 | 2.07e-18 | -4.118 | Col5a3 | 2.05e-06 | 2.134 |
| 1700054A03Rik | 2.51e-17 | -6.348 | 6030443J06Rik | 2.24e-06 | -2.413 |
| Gm26917 | 2.68e-17 | -1.442 | Cntnap2 | 2.62e-06 | -0.705 |
| Ifngas1 | 5.60e-17 | 6.579 | Gh | 4.39e-06 | 4.704 |
| Rsrp1 | 7.85e-17 | 1.018 | Adamtsl1 | 4.45e-06 | -2.001 |
| AC149090.1 | 5.64e-16 | -1.012 | Gm11337 | 6.32e-06 | -4.835 |
| Gm28307 | 5.96e-16 | -5.999 | Ascc2 | 1.26e-05 | 1.279 |
| Ddx5 | 2.65e-15 | 1.140 | Rapgef4 | 1.36e-05 | -1.349 |
| Gm42418 | 1.20e-14 | -0.747 | Arhgap33 | 1.81e-05 | 1.322 |
| Lars2 | 1.58e-14 | -1.247 | Rbm39 | 1.96e-05 | 0.649 |
| Csmd3 | 5.05e-14 | -4.404 | Actb | 2.55e-05 | 1.974 |
| Dgkz | 1.92e-13 | 0.843 | Cpeb1 | 2.69e-05 | 0.959 |
| Shb | 5.31e-13 | -3.488 | Tnrc6a | 3.56e-05 | 0.632 |
| Prph | 5.74e-13 | 0.965 | Mylip | 4.56e-05 | -3.903 |
| Clk1 | 1.22e-12 | 1.333 | 4930557J02Rik | 5.78e-05 | -4.903 |
| Gm10062 | 5.29e-12 | -5.688 | Trim2 | 6.07e-05 | -1.181 |
| Mc1r | 6.98e-12 | -4.811 | Neat1 | 6.10e-05 | 1.289 |
| Gm50306 | 1.40e-11 | -5.989 | Zfp994 | 6.56e-05 | -3.989 |
| 8430422H06Rik | 2.77e-11 | -3.881 | Fyb2 | 8.96e-05 | -4.581 |
| Pde6a | 2.82e-11 | -4.892 | Edc4 | 9.22e-05 | 1.655 |
| Igf2bp3 | 2.91e-11 | -4.903 | Dazap1 | 1.08e-04 | 1.303 |
| Ntm | 6.78e-11 | -2.067 | Diablo | 1.28e-04 | 1.382 |
| Ptgds | 1.11e-10 | 3.512 | Cmpk1 | 2.10e-04 | -2.241 |
| Rnf112 | 1.22e-10 | 0.952 | Ivns1abp | 2.72e-04 | 1.119 |
| Gigyf1 | 4.16e-10 | 1.537 | Sf3b1 | 3.46e-04 | 1.182 |

| Gene Name | adjusted p-value | log2FoldChange | Gene Name | adjusted p-value | log2FoldChange |
|---|---|---|---|---|---|
| mt-Co3 | 5.77e-10 | 0.990 | Acoxl | 3.46e-04 | -3.618 |
| Fbxl7 | 7.80e-10 | -2.040 | Tmem108 | 3.62e-04 | -2.448 |
| Polr1b | 9.98e-10 | -3.084 | Nckap5 | 4.67e-04 | -2.103 |
| Pcbp4 | 3.95e-09 | 1.657 | Gm14582 | 5.13e-04 | -4.608 |
| Pde1c | 3.96e-09 | -0.691 | D430042O09Rik | 6.09e-04 | -1.830 |
| Arglu1 | 6.67e-09 | -0.684 | Lmbr1l | 6.38e-04 | 1.345 |
| Cdh23 | 1.22e-08 | -4.141 | Mtfr1l | 6.42e-04 | 1.224 |
| Prl | 2.63e-08 | 5.927 | Myl6 | 6.53e-04 | 1.719 |
| Naaladl2 | 4.81e-08 | -1.304 | Stx5a | 6.94e-04 | 1.413 |

Injured

| Gene Name | adjusted p-value | log2FoldChange | Gene Name | adjusted p-value | log2FoldChange |
|---|---|---|---|---|---|
| Col5a3 | 6.77e-69 | 3.560 | Smad1 | 1.08e-12 | 1.848 |
| Cacna2d1 | 9.39e-61 | 2.507 | Erc2 | 1.08e-12 | 1.072 |
| Gm29266 | 4.47e-51 | 3.915 | Ubash3b | 1.33e-12 | -2.371 |
| Atf3 | 9.42e-51 | 6.737 | A630023A22Rik | 1.89e-12 | 4.674 |
| Abca1 | 4.10e-47 | 2.976 | Gm29374 | 5.50e-12 | 6.690 |
| Mmp16 | 1.60e-45 | 3.429 | Uck2 | 5.75e-12 | 2.153 |
| Sema6a | 3.09e-45 | 4.214 | Plxna4 | 6.28e-12 | 1.603 |
| Itga7 | 1.85e-44 | 3.373 | Lsamp | 9.13e-12 | -1.964 |
| Nrip1 | 3.44e-43 | 3.575 | Anks1b | 1.29e-11 | 1.319 |
| Chl1 | 2.28e-39 | 3.748 | Shisa9 | 1.34e-11 | 4.467 |
| Sez6l | 8.34e-39 | 4.769 | Fam129b | 1.57e-11 | 2.017 |
| Lmo7 | 2.54e-37 | 4.233 | 4930544M13Rik | 5.21e-11 | 5.637 |
| Glis3 | 5.56e-37 | 4.645 | Igfn1 | 6.03e-11 | 5.052 |
| Zeb1 | 1.34e-36 | 1.977 | Prkg1 | 1.12e-10 | -1.232 |
| Tnik | 4.27e-34 | 2.041 | Bmp1 | 1.13e-10 | 2.115 |
| Ank3 | 9.47e-34 | 1.786 | Zbtb38 | 3.44e-10 | 2.168 |
| Gm33906 | 3.23e-33 | 5.659 | Il1r1 | 3.96e-10 | 3.101 |
| Myo10 | 2.64e-32 | 3.061 | Mfhas1 | 5.67e-10 | 1.715 |
| Nrg1 | 5.81e-30 | 2.001 | Camk2d | 7.41e-10 | 1.391 |
| Apba2 | 6.05e-30 | 2.300 | Sbno2 | 1.25e-09 | 2.077 |
| Sox11 | 4.37e-29 | 3.717 | Camk2b | 1.71e-09 | -4.577 |
| Sox5 | 1.96e-28 | 4.204 | Nav2 | 2.10e-09 | 1.248 |

| Gene Name | adjusted p-value | log2FoldChange | Gene Name | adjusted p-value | log2FoldChange |
|---|---|---|---|---|---|
| Rhoq | 1.02e-26 | 3.034 | Epb41l4a | 2.12e-09 | 2.556 |
| Spata13 | 2.14e-25 | 2.685 | Kirrel3 | 2.59e-09 | -3.855 |
| Tgfbi | 6.22e-25 | 6.467 | Gadd45a | 5.15e-09 | 3.520 |
| Stmn2 | 1.45e-24 | 1.959 | Bach2 | 5.17e-09 | -2.526 |
| Srrm4 | 8.23e-24 | 2.268 | Elavl3 | 6.81e-09 | 0.782 |
| D130009I18Rik | 5.40e-23 | -3.914 | Pou2f1 | 7.39e-09 | 2.308 |
| 1700025G04Rik | 7.84e-22 | 1.930 | Nkain1 | 7.97e-09 | 2.804 |
| Flna | 2.76e-20 | 1.841 | Stmn4 | 8.08e-09 | 2.660 |
| Rab31 | 1.27e-19 | 2.877 | Adam11 | 8.57e-09 | -2.246 |
| Sdk1 | 3.85e-19 | 2.155 | Ppp2r2b | 8.70e-09 | 1.226 |
| Csf1 | 1.22e-18 | 4.655 | Scn10a | 9.13e-09 | -4.120 |
| Ecel1 | 4.66e-18 | 6.052 | Vgf | 1.05e-08 | 3.588 |
| Gap43 | 4.80e-18 | 3.028 | Thsd7a | 1.06e-08 | 1.741 |
| Akap6 | 1.10e-17 | 1.444 | Zhx2 | 1.47e-08 | 2.603 |
| Bach1 | 1.17e-16 | 3.102 | Gpr158 | 1.59e-08 | 1.404 |
| Flrt3 | 8.94e-16 | 4.174 | Sprr1a | 1.67e-08 | 5.690 |
| Scpep1 | 2.60e-15 | 3.486 | Ppp1r1c | 2.48e-08 | -2.950 |
| Tet3 | 6.23e-15 | 1.998 | Runx2 | 2.64e-08 | 2.749 |
| Map7d1 | 9.12e-15 | 1.351 | Lpp | 4.27e-08 | 1.574 |
| Palld | 2.22e-14 | 2.776 | Cacna2d2 | 5.62e-08 | -3.297 |
| Rtn4 | 2.77e-14 | 1.775 | Syne1 | 6.50e-08 | -1.445 |
| Ltbp1 | 2.84e-14 | 3.439 | Dst | 7.68e-08 | 0.697 |
| Macrod2 | 5.03e-14 | -2.040 | Azin2 | 8.76e-08 | 1.768 |
| Adgrd1 | 6.02e-14 | 3.358 | Add2 | 8.81e-08 | 1.348 |
| Galnt9 | 2.60e-13 | 2.722 | Gal | 9.99e-08 | 4.970 |
| Dgkz | 3.18e-13 | -1.860 | Ppfia2 | 1.44e-07 | 1.311 |
| Syne2 | 3.31e-13 | 2.811 | Gm31831 | 1.71e-07 | 5.882 |
| Rnf122 | 1.04e-12 | 3.619 | Adarb1 | 1.80e-07 | 1.913 |

**Appendix Table 3:** IoN-CCI Day 21 top significant neuronal DEGs. Top 100 per subtype; adjusted p-value < 0.05; |log2FC| > 0.5.

NF

| Gene Name | adjusted p-value | log2FoldChange | Gene Name | adjusted p-value | log2FoldChange |
|---|---|---|---|---|---|
| Bc1 | 3.04e-101 | -1.760 | Dync1h1 | 3.29e-18 | -0.664 |
| Camk1d | 8.90e-98 | -0.998 | Ddx17 | 2.08e-17 | -0.625 |
| AC149090.1 | 6.83e-92 | -2.002 | D130009I18Rik | 2.55e-17 | 0.809 |
| Malat1 | 9.59e-80 | -0.531 | 9330159M07Rik | 3.24e-17 | 1.036 |
| Mycbp2 | 2.13e-76 | -1.114 | Pcdh7 | 4.16e-17 | 0.829 |
| Dgkb | 2.27e-70 | 1.670 | Huwe1 | 5.88e-17 | -0.956 |
| mt-Cytb | 4.76e-70 | -1.603 | Kmt2a | 1.04e-16 | -0.649 |
| mt-Atp6 | 7.29e-68 | -1.206 | Smpd4 | 1.41e-16 | -1.533 |
| mt-Nd4 | 1.94e-67 | -1.700 | Tmtc1 | 1.60e-16 | 0.933 |
| mt-Nd2 | 5.58e-57 | -1.682 | Zfand3 | 3.37e-16 | 0.957 |
| Snhg11 | 2.86e-56 | -0.638 | Nkain2 | 3.51e-16 | 0.606 |
| Srrm2 | 8.67e-48 | -0.840 | Prkn | 4.47e-16 | 0.638 |
| Lars2 | 4.74e-39 | -0.897 | Golga4 | 1.09e-15 | -0.988 |
| Son | 2.00e-38 | -1.063 | Atp2b1 | 2.35e-15 | -0.821 |
| Kif1a | 6.12e-34 | -0.884 | Mdga2 | 2.97e-15 | 0.763 |
| Gm47283 | 6.14e-33 | -1.001 | Nlgn1 | 4.26e-15 | 0.866 |
| Nrg3 | 8.03e-33 | 0.938 | Rbms3 | 4.82e-15 | 0.650 |
| Cpne6 | 2.11e-32 | -1.532 | Avil | 6.48e-15 | -0.729 |
| mt-Nd1 | 1.61e-31 | -1.236 | Luzp2 | 1.16e-14 | 0.799 |
| Tnrc6a | 2.66e-30 | -0.911 | Magi2 | 1.20e-14 | 0.607 |
| mt-Co3 | 3.57e-30 | -0.977 | Dync1i2 | 1.77e-14 | -0.745 |
| Prkg1 | 5.69e-30 | 0.893 | Slc35f1 | 3.60e-14 | 1.011 |
| Ttc3 | 7.13e-30 | -0.873 | Nktr | 5.35e-14 | -0.643 |
| Rbfox1 | 1.72e-29 | 0.858 | Sf3b1 | 6.07e-14 | -1.363 |
| Dgkz | 4.30e-29 | -0.951 | Ppm1e | 7.45e-14 | 0.898 |
| Schip1 | 5.03e-29 | 0.931 | Rbfox2 | 1.01e-13 | 0.691 |
| Gabrb3 | 2.54e-25 | 1.139 | Map4 | 1.17e-13 | 0.693 |
| Cdh18 | 3.16e-25 | 1.017 | Fam219a | 1.36e-13 | 0.747 |
| Peg3 | 9.22e-25 | -0.699 | Macrod2 | 1.88e-13 | 0.680 |
| Sgcz | 1.68e-24 | 0.843 | Oxr1 | 3.00e-13 | 0.799 |
| Rbm25 | 1.11e-23 | -0.725 | Agbl4 | 3.18e-13 | 0.707 |
| Akap8l | 1.63e-23 | -0.822 | Snrnp70 | 3.32e-13 | -0.560 |
| Cntnap2 | 1.66e-23 | 0.731 | Srpk2 | 4.51e-13 | 0.721 |
| Tom1l2 | 1.89e-23 | 0.837 | Arhgef25 | 4.82e-13 | -1.397 |

| Gene Name | adjusted p-value | log2FoldChange | Gene Name | adjusted p-value | log2FoldChange |
|---|---|---|---|---|---|
| Fam155a | 4.10e-23 | 0.770 | Rtn3 | 5.10e-13 | 0.673 |
| Dlg2 | 2.10e-22 | 0.998 | Ssh2 | 5.81e-13 | 0.993 |
| Map7d1 | 5.57e-22 | -0.961 | Ubr4 | 8.35e-13 | -0.802 |
| Lrfn5 | 3.75e-21 | 0.793 | Cntn1 | 9.46e-13 | 0.744 |
| Nrxn1 | 6.52e-21 | 0.708 | Nr6a1 | 9.61e-13 | 0.850 |
| Sptan1 | 3.29e-20 | -0.577 | Ccdc85a | 1.65e-12 | 0.692 |
| Ctnna2 | 7.45e-20 | 0.734 | Dtnb | 1.85e-12 | 0.902 |
| Agrn | 7.96e-20 | -1.600 | Cadps | 2.75e-12 | 0.541 |
| Prph | 2.98e-19 | -1.015 | B3galt1 | 3.12e-12 | 0.620 |
| Luc7l3 | 5.17e-19 | -0.618 | Prpf4b | 3.95e-12 | -1.002 |
| Robo1 | 9.26e-19 | 1.044 | Rsrp1 | 4.57e-12 | -0.561 |
| mt-Co2 | 1.29e-18 | -0.936 | Herc2 | 4.83e-12 | -0.947 |
| Gm42418 | 1.88e-18 | -0.646 | Ppp2r2b | 5.13e-12 | 0.692 |
| Pcdh9 | 2.12e-18 | 0.727 | Rcan2 | 5.97e-12 | 0.743 |
| Gpatch8 | 2.18e-18 | 0.679 | mt-Nd5 | 1.07e-11 | -1.496 |
| Ktn1 | 2.65e-18 | -0.883 | Fgf12 | 1.42e-11 | 0.696 |

NP

| Gene Name | adjusted p-value | log2FoldChange | Gene Name | adjusted p-value | log2FoldChange |
|---|---|---|---|---|---|
| AC149090.1 | 8.05e-36 | -1.708 | Tmcc1 | 3.96e-03 | 1.348 |
| Camk1d | 3.58e-27 | -1.113 | Ebf1 | 6.58e-03 | 0.594 |
| Dgkb | 3.26e-20 | 4.024 | Sgcz | 8.07e-03 | 1.093 |
| Bc1 | 3.14e-18 | -1.620 | Grm7 | 8.15e-03 | 0.582 |
| mt-Nd4 | 2.62e-15 | -1.375 | Adgrb3 | 8.37e-03 | 1.727 |
| Syt7 | 1.04e-14 | -0.794 | Gm32442 | 9.04e-03 | 1.101 |
| Srrm2 | 1.11e-12 | -0.792 | Samd12 | 9.09e-03 | 0.835 |
| Gm47283 | 1.89e-12 | -1.131 | Ncald | 1.04e-02 | 0.782 |
| mt-Cytb | 3.90e-12 | -1.275 | Ttc3 | 1.09e-02 | -0.859 |
| Galnt17 | 6.23e-12 | 1.046 | Prkn | 1.38e-02 | 0.561 |
| Snhg11 | 1.21e-11 | -0.597 | Zfp949 | 1.41e-02 | 1.319 |
| mt-Atp6 | 4.19e-10 | -1.012 | Hsp90ab1 | 1.45e-02 | -0.847 |
| Mycbp2 | 1.61e-09 | -0.949 | Sf3b1 | 1.53e-02 | -1.581 |
| Lars2 | 3.20e-09 | -0.933 | Ano3 | 1.89e-02 | 0.611 |
| Dgkz | 2.42e-08 | -0.849 | Tmtc1 | 1.98e-02 | 1.062 |

| Gene Name | adjusted p-value | log2FoldChange | Gene Name | adjusted p-value | log2FoldChange |
|---|---|---|---|---|---|
| Gm49678 | 3.11e-08 | 1.136 | mt-Nd3 | 2.02e-02 | -1.311 |
| mt-Nd2 | 4.34e-08 | -1.295 | Kif5a | 2.17e-02 | -0.656 |
| Ctnna2 | 4.89e-08 | 0.762 | Gna14 | 2.31e-02 | 0.730 |
| Sgcd | 5.59e-08 | 0.847 | Spink10 | 2.47e-02 | 2.178 |
| Nkain2 | 1.10e-07 | 0.918 | Fgf12 | 2.69e-02 | 0.653 |
| Csmd1 | 2.00e-07 | 0.820 | Ube2b | 2.87e-02 | 1.002 |
| Kcnt1 | 6.11e-07 | -0.920 | Rgs7 | 2.88e-02 | 0.554 |
| Gm42418 | 9.17e-07 | -0.673 | Prkg1 | 3.55e-02 | 0.646 |
| Ldb2 | 9.62e-07 | 0.773 | mt-Nd1 | 3.83e-02 | -0.699 |
| Nrg3 | 1.00e-06 | 0.830 | | | |
| Dync1i2 | 2.30e-06 | -0.888 | | | |
| Synpr | 6.00e-06 | 1.032 | | | |
| Lrfn5 | 1.12e-05 | 0.966 | | | |
| Tom1l2 | 2.13e-05 | 0.994 | | | |
| Kcnb2 | 3.09e-05 | 0.760 | | | |
| Map4 | 3.21e-05 | 0.906 | | | |
| Kcnip4 | 3.56e-05 | 0.753 | | | |
| Car10 | 5.06e-05 | 0.863 | | | |
| Gm26871 | 5.08e-05 | 0.540 | | | |
| mt-Co2 | 5.95e-05 | -0.915 | | | |
| Rbfox1 | 1.08e-04 | 0.697 | | | |
| Akap8l | 1.14e-04 | -1.119 | | | |
| mt-Co3 | 1.81e-04 | -0.912 | | | |
| Tnrc6a | 1.87e-04 | -0.980 | | | |
| Hdac9 | 1.92e-04 | 0.718 | | | |
| Robo2 | 4.48e-04 | 0.892 | | | |
| Syt9 | 5.66e-04 | 0.928 | | | |
| Grik1 | 6.58e-04 | 0.527 | | | |
| Kctd16 | 1.04e-03 | 0.674 | | | |
| Kif1a | 1.38e-03 | -0.560 | | | |
| Nrxn1 | 1.65e-03 | 0.583 | | | |
| Prph | 1.92e-03 | -0.793 | | | |
| Zfand3 | 2.11e-03 | 0.941 | | | |
| Polr1b | 2.16e-03 | 3.897 | | | |
| Gm19951 | 2.57e-03 | -0.564 | | | |

PEP1

| Gene Name | adjusted p-value | log2FoldChange | Gene Name | adjusted p-value | log2FoldChange |
|---|---|---|---|---|---|
| Camk1d | 2.27e-24 | -1.112 | | | |
| AC149090.1 | 3.92e-22 | -1.683 | | | |
| Dgkb | 8.93e-17 | 2.794 | | | |
| Snhg11 | 3.84e-15 | -0.713 | | | |
| Bc1 | 6.32e-12 | -1.353 | | | |
| Meg3 | 1.47e-10 | -0.508 | | | |
| Srrm2 | 6.35e-10 | -0.748 | | | |
| Gm47283 | 2.79e-09 | -1.178 | | | |
| Mycbp2 | 1.04e-08 | -0.846 | | | |
| Dync1i2 | 1.50e-07 | -1.148 | | | |
| Ptprf | 3.93e-06 | -0.942 | | | |
| Dgkz | 4.89e-06 | -1.066 | | | |
| Mir124a-1hg | 7.66e-06 | 1.679 | | | |
| Kctd16 | 2.39e-05 | 0.826 | | | |
| Peg3 | 4.07e-05 | -0.976 | | | |
| Bmper | 5.65e-05 | 0.728 | | | |
| Ap5z1 | 1.24e-04 | -3.079 | | | |
| mt-Nd4 | 2.99e-04 | -0.961 | | | |
| Syt7 | 3.81e-04 | -1.144 | | | |
| Lars2 | 5.34e-04 | -0.732 | | | |
| Herc2 | 6.96e-04 | -1.127 | | | |
| Prph | 7.92e-04 | -0.807 | | | |
| Zbtb20 | 1.04e-03 | 0.503 | | | |
| Gria1 | 2.10e-03 | 1.517 | | | |
| Cntnap2 | 2.20e-03 | 0.584 | | | |
| Map4 | 3.38e-03 | 0.698 | | | |
| Cadm2 | 4.05e-03 | 1.302 | | | |
| Pafah1b1 | 6.43e-03 | 0.754 | | | |
| Rian | 7.20e-03 | -0.566 | | | |
| mt-Atp6 | 7.33e-03 | -0.612 | | | |
| Zfand3 | 7.75e-03 | 0.949 | | | |
| Grm7 | 8.73e-03 | 0.535 | | | |
| Tom1l2 | 9.97e-03 | 0.711 | | | |

| Gene Name | adjusted p-value | log2FoldChange | Gene Name | adjusted p-value | log2FoldChange |
|---|---|---|---|---|---|
| Ddx17 | 1.06e-02 | -0.557 | | | |
| Ctnna2 | 1.14e-02 | 0.585 | | | |
| Plekha6 | 1.15e-02 | -0.533 | | | |
| Tnrc6a | 1.20e-02 | -0.694 | | | |
| mt-Cytb | 1.35e-02 | -0.717 | | | |
| Robo2 | 1.65e-02 | 1.262 | | | |
| Snrnp70 | 2.05e-02 | -0.717 | | | |
| Gm19951 | 2.21e-02 | -0.544 | | | |
| Dync1h1 | 3.31e-02 | -0.706 | | | |
| Fam222b | 3.71e-02 | 1.344 | | | |
| Prkcb | 4.32e-02 | 1.238 | | | |
| Herc1 | 4.41e-02 | -0.838 | | | |

PEP2

| Gene Name | adjusted p-value | log2FoldChange | Gene Name | adjusted p-value | log2FoldChange |
|---|---|---|---|---|---|
| AC149090.1 | 6.23e-36 | -2.020 | Ttc3 | 1.57e-05 | -0.719 |
| Camk1d | 5.51e-33 | -1.083 | Tom1l2 | 1.71e-05 | 0.851 |
| Mycbp2 | 1.24e-29 | -1.304 | Syt7 | 1.89e-05 | -0.662 |
| Bc1 | 2.16e-27 | -1.706 | Map7d1 | 3.36e-05 | -1.057 |
| Snhg11 | 1.27e-23 | -0.706 | Zbtb20 | 4.86e-05 | 0.659 |
| Meg3 | 3.66e-22 | -0.538 | Pabpc4 | 5.24e-05 | -2.290 |
| Dgkb | 1.20e-21 | 2.879 | Lsamp | 5.55e-05 | 1.053 |
| Prkg1 | 1.10e-19 | 0.992 | Xkr4 | 7.20e-05 | 1.028 |
| Srrm2 | 2.11e-18 | -0.960 | Dlg2 | 8.19e-05 | 0.840 |
| Grm7 | 3.60e-17 | 1.054 | Upp2 | 8.50e-05 | 0.952 |
| Malat1 | 3.08e-16 | -0.511 | Kcnmb2 | 1.14e-04 | 0.915 |
| Galnt17 | 2.82e-15 | 1.027 | Kcnh7 | 1.16e-04 | 0.748 |
| mt-Atp6 | 7.18e-15 | -1.000 | Akap8l | 1.44e-04 | -0.754 |
| Nrxn1 | 1.97e-14 | 0.938 | Gm26871 | 1.59e-04 | 0.758 |
| mt-Nd4 | 4.32e-14 | -1.292 | Golga4 | 2.44e-04 | -1.133 |
| Peg3 | 1.16e-13 | -1.056 | Luzp2 | 3.17e-04 | 0.765 |
| Nrg3 | 1.25e-13 | 0.746 | Ldb2 | 3.52e-04 | 0.811 |
| Cpne6 | 2.86e-13 | -1.614 | mt-Co3 | 5.40e-04 | -0.661 |
| mt-Cytb | 5.47e-13 | -1.170 | Peak1 | 5.45e-04 | -0.912 |

| Gene Name | adjusted p-value | log2FoldChange | Gene Name | adjusted p-value | log2FoldChange |
|---|---|---|---|---|---|
| Macrod2 | 1.26e-12 | 0.913 | Ccdc85a | 6.16e-04 | 0.947 |
| Schip1 | 2.48e-12 | 0.899 | Ktn1 | 7.32e-04 | -1.096 |
| Pcdh9 | 8.49e-12 | 0.806 | Ankrd24 | 8.87e-04 | -1.697 |
| Rbfox1 | 1.42e-10 | 0.714 | B3galt1 | 1.00e-03 | 0.761 |
| Gm42418 | 1.83e-10 | -0.616 | Tenm1 | 1.60e-03 | 0.845 |
| Kif1a | 4.55e-10 | -0.895 | Nr6a1 | 1.66e-03 | 1.228 |
| Tnrc6a | 5.63e-10 | -1.143 | Ftx | 1.90e-03 | 1.384 |
| Prph | 7.00e-10 | -0.942 | Map3k3 | 2.24e-03 | 1.279 |
| Dgkz | 3.51e-09 | -0.824 | Adam15 | 2.26e-03 | -1.721 |
| Gm47283 | 5.00e-09 | -1.033 | Parm1 | 2.30e-03 | 1.054 |
| Ralyl | 9.36e-09 | 0.872 | Otud7a | 2.33e-03 | 0.744 |
| Hjurp | 1.09e-08 | -0.862 | Gabrb3 | 2.36e-03 | 1.101 |
| Syne1 | 1.61e-08 | -0.776 | Rbm25 | 2.57e-03 | -0.640 |
| mt-Nd2 | 3.10e-08 | -1.221 | Hdac9 | 2.79e-03 | 0.880 |
| Son | 4.41e-08 | -0.949 | Col24a1 | 3.04e-03 | -1.506 |
| Gm49678 | 5.56e-08 | 0.959 | Kirrel3 | 3.20e-03 | 0.732 |
| Ctnna2 | 5.62e-08 | 0.872 | Cntnap2 | 3.72e-03 | 0.659 |
| Sptan1 | 2.28e-07 | -0.740 | Chd4 | 4.47e-03 | -0.840 |
| Lars2 | 3.42e-07 | -0.699 | Srsf11 | 4.75e-03 | -0.869 |
| Prkn | 5.54e-07 | 0.777 | Cacnb4 | 5.20e-03 | 0.582 |
| Slc4a3 | 1.01e-06 | -0.981 | Rsrp1 | 5.60e-03 | -0.709 |
| Kmt2a | 1.19e-06 | -0.819 | Mcu | 6.98e-03 | 1.242 |
| Luc7l3 | 2.49e-06 | -0.771 | Il1rapl1 | 8.67e-03 | 0.877 |
| Rbms3 | 4.74e-06 | 0.528 | 2210408F21Rik | 9.32e-03 | 0.993 |
| Fgf13 | 5.00e-06 | 1.498 | Snrnp70 | 1.04e-02 | -0.825 |
| Zcchc18 | 6.80e-06 | -1.607 | Smim14 | 1.07e-02 | 0.945 |
| Ndst4 | 7.32e-06 | 1.194 | Grid2 | 1.27e-02 | 0.915 |
| Mdga2 | 9.83e-06 | 0.917 | Prl | 1.30e-02 | -2.660 |
| Ddx17 | 1.37e-05 | -0.643 | Mir99ahg | 1.98e-02 | 0.912 |
| Herc1 | 1.37e-05 | -0.838 | Prpf4b | 2.06e-02 | -1.122 |
| Gm36264 | 1.39e-05 | -1.977 | Agrn | 2.19e-02 | -1.367 |

p-cLTMR2

| Gene Name | adjusted p-value | log2FoldChange | Gene Name | adjusted p-value | log2FoldChange |
|---|---|---|---|---|---|
| AC149090.1 | 6.80e-69 | -2.194 | Lars2 | 8.16e-09 | -0.842 |
| Camk1d | 2.09e-46 | -1.083 | Nktr | 8.79e-09 | -0.774 |
| Dgkb | 9.93e-45 | 3.132 | Rbms3 | 1.17e-08 | 0.540 |
| Mycbp2 | 1.87e-41 | -1.346 | Reep1 | 1.44e-08 | 1.051 |
| Meg3 | 2.31e-41 | -0.578 | Pde4d | 1.88e-08 | 0.552 |
| Snhg11 | 1.95e-37 | -0.666 | Hdac9 | 2.53e-08 | 0.628 |
| mt-Cytb | 2.63e-34 | -1.658 | Scn7a | 2.72e-08 | -0.818 |
| Malat1 | 5.91e-34 | -0.547 | Map7d1 | 3.07e-08 | -1.054 |
| Syt7 | 8.52e-31 | -0.980 | mt-Co3 | 3.91e-08 | -0.832 |
| Dgkz | 1.62e-30 | -1.037 | Arhgap33 | 5.37e-08 | -1.842 |
| mt-Atp6 | 2.42e-30 | -1.225 | Grik4 | 7.97e-08 | -0.625 |
| Kcnip4 | 7.96e-29 | 0.835 | Peg3 | 3.40e-07 | -1.132 |
| mt-Nd4 | 4.64e-26 | -1.623 | Rbm25 | 3.94e-07 | -0.583 |
| Schip1 | 4.93e-24 | 0.824 | Dpp10 | 4.18e-07 | 0.536 |
| Csmd1 | 6.14e-23 | 0.682 | 4732471J01Rik | 4.24e-07 | 1.089 |
| Ctnna2 | 3.23e-20 | 0.819 | Ttc9 | 4.29e-07 | 1.179 |
| Srrm2 | 5.48e-20 | -0.721 | Mbd6 | 1.21e-06 | -1.820 |
| D130009I18Rik | 4.34e-19 | 0.864 | Lrfn5 | 1.43e-06 | 0.574 |
| Gm47283 | 6.04e-18 | -1.102 | Grik1 | 1.62e-06 | 0.601 |
| mt-Nd2 | 2.71e-17 | -1.584 | Prkn | 1.98e-06 | 0.654 |
| Kctd16 | 5.43e-17 | 0.867 | Snrnp70 | 2.00e-06 | -0.741 |
| D130079A08Rik | 1.72e-16 | 0.899 | Rsrp1 | 2.23e-06 | -0.700 |
| Gm49678 | 5.67e-16 | 0.856 | Luc7l3 | 2.65e-06 | -0.621 |
| Fat3 | 1.23e-15 | 0.813 | Rcan2 | 2.69e-06 | 0.689 |
| Rbfox1 | 1.14e-14 | 0.648 | Maml2 | 4.88e-06 | 0.713 |
| Galnt17 | 3.27e-13 | 0.623 | Nol4 | 4.89e-06 | 0.835 |
| Ptprf | 3.29e-13 | -0.900 | Rian | 6.66e-06 | -0.552 |
| Kirrel3 | 3.56e-13 | 0.702 | Sptan1 | 7.03e-06 | -0.600 |
| Atp2b1 | 8.91e-13 | -0.736 | Ddx17 | 7.03e-06 | -0.507 |
| Frmd4a | 8.95e-13 | 0.659 | Ccser1 | 7.32e-06 | 0.631 |
| Akap8l | 1.19e-12 | -0.990 | Pex5l | 1.03e-05 | 0.620 |
| Tnrc6a | 1.34e-12 | -0.938 | Frmpd4 | 1.29e-05 | 0.570 |
| Fgf12 | 1.51e-12 | 0.747 | Sf3b1 | 1.34e-05 | -1.423 |
| Lsamp | 1.69e-12 | 0.593 | Map4 | 1.72e-05 | 0.704 |
| Kif1a | 3.87e-12 | -0.718 | Gm32442 | 1.77e-05 | 0.868 |

| Gene Name | adjusted p-value | log2FoldChange | Gene Name | adjusted p-value | log2FoldChange |
|---|---|---|---|---|---|
| Egfem1 | 4.24e-12 | 0.717 | Nhs | 1.89e-05 | 0.888 |
| Prkg1 | 7.65e-11 | 0.605 | Arhgef25 | 1.96e-05 | -1.336 |
| 4930567K12Rik | 8.28e-11 | 2.215 | Map3k3 | 2.30e-05 | 1.242 |
| Chd5 | 8.66e-11 | -0.725 | Nkain2 | 2.96e-05 | 0.588 |
| Rnf112 | 1.74e-10 | -1.166 | Ldb2 | 3.51e-05 | 0.651 |
| Nrxn1 | 2.04e-10 | 0.626 | Hcn1 | 4.01e-05 | 0.948 |
| Luzp2 | 2.56e-10 | 0.586 | Nrg3 | 5.35e-05 | 0.657 |
| Lhfpl3 | 4.19e-10 | 0.734 | Prph | 5.97e-05 | -0.650 |
| Ttc3 | 5.54e-10 | -0.960 | Dync1i2 | 6.04e-05 | -0.931 |
| Bc1 | 1.41e-09 | -1.470 | AY036118 | 8.62e-05 | 0.646 |
| mt-Nd1 | 2.20e-09 | -1.040 | Lingo2 | 8.94e-05 | 0.687 |
| Son | 2.60e-09 | -0.973 | Rgs8 | 9.27e-05 | 0.905 |
| Gm26871 | 2.76e-09 | 0.635 | Syt1 | 1.01e-04 | 0.528 |
| Macrod2 | 3.89e-09 | 0.651 | March11 | 1.45e-04 | 0.626 |
| Garnl3 | 5.04e-09 | 0.576 | Myo1a | 1.63e-04 | -1.214 |

**Appendix Table 4:** IoN Axotomy Day 7 top significant neuronal DEGs. Top 100 per subtype; adjusted p-value < 0.05; |log2FC| > 0.5.

NF

| Gene Name | adjusted p-value | log2FoldChange | Gene Name | adjusted p-value | log2FoldChange |
|---|---|---|---|---|---|
| Caln1 | 4.70e-90 | 4.746 | Htr1f | 2.33e-10 | -1.077 |
| Unc5cl | 7.92e-77 | 3.717 | Arid5b | 3.88e-10 | -1.461 |
| Glg1 | 4.16e-43 | 2.036 | Nat8l | 5.69e-10 | 0.642 |
| Spag5 | 3.39e-39 | 1.469 | Grik2 | 1.06e-09 | -1.082 |
| Rab23 | 2.09e-37 | 4.618 | Endod1 | 1.26e-09 | 0.930 |
| Catsperd | 7.54e-36 | 3.535 | Ncdn | 1.30e-09 | 1.243 |
| Macrod2 | 1.71e-35 | -1.124 | Myo10 | 1.49e-09 | 1.948 |
| Vps8 | 3.61e-35 | 3.324 | Plek2 | 1.68e-09 | 2.367 |
| Nkain2 | 3.82e-33 | 1.421 | Gria4 | 1.78e-09 | -1.027 |
| Nrg3 | 1.87e-31 | -0.930 | Thy1 | 5.24e-09 | 0.907 |
| Cinp | 9.40e-30 | 3.475 | Cplx1 | 5.45e-09 | 0.715 |
| Vps13a | 9.50e-27 | 1.881 | Stmn2 | 6.09e-09 | 0.666 |
| Dlg2.1 | 2.81e-26 | -0.956 | Syt11 | 8.03e-09 | 0.732 |
| Prkg1 | 2.67e-25 | -0.961 | Teddm2 | 8.42e-09 | 3.071 |
| Mad1l1 | 8.03e-25 | 1.278 | Rbfox1 | 8.48e-09 | -0.528 |
| Ebf1 | 3.65e-24 | -1.029 | Arfgef3 | 1.21e-08 | -1.730 |

| Gene Name | adjusted p-value | log2FoldChange | Gene Name | adjusted p-value | log2FoldChange |
|---|---|---|---|---|---|
| Pcdh7 | 6.43e-21 | -0.854 | Nefm | 1.31e-08 | 0.619 |
| Cadps | 2.88e-20 | -0.914 | Gpc5 | 1.95e-08 | -0.952 |
| Ddr2 | 1.16e-19 | 3.275 | Cdh18 | 2.95e-08 | -0.596 |
| Tatdn1 | 5.40e-19 | 3.620 | RP23-106G8.4 | 3.18e-08 | 4.899 |
| Pabpc1 | 7.45e-19 | 2.528 | Cntnap2 | 3.83e-08 | -0.547 |
| Prss41 | 7.98e-19 | 4.665 | Slc2a12 | 3.84e-08 | 2.371 |
| Brca2 | 1.78e-18 | 3.381 | Aldh2 | 4.13e-08 | 2.743 |
| Ptprd | 3.36e-18 | -0.676 | Rims1 | 6.15e-08 | -0.910 |
| Dlgap1 | 1.79e-17 | 1.158 | Heatr5a | 6.17e-08 | 1.898 |
| Ctnna2 | 1.97e-17 | -0.742 | Arhgap6 | 7.96e-08 | -1.567 |
| Trim33 | 2.42e-16 | 2.392 | Slc8a1 | 9.36e-08 | -0.912 |
| Zfp385b | 5.95e-16 | -0.881 | S100b | 2.24e-07 | 0.647 |
| Shcbp1l | 2.95e-15 | 4.104 | Magi2 | 2.27e-07 | -0.884 |
| Nrxn1 | 7.22e-15 | -0.688 | Lrfn5 | 3.04e-07 | -0.527 |
| Pcdh9 | 1.20e-14 | -0.676 | Pcp4 | 4.99e-07 | 0.764 |
| Calm3 | 2.07e-14 | 0.920 | Elavl2 | 5.64e-07 | -0.668 |
| Gabarapl1 | 4.03e-14 | 1.079 | Enox1 | 6.35e-07 | -0.771 |
| Hhla1 | 5.54e-14 | 4.122 | Atp1a1 | 1.11e-06 | 0.634 |
| Pla2g10 | 1.02e-13 | 4.106 | Vamp1 | 2.53e-06 | 0.804 |
| Spock2 | 1.32e-13 | 0.690 | Rplp1 | 2.84e-06 | 0.903 |
| Cpm | 3.44e-13 | 3.522 | Mtmr6 | 3.04e-06 | 0.667 |
| Luzp2 | 3.76e-13 | -0.757 | Plekhg3 | 3.18e-06 | 2.085 |
| Smurf2 | 5.06e-13 | 1.892 | Zwint | 3.31e-06 | 0.935 |
| Mapt | 6.36e-13 | 0.997 | Uchl1 | 3.79e-06 | 0.518 |
| Sgcd | 9.37e-13 | -0.929 | Lrrc17 | 4.26e-06 | 3.807 |
| Adgrl3 | 1.09e-12 | -0.962 | Rasal2 | 6.05e-06 | -0.807 |
| Psap | 3.62e-12 | 0.787 | Tmem245 | 6.25e-06 | 1.795 |
| Agbl4 | 1.61e-11 | -0.688 | Rnf187 | 6.86e-06 | 0.762 |
| Nbea | 3.79e-11 | -1.241 | Klc1 | 7.28e-06 | 0.519 |
| Unc5c | 4.40e-11 | -1.333 | Nrcam | 7.88e-06 | -1.330 |
| Dnajc17 | 6.76e-11 | 1.671 | Ralgapa2 | 8.08e-06 | -0.806 |
| B3galt1 | 1.02e-10 | -0.854 | Ppp2r2b | 9.25e-06 | -0.891 |
| Pex5l | 1.12e-10 | -0.751 | Rbfox2 | 9.67e-06 | 0.539 |
| Slc17a7 | 1.70e-10 | 1.472 | Cadm2 | 1.10e-05 | -0.666 |

PEP1

| Gene Name | adjusted p-value | log2FoldChange | Gene Name | adjusted p-value | log2FoldChange |
|---|---|---|---|---|---|
| Nkain2 | 5.39e-92 | 1.932 | Parp8 | 1.41e-08 | -1.519 |
| Caln1 | 8.93e-64 | 3.466 | Samd14 | 1.87e-08 | 0.895 |
| Catsperd | 2.48e-56 | 3.665 | Flot2 | 2.19e-08 | 0.887 |
| Vps13a | 2.02e-51 | 1.869 | Zfp850 | 2.24e-08 | 4.098 |
| Glg1 | 2.92e-49 | 1.756 | Tsc1 | 2.50e-08 | 1.480 |
| Pabpc1 | 1.06e-45 | 2.783 | Robo2 | 2.75e-08 | -0.805 |
| Unc5cl | 2.08e-45 | 3.212 | Basp1 | 2.87e-08 | 0.665 |
| Spag5 | 6.70e-45 | 1.111 | Dlgap1 | 3.17e-08 | 0.537 |
| Mad1l1 | 1.67e-34 | 1.180 | Ncdn | 5.11e-08 | 0.658 |
| Tatdn1 | 1.40e-33 | 4.085 | Fnbp1 | 5.63e-08 | 0.580 |
| Rab23 | 5.80e-29 | 3.648 | Aox2 | 6.64e-08 | 4.857 |
| Vps8 | 4.56e-23 | 2.162 | Galnt14 | 7.36e-08 | -0.857 |
| Nrxn1 | 5.66e-22 | -0.617 | Dnajc17 | 9.37e-08 | 1.234 |
| Mapt | 3.01e-21 | 0.584 | Mtch1 | 1.13e-07 | 0.920 |
| Gabarapl1 | 7.25e-21 | 1.305 | Rps29 | 1.17e-07 | 1.013 |
| RP23-106G8.4 | 1.05e-20 | 4.495 | Timp2 | 1.28e-07 | 0.610 |
| Cinp | 2.64e-20 | 3.052 | Celf2 | 1.42e-07 | -0.690 |
| Mrps27 | 3.26e-18 | 3.424 | Kcnc2 | 1.69e-07 | -0.832 |
| Trim33 | 4.35e-18 | 2.088 | Zwint | 2.28e-07 | 0.720 |
| Reep5 | 9.91e-18 | 0.914 | Lrp1b | 2.49e-07 | -1.007 |
| Syt11 | 1.02e-15 | 0.740 | Sag | 2.62e-07 | 4.541 |
| Mapk3 | 3.24e-14 | 0.576 | Mpz | 3.28e-07 | 1.383 |
| Myo10 | 4.99e-14 | 2.172 | Bcam | 3.32e-07 | 0.758 |
| Ctnna2 | 8.58e-14 | -0.670 | Vat1l | 3.45e-07 | 0.514 |
| Brca2 | 1.69e-13 | 3.027 | Nck1 | 4.54e-07 | 2.231 |
| Ptprd | 2.96e-13 | -0.921 | Smpdl3b | 8.97e-07 | 3.816 |
| Prss41 | 1.17e-12 | 3.613 | Slc4a4 | 9.34e-07 | -0.576 |
| Gnao1 | 1.56e-12 | 0.674 | Prkg1 | 9.41e-07 | -0.767 |
| Rab6b | 6.17e-12 | 0.577 | Strn3 | 9.86e-07 | -0.585 |
| Mcmbp | 1.02e-11 | 2.345 | Birc6 | 1.03e-06 | -0.995 |
| Lrrc48 | 1.34e-11 | 2.889 | R3hdm2 | 1.22e-06 | 1.101 |
| Shcbp1l | 1.47e-11 | 4.036 | Aes | 1.50e-06 | 1.046 |
| Lrrc17 | 1.55e-11 | 3.934 | Nrxn3 | 1.58e-06 | -0.637 |
| Smurf2 | 3.00e-11 | 1.582 | Sncg | 1.68e-06 | -1.092 |
| Plek2 | 4.12e-11 | 2.721 | Hhla1 | 1.69e-06 | 3.638 |
| Trnp1 | 5.27e-11 | 1.042 | Brip1os | 1.73e-06 | 1.334 |

| Gene Name | adjusted p-value | log2FoldChange | Gene Name | adjusted p-value | log2FoldChange |
|---|---|---|---|---|---|
| Lpp | 1.19e-10 | 0.734 | D17Wsu92e | 1.74e-06 | 0.682 |
| Nisch | 1.38e-10 | 0.612 | Kif13b | 2.86e-06 | -0.818 |
| Spock2 | 1.95e-10 | 0.640 | Luc7l2 | 3.09e-06 | -0.917 |
| Ddr2 | 7.04e-10 | 2.401 | Sufu | 3.18e-06 | 1.414 |
| Tulp4 | 9.08e-10 | -1.186 | Slc6a15 | 3.51e-06 | -0.738 |
| Ebf1 | 1.50e-09 | -0.822 | Cntnap2 | 4.71e-06 | -0.552 |
| Calm3 | 1.85e-09 | 0.670 | Kmt2c | 5.23e-06 | -0.996 |
| Pcgf3 | 2.37e-09 | 1.820 | Wfdc8 | 5.80e-06 | 4.575 |
| Kcnb2 | 4.41e-09 | -0.642 | Tcf20 | 7.42e-06 | -0.960 |
| Cpm | 7.44e-09 | 2.966 | Fus | 7.54e-06 | -0.809 |
| Scn3a | 7.52e-09 | 1.863 | Sacs | 7.81e-06 | -0.851 |
| Dpysl3 | 1.06e-08 | 0.565 | Arhgdia | 7.98e-06 | 0.507 |
| Arfgap3 | 1.31e-08 | 0.941 | Apoe | 1.08e-05 | -1.026 |
| Diablo | 1.40e-08 | 0.678 | Bri3bp | 1.34e-05 | 2.247 |

PEP2

| Gene Name | adjusted p-value | log2FoldChange | Gene Name | adjusted p-value | log2FoldChange |
|---|---|---|---|---|---|
| Caln1 | 2.49e-68 | 3.801 | Pkm | 1.09e-09 | 0.694 |
| Unc5cl | 3.77e-58 | 3.499 | Ldb2 | 1.31e-09 | -0.988 |
| Catsperd | 5.13e-57 | 3.407 | Uchl1 | 1.43e-09 | 0.674 |
| Pcdh9 | 4.65e-52 | -0.872 | Klc1 | 1.66e-09 | 0.658 |
| Nrg3 | 6.25e-50 | -0.904 | Lrrc17 | 1.89e-09 | 3.606 |
| Prkg1 | 5.47e-42 | -0.955 | Trnp1 | 2.08e-09 | 1.124 |
| B3galt1 | 1.02e-39 | -1.057 | Cntn3 | 3.29e-09 | -0.742 |
| Macrod2 | 1.25e-36 | -1.011 | Dynll2 | 3.70e-09 | 0.741 |
| Vps13a | 2.69e-34 | 1.918 | Pla2g10 | 5.08e-09 | 3.531 |
| Glg1 | 2.64e-33 | 1.583 | Hhla1 | 5.14e-09 | 3.745 |
| Vps8 | 1.85e-32 | 2.737 | Sepw1 | 6.52e-09 | 0.920 |
| Pabpc1 | 1.11e-31 | 2.689 | Scn4b | 9.37e-09 | 0.708 |
| Grm7 | 2.81e-28 | -0.833 | Rph3a | 9.50e-09 | 0.656 |
| Tatdn1 | 2.53e-27 | 3.895 | Zbtb20 | 9.74e-09 | -0.541 |
| Rbfox1 | 6.29e-27 | -0.882 | Cadm2 | 2.36e-08 | -0.840 |
| Spag5 | 1.33e-25 | 1.181 | Rims2 | 2.42e-08 | -1.053 |
| Cinp | 1.11e-22 | 3.665 | Syt11 | 2.62e-08 | 0.811 |
| Kcnh7 | 1.29e-22 | -0.818 | Mdga2 | 3.02e-08 | -0.788 |
| Rab23 | 1.25e-21 | 3.350 | Ap1g1 | 3.33e-08 | 1.435 |
| Gabarapl1 | 2.53e-20 | 1.545 | Thy1 | 4.11e-08 | 1.012 |

| Gene Name | adjusted p-value | log2FoldChange | Gene Name | adjusted p-value | log2FoldChange |
|---|---|---|---|---|---|
| Cntnap2 | 2.71e-20 | -0.760 | Rgs4 | 4.62e-08 | 0.552 |
| Mad1l1 | 2.71e-20 | 1.053 | Dnm3 | 4.74e-08 | -0.757 |
| Nrxn1 | 2.85e-20 | -0.731 | Reep5 | 7.47e-08 | 0.900 |
| Ctnna2 | 5.27e-20 | -0.804 | Bcam | 9.60e-08 | 0.731 |
| Ralyl | 1.94e-19 | -0.712 | Mapk3 | 1.03e-07 | 0.732 |
| Lrrc48 | 1.54e-18 | 3.711 | Robo2 | 1.29e-07 | -0.804 |
| Spock2 | 1.08e-16 | 0.782 | Atp1b1 | 1.36e-07 | 0.526 |
| Shcbp1l | 3.36e-16 | 6.457 | Kcnc2 | 1.51e-07 | -0.947 |
| Cplx1 | 4.93e-16 | 0.935 | Lpp | 1.82e-07 | 0.921 |
| RP23-106G8.4 | 6.86e-16 | 3.673 | Kcnab2 | 2.43e-07 | 0.595 |
| Mrps27 | 7.23e-16 | 3.229 | Rnf187 | 2.47e-07 | 1.046 |
| Ddr2 | 1.10e-15 | 2.927 | Hs6st3 | 2.65e-07 | -0.957 |
| Trim33 | 2.70e-14 | 1.944 | Tpbgl | 3.17e-07 | 1.603 |
| Brca2 | 2.94e-14 | 2.821 | Aox2 | 3.30e-07 | 4.459 |
| Lsamp | 4.61e-14 | -0.908 | Tuba1a | 3.62e-07 | 0.693 |
| Smurf2 | 5.55e-14 | 2.001 | Pcp4l1 | 3.64e-07 | 1.286 |
| Cend1 | 1.12e-13 | 1.421 | Fndc5 | 4.33e-07 | 0.940 |
| Calm3 | 1.57e-13 | 0.961 | Mcmbp | 4.73e-07 | 2.206 |
| Prss41 | 3.65e-12 | 5.176 | Cabp1 | 4.94e-07 | 0.859 |
| Hpca | 4.64e-12 | 1.088 | Tenm1 | 5.79e-07 | -0.796 |
| Ncdn | 1.22e-11 | 1.114 | Dlgap1 | 5.89e-07 | 0.857 |
| Rora | 2.67e-11 | -0.689 | Dpysl3 | 1.12e-06 | 0.818 |
| Pam | 2.78e-11 | -1.294 | Erc2 | 1.22e-06 | -0.792 |
| Psap | 2.88e-11 | 0.677 | Eno2 | 1.31e-06 | 0.688 |
| Tppp3 | 8.17e-11 | 0.916 | Ywhab | 2.53e-06 | 1.049 |
| Syt1 | 1.37e-10 | -0.600 | Cox8a | 2.54e-06 | 0.511 |
| Kif5a | 3.19e-10 | 0.702 | Ctsb | 2.89e-06 | 0.760 |
| Ebf1 | 3.88e-10 | -0.894 | Ppp2r2b | 2.99e-06 | -1.106 |
| Rab6b | 6.25e-10 | 0.801 | Xkr4 | 3.02e-06 | -0.998 |
| Arfgap3 | 1.05e-09 | 0.884 | Stmn2 | 3.14e-06 | 0.807 |

NP

| Gene Name | adjusted p-value | log2FoldChange | Gene Name | adjusted p-value | log2FoldChange |
|---|---|---|---|---|---|
| Trim33 | 1.14e-06 | 2.935 | | | |
| Caln1 | 1.87e-06 | 3.144 | | | |
| Unc5cl | 5.13e-03 | 2.452 | | | |
| Catsperd | 2.56e-02 | 3.012 | | | |

SST

| Gene Name | adjusted p-value | log2FoldChange | Gene Name | adjusted p-value | log2FoldChange |
|---|---|---|---|---|---|
| Catsperd | 5.12e-06 | 4.693 | | | |
| Caln1 | 1.18e-05 | 4.639 | | | |
| Mapt | 7.81e-04 | 1.403 | | | |
| Pabpc1 | 1.43e-03 | 3.362 | | | |
| Unc5cl | 2.05e-03 | 4.140 | | | |
| Nkain2 | 1.57e-02 | 1.412 | | | |
| Tatdn1 | 3.96e-02 | 3.815 | | | |

p-cLTMR2

| Gene Name | adjusted p-value | log2FoldChange | Gene Name | adjusted p-value | log2FoldChange |
|---|---|---|---|---|---|
| Caln1 | 1.98e-47 | 3.749 | | | |
| Nkain2 | 4.74e-39 | 1.825 | | | |
| Unc5cl | 2.07e-31 | 3.439 | | | |
| Catsperd | 7.20e-21 | 3.679 | | | |
| Glg1 | 1.02e-20 | 1.722 | | | |
| Pabpc1 | 5.67e-19 | 2.725 | | | |
| Spag5 | 2.22e-18 | 1.112 | | | |
| Lsamp | 9.74e-18 | -0.872 | | | |
| Vps13a | 1.78e-17 | 1.744 | | | |
| Dpp10 | 2.75e-14 | -0.680 | | | |
| Mrps27 | 1.37e-12 | 3.967 | | | |
| Mad1l1 | 4.90e-12 | 1.066 | | | |
| Ebf1 | 1.43e-11 | -0.806 | | | |
| Vps8 | 4.36e-11 | 2.313 | | | |
| Lrrc48 | 4.71e-11 | 3.379 | | | |
| Rab23 | 3.63e-10 | 3.351 | | | |
| Ctnna2 | 3.73e-10 | -0.741 | | | |
| Tatdn1 | 5.84e-10 | 3.003 | | | |
| Cinp | 1.03e-08 | 3.124 | | | |
| Prkg1 | 4.08e-08 | -0.735 | | | |
| Pde4d | 3.16e-07 | -0.643 | | | |
| Smurf2 | 1.13e-06 | 1.864 | | | |
| RP23-106G8.4 | 2.97e-06 | 3.580 | | | |
| Lpp | 3.10e-06 | 1.112 | | | |
| Mcmbp | 3.84e-06 | 2.411 | | | |

| Gene Name | adjusted p-value | log2FoldChange | Gene Name | adjusted p-value | log2FoldChange |
|---|---|---|---|---|---|
| Syt11 | 1.69e-05 | 0.776 | | | |
| Luzp2 | 1.74e-05 | -0.569 | | | |
| Cpm | 3.82e-05 | 3.696 | | | |
| Mapt | 4.53e-05 | 0.600 | | | |
| Syt1 | 5.51e-05 | -0.576 | | | |
| Reep5 | 9.15e-05 | 0.883 | | | |
| Scn3a | 1.64e-04 | 1.548 | | | |
| Brca2 | 4.18e-04 | 3.160 | | | |
| Gnao1 | 5.68e-04 | 0.869 | | | |
| Trim33 | 1.70e-03 | 1.319 | | | |
| Aox2 | 2.61e-03 | 6.324 | | | |
| Cep112 | 4.32e-03 | 2.706 | | | |
| Pacsin1 | 6.14e-03 | 0.607 | | | |
| Hhla1 | 7.29e-03 | 4.075 | | | |
| Mapk8ip1 | 8.46e-03 | 1.399 | | | |
| Dnm3 | 1.22e-02 | -0.790 | | | |
| Uchl1 | 1.63e-02 | 0.575 | | | |
| Pigk | 1.81e-02 | -1.050 | | | |
| Nrg4 | 1.85e-02 | 3.214 | | | |
| Grik1 | 2.47e-02 | -0.652 | | | |
| Pam | 2.76e-02 | -1.100 | | | |
| Ano3 | 3.06e-02 | -0.555 | | | |
| Gata4 | 3.47e-02 | 5.717 | | | |
| Snx7 | 4.44e-02 | -1.104 | | | |

Injured

| Gene Name | adjusted p-value | log2FoldChange | Gene Name | adjusted p-value | log2FoldChange |
|---|---|---|---|---|---|
| Cacna2d1 | 2.83e-191 | 2.402 | Prdm12 | 2.14e-47 | -2.776 |
| Sema6a | 1.74e-177 | 3.974 | Palld | 2.24e-47 | 2.025 |
| Tnik | 1.98e-173 | 3.050 | Sdk1 | 1.66e-45 | 1.802 |
| Zfp13 | 3.59e-164 | 3.853 | Brca2 | 2.58e-45 | 1.776 |
| Atf3 | 1.49e-148 | 5.090 | Ddr2 | 6.86e-45 | 1.784 |
| Mmp16 | 7.02e-143 | 3.078 | Plek2 | 1.54e-44 | 1.953 |
| Gfra1 | 5.51e-127 | 3.703 | Pou2f1 | 2.32e-44 | 2.429 |
| Mad1l1 | 1.34e-113 | 1.629 | Abca1 | 8.41e-44 | 2.198 |
| Spag5 | 1.05e-110 | 1.629 | Pcdh15 | 1.16e-43 | 2.413 |
| Unc5cl | 3.48e-109 | 1.842 | Srxn1 | 4.79e-43 | 1.998 |
| Rbms3 | 4.26e-103 | -1.257 | Kcnab2 | 1.07e-42 | -1.595 |

| Gene Name | adjusted p-value | log2FoldChange | Gene Name | adjusted p-value | log2FoldChange |
|---|---|---|---|---|---|
| Caln1 | 8.71e-103 | 1.721 | Kcnb2 | 1.25e-42 | -1.673 |
| Etl4 | 2.70e-91 | 2.749 | Rab31 | 1.89e-42 | 2.466 |
| Sprr1a | 1.19e-89 | 6.348 | Nts | 2.31e-42 | 3.895 |
| Dlgap1 | 5.06e-89 | 1.668 | Macrod2 | 1.80e-41 | -1.251 |
| Sox11 | 1.47e-87 | 2.745 | Auts2 | 2.65e-41 | -1.711 |
| Mapt | 3.58e-87 | 1.509 | Ldb2 | 4.51e-41 | -2.860 |
| Glis3 | 2.46e-85 | 5.556 | Adgrl3 | 7.87e-41 | -1.444 |
| Tshz2 | 6.39e-79 | -2.134 | Cinp | 8.01e-41 | 1.594 |
| Itga7 | 4.59e-76 | 2.631 | Elavl4 | 9.11e-41 | -1.094 |
| Myo10 | 4.82e-75 | 1.958 | Cadm1 | 3.43e-40 | -1.499 |
| Gpr158 | 7.79e-73 | 2.070 | Kcnq3 | 4.12e-40 | 2.184 |
| Zbtb20 | 1.41e-71 | -2.056 | Pirt | 4.59e-40 | -2.466 |
| Nrip1 | 2.72e-70 | 2.489 | Calca | 5.72e-40 | -3.048 |
| Chl1 | 1.48e-69 | 2.441 | Shcbp1l | 7.78e-40 | 2.299 |
| Itgb6 | 1.30e-68 | 4.777 | Klhl1 | 1.88e-39 | 2.284 |
| Prkg1 | 1.85e-68 | -1.061 | Flrt3 | 2.29e-39 | 2.994 |
| RP24-348E18.1 | 2.13e-67 | 4.735 | Dgkh | 3.84e-39 | -2.006 |
| Map7d1 | 2.20e-67 | 1.591 | Calm1 | 6.58e-39 | -1.205 |
| Ube2l3 | 1.22e-65 | 1.972 | Spata13 | 1.78e-38 | 1.722 |
| Akap6 | 1.66e-62 | 1.365 | Terb1 | 2.48e-38 | 2.732 |
| Pcnx | 1.66e-62 | 2.472 | Chst2 | 3.08e-38 | 2.837 |
| Ank3 | 4.18e-62 | 1.331 | Rhd | 4.19e-38 | 1.684 |
| Fstl1 | 1.37e-61 | -1.495 | Fam169a | 7.66e-38 | 2.988 |
| Pip5k1b | 2.35e-61 | 2.713 | Fst | 1.17e-37 | 5.579 |
| Atp1a1 | 1.05e-60 | -1.513 | Rbms1 | 1.44e-37 | -1.197 |
| Grid2 | 2.20e-60 | 1.773 | Anks1b | 1.70e-37 | 1.539 |
| Tac1 | 9.48e-59 | -3.708 | Adam8 | 4.68e-37 | 3.673 |
| Rasgrf2 | 1.24e-58 | 2.804 | Gap43 | 1.92e-36 | 1.501 |
| Rhoq | 2.47e-56 | 2.442 | Fam19a2 | 5.27e-36 | 2.132 |
| Rab23 | 1.12e-55 | 1.865 | Dgkz | 5.68e-36 | -1.639 |
| Nav2 | 2.11e-54 | 1.876 | Kcna2 | 1.81e-35 | -2.515 |
| H2-T23 | 2.90e-52 | 3.220 | Enox1 | 5.48e-35 | -1.311 |
| RP23-416J8.1 | 8.63e-52 | 2.215 | Runx1 | 9.12e-35 | -2.488 |
| Man1a | 8.63e-51 | 3.410 | Frem3 | 1.25e-34 | 4.963 |
| Brip1os | 1.85e-50 | 2.003 | Ubash3b | 2.96e-34 | -2.049 |
| Bptf | 2.22e-50 | 1.824 | Spock2 | 3.23e-34 | -1.114 |
| Col5a3 | 2.27e-49 | 2.440 | Dmd | 5.05e-34 | 1.625 |

| Gene Name | adjusted p-value | log2FoldChange | Gene Name | adjusted p-value | log2FoldChange |
|---|---|---|---|---|---|
| Phf24 | 7.34e-49 | -2.135 | Arhgap42 | 6.21e-34 | 2.775 |
| Stmn2 | 1.06e-47 | 1.167 | Rgs4 | 7.79e-34 | -1.968 |

**Appendix Table 5:** IoN Axotomy Day 21 top significant neuronal DEGs. Top 100 per subtype; adjusted p-value < 0.05; |log2FC| > 0.5.

NF

| Gene Name | adjusted p-value | log2FoldChange | Gene Name | adjusted p-value | log2FoldChange |
|---|---|---|---|---|---|
| Rpl41 | 8.37e-23 | -1.683 | Plec | 1.32e-04 | -0.769 |
| Mt3 | 2.55e-15 | -2.186 | Pnn | 1.36e-04 | 1.156 |
| Rpl38-ps2 | 2.87e-15 | -3.084 | Arhgap35 | 1.91e-04 | -0.825 |
| Rbm4b | 2.40e-14 | 2.243 | Them6 | 2.27e-04 | -1.569 |
| Kif1a | 3.52e-13 | -0.938 | Rps25 | 2.50e-04 | -2.345 |
| Rsrp1 | 4.69e-13 | 0.809 | Rps15 | 2.83e-04 | -1.520 |
| Fth1.1 | 5.35e-13 | 1.212 | Tubb4a | 3.06e-04 | -1.651 |
| Spock2 | 1.38e-10 | -0.865 | Sepw1 | 3.43e-04 | -0.962 |
| Snhg11 | 6.33e-10 | 0.539 | Kcna1 | 4.51e-04 | -0.989 |
| Gphn | 1.94e-09 | 0.625 | Map1a | 4.76e-04 | -0.586 |
| Maf | 2.02e-09 | -0.708 | Cox6c | 7.62e-04 | -0.989 |
| Mad1l1 | 3.00e-09 | -0.860 | Rapgef4os1 | 7.65e-04 | 1.020 |
| Nat8l | 3.25e-09 | -0.814 | Nfia | 8.53e-04 | -0.597 |
| Arf3 | 6.48e-09 | -1.261 | Vmn2r30 | 8.72e-04 | 3.105 |
| Ndufa5 | 7.07e-09 | -1.834 | Fam160b2 | 1.00e-03 | -1.649 |
| Cox7c | 1.66e-08 | -2.183 | Spock1 | 1.02e-03 | -0.868 |
| Actb | 1.76e-08 | -1.176 | Scn8a | 1.16e-03 | -0.652 |
| Pygb | 1.80e-08 | -0.859 | Stxbp5l | 1.30e-03 | 0.566 |
| Scn3a | 2.27e-08 | 2.341 | S100a6 | 1.30e-03 | -1.701 |
| Tmsb10 | 2.50e-08 | -1.748 | Ndufa2 | 1.45e-03 | -1.323 |
| Slc24a5 | 3.54e-08 | 1.696 | Arhgef25 | 1.85e-03 | -1.182 |
| Tacc1 | 4.43e-08 | -1.330 | Ndufb7 | 2.26e-03 | -1.415 |
| Mettl23 | 7.37e-08 | 1.103 | Rpl37rt | 3.12e-03 | -1.501 |
| Zfhx3 | 9.84e-08 | -0.835 | Clstn1 | 3.24e-03 | -0.764 |
| Rtn1 | 1.86e-07 | -0.631 | Rps28 | 3.33e-03 | -2.935 |
| Srrm2 | 5.20e-07 | -1.118 | Grin1 | 3.57e-03 | -1.030 |
| Ebf1 | 6.29e-07 | -0.692 | Tceb2 | 3.71e-03 | -1.571 |
| Grina | 8.23e-07 | -1.295 | Prune2 | 4.40e-03 | 0.612 |
| Mdga2 | 9.02e-07 | 0.513 | Kcnc3 | 4.45e-03 | -1.115 |
| Igf2bp3 | 1.15e-06 | 1.697 | Spag5 | 5.16e-03 | -0.633 |

| Gene Name | adjusted p-value | log2FoldChange | Gene Name | adjusted p-value | log2FoldChange |
|---|---|---|---|---|---|
| Srsf2 | 1.45e-06 | 1.117 | Tomm7 | 5.32e-03 | -1.633 |
| Zranb2 | 3.64e-06 | 1.051 | Bsn | 6.38e-03 | -1.127 |
| Eno2 | 5.50e-06 | -0.707 | Myeov2 | 6.38e-03 | -0.955 |
| Rps29 | 7.46e-06 | -2.377 | Agpat1 | 8.43e-03 | -2.331 |
| Uqcr11 | 8.50e-06 | -1.686 | Caln1 | 8.57e-03 | 1.229 |
| Psap | 9.27e-06 | -0.594 | Trpc1 | 8.92e-03 | 0.889 |
| Cox6a1 | 1.37e-05 | -1.007 | Gigyf1 | 9.15e-03 | -2.340 |
| Romo1 | 1.48e-05 | -2.287 | Rpl23 | 1.03e-02 | -1.600 |
| Kcnq1 | 2.04e-05 | 1.770 | Ptprn | 1.06e-02 | -0.830 |
| Atp1a1 | 2.44e-05 | -0.703 | Cox5b | 1.13e-02 | -1.468 |
| Vps13a | 2.93e-05 | -1.119 | Tuba1b | 1.19e-02 | -0.505 |
| Sult4a1 | 3.40e-05 | -0.759 | Cox6b1 | 1.26e-02 | -0.893 |
| Rpl28-ps1 | 3.76e-05 | -2.615 | Abhd2 | 1.34e-02 | -1.243 |
| Tom1l2 | 3.98e-05 | -0.771 | Sept1 | 1.41e-02 | 2.858 |
| Aplp2 | 5.00e-05 | -0.754 | Rplp1 | 1.57e-02 | -0.806 |
| Dpp6 | 5.16e-05 | -0.719 | Suv420h2 | 1.67e-02 | -2.769 |
| L1cam | 6.54e-05 | -1.561 | Iqsec1 | 1.90e-02 | -0.910 |
| Pet100 | 1.11e-04 | -1.568 | Tnrc6a | 2.00e-02 | -1.092 |
| Clasp2 | 1.22e-04 | -0.908 | Atp5k | 2.09e-02 | -2.090 |
| Tmem126a | 1.26e-04 | 1.275 | Atp8a2 | 2.73e-02 | -0.671 |

PEP1

| Gene Name | adjusted p-value | log2FoldChange | Gene Name | adjusted p-value | log2FoldChange |
|---|---|---|---|---|---|
| Tmsb10 | 3.09e-43 | -2.073 | Cbfb | 1.25e-05 | 0.555 |
| Rpl41 | 9.89e-41 | -1.708 | Rpl21 | 1.35e-05 | -1.562 |
| Fth1.1 | 1.38e-38 | 1.118 | Pcdh1 | 1.55e-05 | -1.505 |
| Mt3 | 5.42e-26 | -2.338 | Tmsb4x | 1.69e-05 | 0.539 |
| Rpl38-ps2 | 1.07e-22 | -2.939 | Mrps27 | 3.21e-05 | 2.233 |
| Rps29 | 1.35e-22 | -3.314 | Cox6c | 3.23e-05 | -1.224 |
| L1cam | 9.00e-22 | -1.743 | St13 | 6.46e-05 | 1.082 |
| Calm2 | 4.50e-21 | 0.934 | Krit1 | 6.62e-05 | 0.757 |
| RP24-163J22.10 | 4.38e-20 | 3.681 | Add2 | 6.71e-05 | -0.802 |
| S100a6 | 3.91e-18 | -1.224 | Aqp1 | 7.05e-05 | -0.967 |
| Gnas | 2.06e-17 | 0.960 | Pygb | 7.09e-05 | -1.064 |
| Prune2 | 7.22e-17 | 0.743 | Mcmbp | 7.35e-05 | 1.729 |
| Mettl23 | 1.06e-14 | 1.390 | Atp1a1 | 7.67e-05 | -0.525 |

| Gene Name | adjusted p-value | log2FoldChange | Gene Name | adjusted p-value | log2FoldChange |
|---|---|---|---|---|---|
| Slc24a5 | 8.47e-14 | 1.860 | Gabarapl2 | 1.01e-04 | 0.913 |
| Kif1a | 1.69e-13 | -0.778 | Rps15 | 1.15e-04 | -1.378 |
| Rbm4b | 5.13e-13 | 2.060 | Strbp | 1.20e-04 | 0.811 |
| Slc25a4 | 2.89e-12 | 1.080 | Ndufa5 | 1.39e-04 | -1.508 |
| Arf3 | 2.84e-11 | -1.100 | Vstm2l | 1.57e-04 | -1.374 |
| Sepw1 | 8.16e-11 | -1.071 | Mapt | 1.87e-04 | -0.537 |
| Prph | 9.06e-11 | 0.595 | Pfn2 | 1.95e-04 | 0.680 |
| Uqcr11 | 3.81e-10 | -1.972 | Ndufa2 | 2.64e-04 | -1.367 |
| Romo1 | 5.54e-10 | -2.096 | Erdr1 | 3.09e-04 | 1.513 |
| Actb | 3.06e-09 | -0.746 | Slc43a2 | 3.98e-04 | -1.561 |
| Srsf2 | 4.55e-09 | 1.093 | Calca | 4.13e-04 | 0.560 |
| Ctsl | 4.60e-09 | 1.247 | Nrg3os | 4.67e-04 | 1.035 |
| Spock2 | 5.05e-09 | -0.929 | Dlg4 | 5.38e-04 | -1.453 |
| Grip1 | 8.84e-09 | -0.642 | Ftl1 | 5.73e-04 | 0.545 |
| Tac1 | 1.00e-08 | 0.794 | Cep170b | 6.27e-04 | -1.174 |
| Atp5k | 2.25e-08 | -3.126 | Cox8a | 6.86e-04 | -0.723 |
| Cd9 | 2.27e-08 | 0.697 | Sdc4 | 7.38e-04 | -3.859 |
| Rpl37rt | 3.49e-08 | -1.530 | Rps15-ps2 | 7.95e-04 | -1.716 |
| Cox6a1 | 3.69e-08 | -0.945 | Rpl28-ps1 | 1.54e-03 | -1.991 |
| Nefl | 3.96e-08 | 0.705 | Igf2bp3 | 1.61e-03 | 1.576 |
| Rsrp1 | 4.46e-08 | 0.749 | Nrn1 | 1.71e-03 | 0.888 |
| Nlrc4 | 5.25e-08 | -1.543 | Syp | 1.76e-03 | -1.470 |
| Bri3bp | 5.43e-08 | 2.194 | Ndufa3 | 1.87e-03 | -1.897 |
| Fnbp1 | 8.42e-08 | -1.095 | Rnf7 | 1.91e-03 | 1.211 |
| St3gal4 | 1.04e-07 | -1.225 | RP23-112D14.3 | 2.88e-03 | 2.558 |
| Scn9a | 5.84e-07 | 0.571 | Prkar1b | 3.65e-03 | -0.762 |
| Psap | 1.49e-06 | -0.678 | Fxyd4 | 4.68e-03 | 2.564 |
| Srsf5 | 1.55e-06 | 1.005 | Sept7 | 6.38e-03 | 0.845 |
| Gphn | 1.63e-06 | 0.665 | Scn3a | 7.94e-03 | 1.304 |
| Cox7c | 1.94e-06 | -1.942 | Fam160b2 | 9.09e-03 | -1.288 |
| Atp9a | 2.14e-06 | -0.812 | Glg1 | 9.67e-03 | -0.738 |
| Arl8a | 2.46e-06 | -0.901 | Wdr89 | 1.07e-02 | -0.841 |
| Igf1r | 2.60e-06 | -0.863 | Rpl3-ps1 | 1.13e-02 | 1.222 |
| Anxa5 | 6.37e-06 | 0.768 | RP23-106G8.4 | 1.14e-02 | 2.730 |
| Lrrc48 | 7.62e-06 | 2.007 | B3galt1 | 1.17e-02 | 0.759 |
| Stmn2 | 9.29e-06 | 0.526 | Auts2 | 1.32e-02 | -0.526 |
| Gnb1 | 1.11e-05 | 0.582 | Rps19 | 1.41e-02 | -1.192 |

PEP2

| Gene Name | adjusted p-value | log2FoldChange | Gene Name | adjusted p-value | log2FoldChange |
|---|---|---|---|---|---|
| Fth1.1 | 7.83e-50 | 1.382 | Mif | 9.23e-07 | -1.839 |
| Mt3 | 2.04e-36 | -2.521 | Gnas | 1.72e-06 | 0.802 |
| Rpl41 | 9.44e-27 | -1.594 | Nlrc4 | 1.72e-06 | -1.633 |
| Tmsb10 | 2.14e-25 | -2.256 | Tceb2 | 3.12e-06 | -1.582 |
| Pla2g3 | 6.87e-25 | 2.076 | Dgkz | 4.99e-06 | -0.966 |
| Rpl38-ps2 | 1.13e-20 | -2.915 | Psap | 5.45e-06 | -0.624 |
| Rbm4b | 1.11e-16 | 2.119 | Vamp2 | 6.00e-06 | -1.137 |
| Prune2 | 5.07e-15 | 0.593 | Clasp2 | 6.60e-06 | -0.740 |
| Nefl | 1.34e-14 | 0.804 | Ube2m | 8.92e-06 | -2.545 |
| Rora | 1.42e-14 | -0.585 | Vwa1 | 1.12e-05 | 1.804 |
| Lrrc48 | 2.83e-14 | 2.701 | Mapk3 | 1.39e-05 | -0.810 |
| S100a6 | 4.02e-14 | -1.163 | Bri3bp | 2.04e-05 | 1.392 |
| Grip1 | 5.00e-13 | -0.642 | Abhd2 | 2.67e-05 | -1.055 |
| Mrps27 | 6.17e-13 | 2.416 | Fam160b2 | 3.09e-05 | -1.642 |
| Cox7c | 1.25e-12 | -1.997 | Igf2bp3 | 3.44e-05 | 1.569 |
| Mettl23 | 1.43e-12 | 1.195 | Rps27rt | 3.54e-05 | -0.986 |
| Trpm8 | 3.69e-12 | -0.675 | Tomm7 | 5.78e-05 | -1.941 |
| Rps29 | 6.30e-12 | -2.850 | St3gal5 | 6.64e-05 | -3.171 |
| Kif1a | 3.46e-11 | -0.694 | Slc25a4 | 7.72e-05 | 0.871 |
| Rps15 | 4.03e-11 | -1.732 | Grina | 9.46e-05 | -0.988 |
| Atp5j2 | 4.81e-11 | -1.776 | Romo1 | 1.19e-04 | -1.676 |
| Uqcr11 | 6.99e-11 | -1.663 | Rps13-ps2 | 1.25e-04 | -2.066 |
| Spock2 | 7.57e-11 | -0.948 | Kcnq1 | 1.46e-04 | 2.083 |
| Atp1a1 | 1.45e-10 | -0.714 | Nap1l5 | 1.50e-04 | 1.005 |
| Srsf2 | 5.88e-10 | 1.141 | Atp9a | 1.83e-04 | -0.875 |
| Rph3a | 8.05e-10 | -0.831 | Ppp6r1 | 2.07e-04 | -2.305 |
| Prph | 1.95e-09 | 0.522 | Ptprs | 2.14e-04 | -0.702 |
| Cox6b1 | 2.07e-09 | -1.217 | Ifi27 | 3.17e-04 | 1.128 |
| Ndufa5 | 3.63e-09 | -1.721 | Tubb4a | 3.83e-04 | -1.452 |
| RP23-112D14.3 | 7.01e-09 | 1.937 | Shcbp1l | 4.08e-04 | 4.611 |
| Scn3a | 7.39e-09 | 1.325 | Rhd | 4.90e-04 | -1.548 |
| L1cam | 1.01e-08 | -1.303 | Spag5 | 5.06e-04 | -0.534 |
| Spag16 | 1.73e-08 | 2.140 | Ndufa4 | 5.30e-04 | -0.861 |
| Mad1l1 | 1.74e-08 | -0.703 | Rps6ka2 | 5.68e-04 | -0.870 |
| Calm2 | 1.99e-08 | 0.789 | Arf3 | 6.57e-04 | -0.845 |
| Tmsb4x | 2.33e-08 | 0.756 | Gabarapl2 | 7.77e-04 | 0.974 |
| Ndufa3 | 2.92e-08 | -2.055 | Zranb2 | 7.94e-04 | 0.870 |

| Gene Name | adjusted p-value | log2FoldChange | Gene Name | adjusted p-value | log2FoldChange |
|---|---|---|---|---|---|
| Fnbp1 | 3.42e-08 | -1.027 | Tuba1a | 8.50e-04 | 0.621 |
| Spock1 | 4.67e-08 | -0.827 | Pygb | 9.57e-04 | -0.877 |
| Ebf1 | 5.18e-08 | -0.509 | Uqcrq | 1.15e-03 | -1.085 |
| Rsrp1 | 6.93e-08 | 0.720 | Rps23-ps1 | 1.26e-03 | -1.588 |
| Actb | 1.51e-07 | -0.900 | Pet100 | 1.38e-03 | -1.149 |
| Mcmbp | 1.87e-07 | 1.658 | Krit1 | 1.51e-03 | 0.560 |
| Cox6c | 2.08e-07 | -0.969 | Slc43a2 | 1.77e-03 | -1.203 |
| Slc24a5 | 2.91e-07 | 1.558 | Agpat1 | 1.99e-03 | -2.535 |
| RP23-357C16.1 | 3.71e-07 | -0.744 | Ccdc88a | 2.03e-03 | 0.579 |
| Erdr1 | 4.46e-07 | 1.757 | Vmn2r30 | 2.23e-03 | 3.063 |
| Tmem126a | 4.93e-07 | 1.390 | Prss41 | 2.28e-03 | 2.941 |
| Hpca | 7.53e-07 | -1.222 | Stmn2 | 2.37e-03 | 0.598 |
| Ftl1 | 9.13e-07 | 0.753 | Rps15-ps2 | 2.55e-03 | -2.087 |

NP

| Gene Name | adjusted p-value | log2FoldChange | Gene Name | adjusted p-value | log2FoldChange |
|---|---|---|---|---|---|
| Dgkz | 2.77e-44 | -1.006 | Rpl37rt | 1.40e-07 | -1.263 |
| Fth1.1 | 1.43e-41 | 1.175 | Pet100 | 1.62e-07 | -1.702 |
| Tmsb10 | 6.21e-38 | -2.082 | Ap3s1 | 1.82e-07 | 0.914 |
| Rpl38-ps2 | 1.26e-30 | -3.775 | Prmt8 | 2.16e-07 | -1.559 |
| Rpl41 | 2.32e-28 | -1.462 | Slc25a4 | 3.30e-07 | 0.865 |
| S100a6 | 5.67e-27 | -1.288 | Ctsl | 3.39e-07 | 1.084 |
| Kif1a | 2.41e-26 | -0.804 | Pygb | 4.18e-07 | -1.084 |
| Grip1 | 5.39e-25 | -0.789 | Lrrc48 | 6.20e-07 | 2.130 |
| Ebf1 | 5.94e-24 | -0.611 | Spock2 | 6.34e-07 | -0.896 |
| Mt3 | 4.00e-23 | -2.533 | Vamp2 | 7.60e-07 | -1.120 |
| Calm2 | 1.12e-22 | 0.909 | Aqp1 | 9.18e-07 | -1.172 |
| Prune2 | 1.21e-22 | 0.677 | St3gal4 | 9.80e-07 | -1.230 |
| L1cam | 1.20e-20 | -1.390 | Cox6a1 | 1.87e-06 | -0.856 |
| Rbm4b | 6.07e-20 | 2.113 | Slc43a2 | 2.75e-06 | -1.618 |
| Ssbp3 | 3.63e-19 | -1.166 | Cd55 | 3.08e-06 | 0.643 |
| Rps29 | 6.59e-19 | -2.837 | Srsf5 | 6.53e-06 | 0.842 |
| Scn3a | 1.49e-17 | 2.180 | Nrn1 | 6.89e-06 | 0.665 |
| Arpc1b | 2.46e-17 | 1.271 | Tceb2 | 7.21e-06 | -1.281 |
| Spag5 | 8.83e-17 | -0.723 | Pcmtd2 | 8.81e-06 | 0.996 |
| Nefl | 6.08e-16 | 0.825 | Ubash3b | 9.47e-06 | -0.772 |
| Uqcr11 | 8.07e-16 | -2.260 | Rps19 | 1.35e-05 | -1.917 |

| Gene Name | adjusted p-value | log2FoldChange | Gene Name | adjusted p-value | log2FoldChange |
|---|---|---|---|---|---|
| RP24-163J22.10 | 1.32e-15 | 3.878 | Capn1 | 1.42e-05 | -1.454 |
| Fnbp1 | 1.54e-14 | -1.050 | Syp | 1.44e-05 | -1.247 |
| Mad1l1 | 1.68e-14 | -0.724 | Cdkn1b | 1.69e-05 | -2.168 |
| Arf3 | 2.51e-14 | -1.157 | Pfn2 | 1.82e-05 | 0.841 |
| Mettl23 | 4.18e-14 | 1.188 | Slc36a1 | 1.87e-05 | -0.817 |
| Rps27rt | 1.40e-13 | -1.565 | Bsn | 1.94e-05 | -0.823 |
| Scoc | 7.84e-13 | 1.275 | Rps16-ps2 | 2.67e-05 | -1.434 |
| Ftl1 | 2.27e-12 | 0.797 | Igf2bp3 | 2.77e-05 | 1.868 |
| Fxyd2 | 6.41e-12 | -0.790 | Nap1l5 | 2.86e-05 | 1.109 |
| Nedd4l | 3.78e-11 | -0.536 | Caln1 | 3.00e-05 | 1.090 |
| Il6st | 1.54e-10 | -0.917 | Romo1 | 3.18e-05 | -1.371 |
| Cox7c | 2.31e-10 | -2.142 | Apc2 | 4.11e-05 | -1.359 |
| Ndufa5 | 2.50e-10 | -1.971 | Atp9a | 4.21e-05 | -0.774 |
| Actb | 2.84e-10 | -0.976 | RP23-106G8.4 | 5.37e-05 | 2.502 |
| Mapt | 7.30e-10 | -0.584 | Cox5b | 6.19e-05 | -1.240 |
| Add2 | 9.04e-10 | -0.801 | Rtn4 | 6.27e-05 | 0.550 |
| Rsrp1 | 1.07e-09 | 0.684 | Rps15 | 6.43e-05 | -1.339 |
| Aplp2 | 1.16e-09 | -0.865 | Cd44 | 7.39e-05 | -0.551 |
| Gnas | 1.42e-09 | 0.761 | Ndufs6 | 7.73e-05 | -2.548 |
| Zwint | 1.66e-09 | 0.825 | Rpl28-ps1 | 1.21e-04 | -1.619 |
| Tmem126a | 3.47e-09 | 1.365 | Plec | 1.46e-04 | -0.849 |
| Pifo | 5.02e-09 | 1.568 | Rps15-ps2 | 1.57e-04 | -2.008 |
| Nlrc4 | 6.46e-09 | -1.711 | Bag1 | 1.57e-04 | 1.347 |
| Slc24a5 | 6.46e-09 | 1.388 | Dtx3 | 1.66e-04 | -1.248 |
| Srsf2 | 7.41e-09 | 1.011 | Apba1 | 1.75e-04 | -0.513 |
| Bri3bp | 9.48e-09 | 2.155 | Madd | 1.87e-04 | -1.114 |
| Gphn | 1.02e-08 | 0.696 | Camk2a | 1.99e-04 | -0.840 |
| Cep170b | 2.44e-08 | -1.275 | Dhcr24 | 2.37e-04 | -0.860 |
| Mrps27 | 7.63e-08 | 2.189 | Sumo1 | 2.56e-04 | 1.153 |

SST

| Gene Name | adjusted p-value | log2FoldChange | Gene Name | adjusted p-value | log2FoldChange |
|---|---|---|---|---|---|
| S100a6 | 1.46e-04 | -1.079 | | | |
| Rpl41 | 9.15e-04 | -1.224 | | | |
| Arf3 | 2.59e-02 | -1.870 | | | |

p-cLTMR2

| Gene Name | adjusted p-value | log2FoldChange | Gene Name | adjusted p-value | log2FoldChange |
|---|---|---|---|---|---|
| Fth1.1 | 2.45e-20 | 1.172 | | | |
| Rpl41 | 1.61e-19 | -1.906 | | | |
| Prune2 | 3.67e-17 | 0.744 | | | |
| Grip1 | 1.38e-14 | -0.822 | | | |
| Tmem126a | 1.08e-09 | 1.476 | | | |
| Tmsb4x | 1.64e-08 | 0.812 | | | |
| Tmsb10 | 2.08e-08 | -1.786 | | | |
| Mapt | 3.16e-08 | -1.033 | | | |
| Rbm4b | 4.44e-08 | 2.363 | | | |
| Rpl38-ps2 | 1.92e-07 | -2.933 | | | |
| Nefl | 2.89e-07 | 0.849 | | | |
| Mettl23 | 5.21e-07 | 1.275 | | | |
| Gnas | 8.13e-07 | 1.185 | | | |
| Calm2 | 9.42e-07 | 0.808 | | | |
| Rps29 | 1.90e-06 | -2.799 | | | |
| Srsf2 | 4.38e-06 | 1.236 | | | |
| Arf3 | 1.51e-05 | -1.364 | | | |
| Fam160b2 | 5.71e-05 | -2.602 | | | |
| Pfkfb2 | 6.79e-05 | 3.733 | | | |
| Rpl37rt | 1.08e-04 | -1.781 | | | |
| Mt3 | 1.11e-04 | -2.520 | | | |
| Mrps27 | 2.32e-04 | 2.449 | | | |
| Apba1 | 2.55e-04 | -0.780 | | | |
| Zfhx3 | 3.13e-04 | -0.677 | | | |
| Vmn2r30 | 7.26e-04 | 4.407 | | | |
| Kif1a | 7.39e-04 | -0.620 | | | |
| Prph | 8.44e-04 | 0.517 | | | |
| Rps15 | 1.23e-03 | -1.919 | | | |
| Add2 | 1.41e-03 | -1.037 | | | |
| Ssbp3 | 1.64e-03 | -1.018 | | | |
| Fxyd2 | 2.58e-03 | -0.749 | | | |
| Pygb | 2.96e-03 | -1.256 | | | |
| Vamp2 | 3.48e-03 | -1.505 | | | |
| Spag5 | 4.29e-03 | -0.760 | | | |
| Glg1 | 5.58e-03 | -0.926 | | | |
| Spag16 | 7.68e-03 | 2.638 | | | |
| Atp1a1 | 9.91e-03 | -0.701 | | | |

| Gene Name | adjusted p-value | log2FoldChange | Gene Name | adjusted p-value | log2FoldChange |
|---|---|---|---|---|---|
| Ndufa4 | 1.35e-02 | -1.164 | | | |
| Cdh9 | 1.58e-02 | 0.793 | | | |
| Rpl34-ps1 | 2.31e-02 | -2.003 | | | |
| Slc25a4 | 2.64e-02 | 1.008 | | | |
| Tubb4a | 2.82e-02 | -1.750 | | | |
| Ftl1 | 3.57e-02 | 0.609 | | | |
| Nedd4l | 4.02e-02 | -0.603 | | | |
| Mad1l1 | 4.22e-02 | -0.784 | | | |
| S100a6 | 4.67e-02 | -1.125 | | | |

Injured

| Gene Name | adjusted p-value | log2FoldChange | Gene Name | adjusted p-value | log2FoldChange |
|---|---|---|---|---|---|
| Tnik | 8.05e-195 | 2.994 | Nlgn1 | 2.35e-47 | 1.791 |
| Mmp16 | 1.43e-183 | 2.994 | Gadd45a | 1.44e-46 | 2.838 |
| Cacna2d1 | 7.11e-165 | 2.123 | Syt7 | 4.26e-46 | -3.203 |
| Sema6a | 2.03e-141 | 4.038 | Cacna1c | 5.01e-46 | -2.144 |
| Sox11 | 1.74e-137 | 2.971 | Grm7 | 2.38e-45 | -1.424 |
| Atf3 | 6.94e-137 | 5.398 | Ppp1r1c | 4.03e-45 | -1.711 |
| Etl4 | 1.19e-120 | 2.754 | RP23-416J8.1 | 1.43e-44 | 2.186 |
| Pcdh15 | 8.57e-113 | 3.247 | Mad1l1 | 1.75e-44 | 1.227 |
| Gfra1 | 1.82e-110 | 3.780 | Dlg2.1 | 2.91e-44 | 0.881 |
| Man1a | 1.12e-103 | 5.129 | Srxn1 | 5.86e-44 | 2.175 |
| Zfp13 | 1.79e-102 | 3.466 | Flrt3 | 3.53e-43 | 2.665 |
| Gpr158 | 1.04e-100 | 2.289 | Pcdh7 | 4.83e-43 | 0.744 |
| Grid2 | 5.23e-90 | 1.910 | Enox1 | 8.54e-43 | -1.431 |
| Phactr2 | 1.70e-88 | 1.968 | Itga7 | 6.34e-41 | 2.244 |
| Chl1 | 1.48e-87 | 2.938 | Synpr | 4.01e-40 | -3.391 |
| Fam19a2 | 5.19e-86 | 3.026 | Rbms3 | 6.58e-40 | -0.908 |
| Rasgrf2 | 1.86e-84 | 3.205 | Prkg1 | 1.15e-39 | -0.935 |
| Glis3 | 1.87e-84 | 5.136 | Itgb6 | 1.52e-39 | 3.918 |
| Tshz2 | 5.46e-83 | -2.479 | Dlgap1 | 5.35e-39 | 1.174 |
| Sprr1a | 2.46e-79 | 5.995 | Dgki | 6.23e-39 | -2.613 |
| Map7d1 | 8.52e-77 | 1.883 | Auts2 | 8.12e-39 | -1.750 |
| Gap43 | 5.74e-76 | 2.285 | Stxbp5l | 1.21e-38 | 1.552 |
| Nrip1 | 1.26e-74 | 2.331 | Rab31 | 1.57e-38 | 2.769 |
| Nts | 4.45e-72 | 4.314 | Snrpn | 2.74e-38 | 2.570 |
| Stmn2 | 3.24e-71 | 1.452 | Myo10 | 6.40e-38 | 2.062 |

| Gene Name | adjusted p-value | log2FoldChange | Gene Name | adjusted p-value | log2FoldChange |
|---|---|---|---|---|---|
| Dmd | 2.76e-70 | 1.990 | Pde11a | 9.61e-38 | -4.206 |
| Zbtb20 | 4.14e-68 | -2.180 | Scn10a | 1.44e-37 | -3.743 |
| Ldb2 | 6.79e-67 | -3.697 | Ano4 | 3.65e-37 | 1.577 |
| Pappa | 8.24e-67 | 3.849 | Rgs4 | 5.37e-37 | -2.297 |
| Pcnx | 3.25e-66 | 2.516 | Gabrg1 | 1.30e-36 | 2.519 |
| Pou2f1 | 4.85e-65 | 2.692 | Prss12 | 1.39e-36 | 3.060 |
| Akap6 | 1.20e-64 | 1.224 | Abca1 | 4.14e-36 | 1.732 |
| Cadm1 | 2.21e-63 | -2.661 | Nedd4l | 6.28e-36 | -2.109 |
| Chst2 | 1.14e-62 | 3.313 | Grik1 | 9.57e-36 | -4.450 |
| Nav2 | 2.01e-62 | 2.090 | Pbx1 | 3.58e-35 | 1.770 |
| Map2 | 5.27e-61 | 1.792 | Dennd1a | 9.71e-35 | 1.595 |
| Fam155a | 3.14e-60 | 1.511 | Hs3st5 | 2.65e-34 | 1.272 |
| Spag5 | 3.77e-59 | 1.340 | Sorcs3 | 3.94e-33 | 2.217 |
| Palld | 5.45e-59 | 2.571 | Fhod3 | 6.54e-33 | 2.776 |
| Sdk1 | 3.58e-57 | 1.925 | Cd55 | 2.70e-32 | -3.454 |
| Kcnb2 | 4.11e-55 | -2.045 | Fam171b | 2.94e-32 | 1.578 |
| Npy | 8.30e-55 | 6.046 | Fndc3a | 3.04e-32 | 1.835 |
| Arhgap42 | 3.55e-53 | 2.828 | Sgcd | 6.14e-32 | -1.651 |
| Nkain3 | 8.20e-53 | 3.479 | Pirt | 6.24e-32 | -2.320 |
| Nwd2 | 9.50e-53 | -3.000 | Enah | 6.35e-32 | 1.174 |
| Rhoq | 2.98e-51 | 2.139 | Ltbp1 | 9.79e-32 | 4.022 |
| Kcnq3 | 2.95e-50 | 2.243 | Pappa2 | 2.22e-31 | 5.088 |
| Dgkh | 1.54e-49 | -2.377 | Camk1 | 3.78e-31 | 2.760 |
| Cttnbp2 | 3.73e-49 | 2.664 | Runx1 | 4.85e-31 | -2.480 |
| Ell2 | 6.82e-48 | 2.688 | Plcxd3 | 6.47e-31 | -2.209 |

**Appendix Table 6:** DRG ScNT Day 7 top significant neuronal DEGs. Top 100 per subtype; adjusted p-value < 0.05; |log2FC| > 0.5.

NF

| Gene Name | adjusted p-value | log2FoldChange | Gene Name | adjusted p-value | log2FoldChange |
|---|---|---|---|---|---|
| Hbb-bs | 2.05e-50 | 2.919 | Onecut2 | 3.70e-07 | -1.141 |
| Map1b | 1.66e-40 | -0.725 | Gap43 | 3.91e-07 | 0.899 |
| Nefh | 1.48e-37 | -1.063 | P3h3 | 3.99e-07 | 2.403 |
| Sprr1a | 1.56e-36 | 7.161 | Samhd1 | 4.66e-07 | 1.417 |
| Ttf1 | 2.86e-35 | 2.523 | Abhd12 | 5.32e-07 | -1.319 |
| Hbb-bt | 7.46e-34 | 3.465 | Rab33b | 6.18e-07 | 1.667 |
| Hspa8 | 2.34e-24 | 0.696 | Rps2 | 1.32e-06 | 1.314 |
| Phkg1 | 2.73e-23 | 1.766 | Nptx2 | 2.34e-06 | 1.821 |

| Gene Name | adjusted p-value | log2FoldChange | Gene Name | adjusted p-value | log2FoldChange |
|---|---|---|---|---|---|
| Ccdc62 | 2.65e-22 | 2.054 | Etl4 | 2.58e-06 | -1.662 |
| Ngfr | 1.13e-21 | 0.980 | Nefm | 3.38e-06 | -0.604 |
| Gal | 1.74e-21 | 4.345 | Kcna1 | 3.47e-06 | -0.826 |
| Bcam | 1.73e-18 | 1.168 | Serinc1 | 4.23e-06 | -0.590 |
| S100a6 | 1.01e-17 | 1.139 | Apoe | 4.57e-06 | 0.944 |
| Hba-a2 | 1.60e-17 | 2.629 | Necab1 | 4.77e-06 | -1.385 |
| Tubb5 | 7.75e-17 | 0.653 | Ap1ar | 6.19e-06 | -1.200 |
| Pmp22 | 3.07e-16 | 0.933 | Scg5 | 8.52e-06 | -1.538 |
| Als2 | 3.85e-16 | 1.714 | Apod | 9.47e-06 | 1.672 |
| Myl6 | 4.36e-15 | 0.781 | Rps18 | 1.01e-05 | 1.215 |
| Stmn3 | 8.02e-14 | 0.599 | Ntrk1 | 1.28e-05 | 1.039 |
| Thy1 | 8.56e-14 | 0.546 | Slc24a2 | 1.44e-05 | -1.012 |
| Noct | 3.18e-13 | -1.188 | Exosc2 | 1.77e-05 | 1.890 |
| Vamp2 | 3.49e-13 | 0.906 | Nedd4 | 1.87e-05 | -0.707 |
| Vgf | 2.13e-12 | 1.201 | Calca | 2.02e-05 | 0.751 |
| Nefl | 3.70e-12 | -0.604 | Vamp1 | 2.60e-05 | -0.957 |
| Map7d2 | 3.82e-12 | -0.899 | Rpl38 | 2.64e-05 | 0.612 |
| Unc13d | 7.87e-12 | 1.498 | Fhl1 | 2.87e-05 | -1.194 |
| Selenow | 3.71e-11 | 1.388 | Cdkn1a | 2.87e-05 | 1.820 |
| Snap25 | 5.51e-11 | -0.638 | Eif5a2 | 3.17e-05 | -0.911 |
| Eef1a2 | 6.54e-11 | 0.624 | Rps29 | 3.94e-05 | 0.820 |
| Npy | 8.40e-11 | 5.615 | Osbpl9 | 4.07e-05 | -1.614 |
| Ahnak2 | 8.99e-11 | -0.708 | Nup98 | 4.64e-05 | 1.901 |
| BC051077 | 1.06e-10 | 1.714 | Atf5 | 4.72e-05 | 1.637 |
| Elmo1 | 2.22e-10 | -1.277 | Sept5 | 5.24e-05 | 0.749 |
| Hpgds | 2.69e-10 | 1.982 | Ntrk3 | 5.33e-05 | -1.257 |
| Tubb2b | 3.77e-10 | 1.000 | Taldo1 | 5.33e-05 | 0.954 |
| Dynll2 | 2.01e-09 | 0.510 | Atp9a | 6.24e-05 | 0.687 |
| Cd9 | 5.20e-09 | 0.849 | Etv5 | 6.71e-05 | 1.119 |
| Psap | 6.22e-09 | 0.519 | Sparcl1 | 7.46e-05 | -1.323 |
| Kcnab2 | 1.01e-08 | 0.596 | Mapk3 | 9.07e-05 | 0.622 |
| Mtmr6 | 1.18e-08 | -0.826 | Golga3 | 1.06e-04 | 1.023 |
| Trim43b | 1.72e-08 | 2.322 | Unc80 | 1.39e-04 | -1.064 |
| Sptan1 | 2.09e-08 | -0.905 | Dpysl2 | 1.42e-04 | -0.552 |
| Glg1 | 2.97e-08 | -0.813 | Itch | 1.50e-04 | 1.117 |
| Wdr89 | 3.66e-08 | 2.521 | Scn3a | 1.71e-04 | -6.197 |
| BC024386 | 8.69e-08 | 2.370 | C1qa | 2.06e-04 | 4.983 |
| Sag | 9.13e-08 | 1.680 | Hn1 | 2.16e-04 | 1.084 |
| Tecr | 9.14e-08 | 0.622 | Basp1 | 2.18e-04 | 0.817 |

| Gene Name | adjusted p-value | log2FoldChange | Gene Name | adjusted p-value | log2FoldChange |
|---|---|---|---|---|---|
| Camta1 | 1.61e-07 | -0.801 | Atp5l | 2.28e-04 | 0.882 |
| Flrt3 | 1.94e-07 | 1.568 | Rbms3 | 2.45e-04 | -0.624 |
| Wdr31 | 1.99e-07 | -0.956 | Cntn1 | 2.51e-04 | -0.571 |

PEP1

| Gene Name | adjusted p-value | log2FoldChange | Gene Name | adjusted p-value | log2FoldChange |
|---|---|---|---|---|---|
| Ttf1 | 1.74e-35 | 1.780 | Ldlr | 2.82e-04 | 0.534 |
| Als2 | 3.52e-25 | 1.227 | Atp6v0b | 2.92e-04 | 0.835 |
| Ccdc62 | 7.39e-24 | 1.377 | Armt1 | 3.15e-04 | -1.048 |
| Hpgds | 2.30e-23 | 1.688 | Atp6ap1 | 3.55e-04 | 0.681 |
| Sprr1a | 1.44e-22 | 8.278 | Cox5b | 3.78e-04 | 0.788 |
| Golga3 | 1.85e-22 | 1.342 | Arhgdig | 4.97e-04 | 0.627 |
| Unc13d | 2.76e-22 | 1.419 | Irak1 | 5.20e-04 | 0.612 |
| Phkg1 | 9.15e-20 | 1.159 | Vwa5a | 5.84e-04 | -0.696 |
| BC051077 | 4.07e-19 | 1.301 | Arf5 | 6.43e-04 | 0.598 |
| Itch | 3.75e-18 | 1.294 | Fam102b | 7.24e-04 | -0.694 |
| Pycr2 | 1.33e-16 | 1.609 | Samd14 | 8.43e-04 | 0.519 |
| Exosc2 | 2.89e-16 | 1.389 | Esr1 | 9.94e-04 | 2.170 |
| Hbb-bs | 1.16e-14 | 2.145 | Adamts13 | 1.37e-03 | -1.775 |
| Bcas2 | 1.47e-14 | 0.930 | Pea15a | 1.46e-03 | 0.792 |
| Sag | 1.11e-13 | 1.656 | Atpaf2 | 1.50e-03 | 1.380 |
| Camta1 | 7.64e-13 | -0.789 | Cisd3 | 1.62e-03 | 1.016 |
| BC024386 | 2.10e-11 | 1.469 | AU022252 | 1.64e-03 | 0.735 |
| P3h3 | 7.06e-11 | 1.420 | Steap3 | 1.89e-03 | 1.349 |
| Samhd1 | 2.10e-10 | 1.054 | Pura | 1.90e-03 | -0.568 |
| Hoxd10 | 3.25e-10 | 2.773 | Cst3 | 2.12e-03 | 0.627 |
| Trim43b | 5.64e-10 | 1.568 | Tax1bp1 | 2.30e-03 | -1.111 |
| Npy | 5.96e-10 | 8.194 | Zfas1 | 2.36e-03 | 0.982 |
| Gpx3 | 6.07e-10 | 0.877 | Cflar | 2.46e-03 | 1.167 |
| Hbb-bt | 2.23e-09 | 2.046 | Uqcrq | 2.66e-03 | 0.822 |
| Ap1ar | 6.49e-09 | -1.274 | Dnajc7 | 3.44e-03 | 0.613 |
| Nup98 | 2.31e-08 | 1.487 | Enpp3 | 3.87e-03 | 0.995 |
| Nedd4 | 6.13e-08 | -0.828 | Zbtb20 | 4.52e-03 | -0.733 |
| Rab33b | 2.10e-07 | 1.135 | Magt1 | 7.37e-03 | 0.899 |
| Cd63 | 2.44e-07 | 1.386 | Adora3 | 8.11e-03 | 1.220 |
| Casc4 | 3.40e-07 | 1.086 | Tro | 8.47e-03 | 1.090 |
| Serinc1 | 6.61e-07 | -0.578 | Ahnak2 | 1.38e-02 | -0.560 |
| Ngfr | 1.42e-06 | 0.701 | Selenof | 1.44e-02 | -0.780 |

| Gene Name | adjusted p-value | log2FoldChange | Gene Name | adjusted p-value | log2FoldChange |
|---|---|---|---|---|---|
| Gcamp6 | 1.46e-06 | -6.684 | Clip4 | 1.46e-02 | -0.813 |
| Rpl38 | 2.25e-06 | 0.693 | Gabra3 | 1.62e-02 | 2.168 |
| Shc1 | 2.45e-06 | 0.965 | Cntnap2 | 1.81e-02 | -0.746 |
| Noct | 3.52e-06 | -0.718 | Rps23 | 1.81e-02 | -0.573 |
| Zwint | 5.91e-06 | -0.635 | Hba-a2 | 2.17e-02 | 1.737 |
| Selenow | 1.14e-05 | 0.873 | Jak1 | 2.20e-02 | 0.635 |
| Nr3c1 | 1.79e-05 | -0.956 | Slc23a1 | 2.87e-02 | 1.505 |
| Ptpro | 1.91e-05 | 2.096 | Gstm5 | 3.74e-02 | 0.954 |
| Ash2l | 2.77e-05 | 1.059 | Vgf | 3.96e-02 | 0.970 |
| Fgf13 | 2.88e-05 | -0.535 | Cd59a | 4.55e-02 | 0.910 |
| Chst2 | 5.72e-05 | 1.627 | Bscl2 | 4.85e-02 | 0.812 |
| Loxl2 | 5.93e-05 | 1.356 | | | |
| Rmi2 | 6.81e-05 | 0.954 | | | |
| Ap2s1 | 9.91e-05 | 0.864 | | | |
| Brap | 1.08e-04 | 0.878 | | | |
| Ralbp1 | 1.09e-04 | -0.948 | | | |
| Jund | 1.71e-04 | 0.550 | | | |
| Nts | 2.15e-04 | 4.413 | | | |

PEP2

| Gene Name | adjusted p-value | log2FoldChange | Gene Name | adjusted p-value | log2FoldChange |
|---|---|---|---|---|---|
| Ttf1 | 2.96e-18 | 1.785 | | | |
| BC051077 | 4.26e-11 | 1.663 | | | |
| Sprr1a | 5.35e-10 | 7.758 | | | |
| Hbb-bs | 2.61e-09 | 2.290 | | | |
| Samhd1 | 1.26e-07 | 1.496 | | | |
| Hpgds | 2.09e-07 | 1.476 | | | |
| Als2 | 5.99e-07 | 1.252 | | | |
| Bckdk | 2.02e-06 | 1.814 | | | |
| Bcas2 | 3.80e-06 | 1.158 | | | |
| Unc13d | 4.25e-06 | 1.361 | | | |
| Rab33b | 8.49e-06 | 1.411 | | | |
| Gpx3 | 1.46e-05 | 3.097 | | | |
| Tac1 | 3.21e-05 | 0.805 | | | |
| Phkg1 | 3.92e-05 | 1.167 | | | |
| Golga3 | 4.14e-05 | 1.134 | | | |
| Cst3 | 2.36e-04 | 1.409 | | | |
| Ccdc62 | 4.64e-04 | 1.183 | | | |

| Gene Name | adjusted p-value | log2FoldChange | Gene Name | adjusted p-value | log2FoldChange |
|---|---|---|---|---|---|
| Serinc1 | 7.76e-04 | -0.861 | | | |
| Scn7a | 1.57e-03 | -0.901 | | | |
| Was | 1.67e-03 | 1.827 | | | |
| Npy | 4.33e-03 | 7.455 | | | |
| Loxl2 | 5.64e-03 | 1.406 | | | |
| Itch | 7.07e-03 | 1.163 | | | |
| Glg1 | 2.84e-02 | -0.718 | | | |
| Tmem229b | 2.86e-02 | 1.504 | | | |
| P3h3 | 3.96e-02 | 1.073 | | | |

NP

| Gene Name | adjusted p-value | log2FoldChange | Gene Name | adjusted p-value | log2FoldChange |
|---|---|---|---|---|---|
| Sprr1a | 7.83e-29 | 9.466 | Sag | 5.59e-06 | 0.968 |
| Gcamp6 | 5.57e-25 | -9.649 | Pcsk1n | 5.70e-06 | 0.741 |
| Ap1ar | 5.35e-21 | -1.388 | Rps21 | 6.15e-06 | 0.856 |
| Tubb3 | 2.37e-19 | 0.600 | Pura | 1.51e-05 | -0.825 |
| Ttf1 | 6.00e-16 | 1.285 | Lgals1 | 1.65e-05 | 2.700 |
| Hbb-bs | 6.49e-15 | 1.730 | Ctsd | 1.69e-05 | 0.743 |
| Npy | 6.33e-14 | 8.771 | P3h3 | 2.32e-05 | 1.274 |
| Camta1 | 6.61e-13 | -0.820 | Fryl | 2.34e-05 | -0.945 |
| Gnaq | 1.48e-12 | -0.688 | Grina | 2.78e-05 | 0.608 |
| Gal | 3.05e-12 | 7.059 | Hsd17b12 | 3.41e-05 | 0.880 |
| S100a6 | 3.58e-12 | 0.606 | Tecr | 3.45e-05 | 0.855 |
| Bcam | 1.14e-11 | 0.886 | Synm | 6.22e-05 | -0.676 |
| Nts | 3.22e-11 | 4.868 | Hspb1 | 7.54e-05 | 1.605 |
| BC051077 | 4.75e-11 | 1.204 | Selenof | 7.88e-05 | -0.714 |
| Hpgds | 7.61e-11 | 1.267 | C2cd5 | 8.73e-05 | -1.460 |
| Syt11 | 5.75e-10 | -0.503 | Dpysl2 | 9.21e-05 | -0.577 |
| BC024386 | 1.43e-09 | 1.339 | Syap1 | 1.03e-04 | -3.719 |
| Armt1 | 1.85e-09 | -1.583 | Pmp22 | 1.23e-04 | 0.531 |
| Golga3 | 2.43e-09 | 1.028 | Ccser2 | 1.98e-04 | -0.630 |
| Samhd1 | 7.60e-09 | 1.134 | Tmem63b | 2.32e-04 | 0.854 |
| Noct | 8.77e-09 | -0.840 | Pycr2 | 3.86e-04 | 0.996 |
| Bcas2 | 1.51e-08 | 0.878 | Hba-a2 | 5.64e-04 | 1.798 |
| Rpl38 | 2.12e-08 | 0.897 | Slfn5 | 6.37e-04 | 1.146 |
| Selenow | 2.37e-08 | 1.264 | Cd55 | 6.68e-04 | -0.505 |
| Rps2 | 3.22e-08 | 1.129 | Snhg11 | 6.94e-04 | -1.006 |

| Gene Name | adjusted p-value | log2FoldChange | Gene Name | adjusted p-value | log2FoldChange |
|---|---|---|---|---|---|
| Ccdc62 | 3.93e-08 | 1.109 | Npc2 | 7.75e-04 | 1.215 |
| Itch | 4.53e-08 | 1.066 | Ebf1 | 8.61e-04 | -0.574 |
| Phkg1 | 9.46e-08 | 0.964 | Cap1 | 9.27e-04 | 1.316 |
| Pkm | 9.69e-08 | 0.763 | Magt1 | 1.15e-03 | 1.007 |
| Grik1 | 9.85e-08 | -0.867 | Tle1 | 1.17e-03 | -0.703 |
| Apod | 1.09e-07 | 2.056 | Tshz2 | 1.32e-03 | -0.614 |
| Chchd2 | 2.06e-07 | 0.632 | Etnk1 | 1.42e-03 | -0.525 |
| Unc13d | 2.86e-07 | 0.965 | AU022252 | 1.59e-03 | 0.773 |
| Als2 | 4.04e-07 | 0.899 | Cacna2d2 | 1.61e-03 | -1.911 |
| Rab33b | 4.21e-07 | 0.969 | Scn9a | 1.62e-03 | -0.731 |
| Stmn3 | 4.74e-07 | 0.637 | Trim43b | 2.71e-03 | 1.073 |
| Thy1 | 5.32e-07 | 1.688 | Cflar | 3.11e-03 | 0.958 |
| Rbfox1 | 5.53e-07 | -0.700 | Kcnab2 | 3.23e-03 | 0.664 |
| Gap43 | 6.36e-07 | 1.238 | Rps11 | 3.40e-03 | 0.757 |
| Nnat | 7.00e-07 | 1.211 | Sept5 | 3.57e-03 | 0.704 |
| Exosc2 | 7.27e-07 | 1.017 | Atp6v0b | 4.07e-03 | 0.651 |
| Hsd3b2 | 9.70e-07 | -1.457 | Syp | 4.17e-03 | 0.724 |
| Atf3 | 1.13e-06 | 4.411 | Mvd | 4.18e-03 | 1.382 |
| Serinc1 | 1.78e-06 | -0.683 | Nme2 | 4.75e-03 | 1.079 |
| Hbb-bt | 1.91e-06 | 1.939 | Zbtb20 | 5.67e-03 | -1.513 |
| Pten | 2.48e-06 | -0.994 | Tkt | 7.41e-03 | 0.686 |
| Rgs7bp | 2.72e-06 | -0.672 | Cdc37 | 7.59e-03 | 0.702 |
| Eef1a2 | 2.87e-06 | 0.572 | Homer2 | 7.95e-03 | -1.171 |
| G3bp2 | 4.41e-06 | -0.773 | Jhy | 8.24e-03 | -2.424 |
| Aqp1 | 5.20e-06 | 0.793 | Rbms1 | 9.24e-03 | -0.547 |

SST

| Gene Name | adjusted p-value | log2FoldChange | Gene Name | adjusted p-value | log2FoldChange |
|---|---|---|---|---|---|
| Ttf1 | 8.56e-05 | 2.282 | | | |
| Hpgds | 2.60e-03 | 2.808 | | | |
| Phkg1 | 6.32e-03 | 1.806 | | | |

p-cLTMR2

| Gene Name | adjusted p-value | log2FoldChange | Gene Name | adjusted p-value | log2FoldChange |
|---|---|---|---|---|---|
| Sprr1a | 1.57e-18 | 9.407 | | | |
| Gal | 1.13e-08 | 8.674 | | | |
| Hbb-bs | 2.72e-07 | 1.559 | | | |

| Gene Name | adjusted p-value | log2FoldChange | Gene Name | adjusted p-value | log2FoldChange |
|---|---|---|---|---|---|
| Npy | 1.89e-06 | 8.272 | | | |
| Ttf1 | 9.88e-06 | 1.488 | | | |
| Pycr2 | 7.03e-05 | 1.963 | | | |
| Tubb3 | 2.79e-04 | 0.792 | | | |
| Ccdc62 | 3.17e-04 | 1.492 | | | |
| Wdr89 | 5.52e-04 | 1.670 | | | |
| Hspa8 | 3.15e-03 | 0.705 | | | |
| Ap1ar | 3.68e-03 | -1.187 | | | |
| Hpgds | 3.89e-03 | 1.394 | | | |
| Hba-a2 | 4.64e-03 | 2.049 | | | |
| Phkg1 | 4.75e-03 | 1.247 | | | |
| Als2 | 9.64e-03 | 1.116 | | | |
| Cpne3 | 1.17e-02 | -0.588 | | | |
| Tubb5 | 1.62e-02 | 0.548 | | | |
| P3h3 | 2.14e-02 | 2.039 | | | |
| BC051077 | 2.38e-02 | 1.271 | | | |
| Sag | 2.85e-02 | 1.343 | | | |
| Abtb2 | 3.87e-02 | -1.233 | | | |

Injured

| Gene Name | adjusted p-value | log2FoldChange | Gene Name | adjusted p-value | log2FoldChange |
|---|---|---|---|---|---|
| Sox11 | 6.98e-273 | 3.643 | Atp1a1 | 8.84e-78 | -1.176 |
| Sprr1a | 1.25e-256 | 3.493 | Calcb | 1.05e-77 | -2.675 |
| Flrt3 | 8.23e-253 | 3.457 | Sdc1 | 3.96e-77 | 2.771 |
| Atf3 | 2.48e-234 | 3.824 | Apba2 | 9.35e-77 | 2.091 |
| Stmn2 | 3.01e-204 | 1.135 | Kcnb1 | 2.85e-76 | -2.203 |
| Cdkn1a | 1.80e-193 | 2.605 | S100a11 | 1.07e-75 | 1.986 |
| Itga7 | 2.54e-192 | 2.579 | Ubash3b | 2.12e-75 | -1.597 |
| Nrn1 | 1.08e-178 | -2.998 | Nmb | 1.27e-74 | -2.705 |
| Sema6a | 3.27e-177 | 3.464 | Tmem233 | 2.54e-74 | -2.972 |
| Stmn4 | 7.50e-173 | 3.141 | Ldb2 | 4.22e-74 | -2.656 |
| Rhoq | 1.58e-172 | 2.260 | Pde11a | 5.07e-74 | -3.388 |
| Gadd45a | 1.12e-154 | 2.767 | Phlda1 | 1.71e-73 | 2.135 |
| Chst2 | 2.04e-142 | 2.636 | S100a10 | 6.71e-73 | 1.149 |
| Cacna2d1 | 3.69e-138 | 1.733 | Rasgrf2 | 5.32e-72 | 2.608 |
| Atp1b3 | 1.56e-136 | -2.172 | Nrip1 | 8.96e-72 | 2.010 |
| Gap43 | 1.66e-128 | 1.616 | Kcna2 | 1.19e-71 | -2.273 |
| Gal | 3.36e-128 | 3.196 | Eml1 | 7.31e-71 | -1.959 |
| Cd9 | 4.31e-125 | -1.722 | Kif22 | 9.20e-71 | 2.919 |

| Gene Name | adjusted p-value | log2FoldChange | Gene Name | adjusted p-value | log2FoldChange |
|---|---|---|---|---|---|
| Tubb3 | 5.26e-124 | 0.869 | Fez1 | 9.67e-71 | -1.716 |
| Tubb2b | 1.22e-122 | 1.272 | Myo10 | 1.38e-69 | 2.747 |
| Nsg1 | 3.65e-122 | 2.504 | Parm1 | 2.39e-69 | -1.298 |
| Scn10a | 5.19e-122 | -3.380 | Thy1 | 1.22e-67 | 0.982 |
| Fstl1 | 1.62e-120 | -1.282 | Scn11a | 1.91e-67 | -2.893 |
| Sez6l | 1.93e-120 | 4.038 | Ncald | 4.16e-67 | -2.316 |
| Csf1 | 2.77e-120 | 3.346 | Tmem88b | 8.82e-67 | 3.826 |
| Tubb6 | 3.13e-120 | 3.252 | Lmo7 | 1.81e-66 | 3.122 |
| Hn1 | 4.93e-117 | 1.678 | Cckbr | 2.15e-66 | 3.633 |
| Srxn1 | 1.23e-116 | 1.879 | Sorl1 | 4.03e-66 | -2.536 |
| Cd63 | 3.64e-116 | 2.180 | Clic1 | 4.99e-66 | 1.382 |
| Abca1 | 2.46e-112 | 2.465 | Trib2 | 6.07e-66 | -3.083 |
| Calca | 1.40e-103 | -2.739 | Rtn4 | 8.09e-66 | 0.939 |
| Chl1 | 1.12e-99 | 2.568 | Mthfd2 | 1.12e-65 | 2.057 |
| Tnik | 2.16e-99 | 2.734 | Dgkz | 1.74e-64 | -2.037 |
| Tuba1a | 1.50e-96 | 0.745 | Nap1l5 | 6.83e-64 | -1.699 |
| Hspb1 | 1.96e-95 | 1.469 | Clic4 | 1.16e-63 | 2.234 |
| Gnao1 | 1.85e-94 | -1.152 | Igfbp3 | 2.85e-63 | 3.176 |
| Jun | 2.08e-93 | 1.772 | Scg3 | 1.76e-62 | -1.820 |
| Mmp16 | 5.84e-92 | 3.329 | Scg2 | 2.24e-62 | -2.308 |
| Tac1 | 1.95e-91 | -3.853 | Prkca | 1.19e-61 | -2.180 |
| Phf24 | 1.44e-90 | -2.268 | Ecel1 | 7.19e-61 | 3.745 |
| Pirt | 2.68e-88 | -1.629 | Cadm1 | 1.50e-60 | -1.293 |
| Fgf13 | 1.89e-85 | -1.854 | Elavl3 | 3.30e-60 | 1.066 |
| Trp53i11 | 8.84e-85 | 3.877 | Fxyd2 | 1.76e-59 | -1.858 |
| Irs2 | 1.17e-84 | 1.765 | Npy | 2.06e-58 | 3.556 |
| Mal2 | 1.38e-84 | -2.755 | Glul | 2.83e-58 | 1.420 |
| Ppp1r1c | 2.31e-84 | -2.094 | Eepd1 | 2.85e-58 | -2.502 |
| Tusc5 | 5.15e-83 | -2.013 | Fam171b | 1.39e-57 | 1.324 |
| Camk1 | 9.67e-82 | 2.311 | Rasgrp1 | 3.04e-57 | -4.352 |
| Nts | 2.03e-81 | 2.615 | Kcnab2 | 4.66e-57 | -1.050 |
| Rhoc | 5.40e-79 | 2.071 | Tmem255a | 8.11e-57 | -2.623 |

**Appendix Table 7:** DRG ScNT Day 28 top significant neuronal DEGs. Top 100 per subtype; adjusted p-value < 0.05; |log2FC| > 0.5.

NF

| Gene Name | adjusted p-value | log2FoldChange | Gene Name | adjusted p-value | log2FoldChange |
|---|---|---|---|---|---|
| Vgf | 8.58e-12 | 1.528 | | | |
| Npy | 1.16e-08 | 5.650 | | | |
| Phkg1 | 3.09e-08 | 1.589 | | | |
| Tmsb10 | 5.94e-07 | 1.301 | | | |
| Scg5 | 4.41e-06 | -1.894 | | | |
| Sprr1a | 6.93e-06 | 5.431 | | | |
| Nts | 1.01e-04 | 3.055 | | | |
| Gap43 | 1.12e-04 | 0.806 | | | |
| Als2 | 1.73e-04 | 1.538 | | | |
| Vamp2 | 6.71e-04 | 0.732 | | | |
| Ccdc62 | 1.01e-03 | 1.188 | | | |
| Zc3hc1 | 1.04e-03 | -1.309 | | | |
| Fxyd7 | 2.51e-03 | 0.628 | | | |
| Tecr | 4.51e-03 | 0.658 | | | |
| Palm | 5.90e-03 | 1.232 | | | |
| S100a11 | 1.02e-02 | 1.513 | | | |
| Ngfr | 2.42e-02 | 0.563 | | | |
| Fscn1 | 2.93e-02 | 1.961 | | | |
| Etv5 | 3.44e-02 | 1.096 | | | |
| Rac1 | 3.82e-02 | 0.615 | | | |

PEP1

| Gene Name | adjusted p-value | log2FoldChange | Gene Name | adjusted p-value | log2FoldChange |
|---|---|---|---|---|---|
| Zc3hc1 | 2.23e-16 | -1.526 | | | |
| Tubb4b | 3.05e-14 | -0.606 | | | |
| Als2 | 1.51e-13 | 0.998 | | | |
| Phkg1 | 4.30e-12 | 1.005 | | | |
| Hsp90aa1 | 1.55e-11 | -0.537 | | | |
| Nefl | 4.90e-11 | -0.634 | | | |
| Socs2 | 4.75e-07 | 1.169 | | | |
| Ntrk1 | 6.27e-07 | -0.681 | | | |
| Vgf | 1.02e-06 | 1.334 | | | |
| Unc13d | 1.35e-06 | 0.924 | | | |
| Golga3 | 8.15e-06 | 0.864 | | | |
| Gcamp6 | 1.13e-05 | -7.011 | | | |

| **Gene Name** | **adjusted p-value** | **log2FoldChange** | **Gene Name** | **adjusted p-value** | **log2FoldChange** |
|---|---|---|---|---|---|
| Tkt | 1.19e-05 | -0.790 | | | |
| Pappa | 1.37e-05 | 1.447 | | | |
| Ly6e | 2.73e-05 | 1.924 | | | |
| Car8 | 4.39e-05 | 0.793 | | | |
| Adamts13 | 4.86e-05 | -1.957 | | | |
| Irgm1 | 3.30e-04 | 0.895 | | | |
| Npy1r | 4.08e-04 | 0.873 | | | |
| Vamp2 | 5.53e-04 | 0.507 | | | |
| Ly6a | 6.01e-04 | 4.166 | | | |
| Tmsb10 | 6.70e-04 | 0.719 | | | |
| Vat1 | 1.00e-03 | -0.539 | | | |
| Flrt3 | 1.52e-03 | 1.522 | | | |
| Wipi2 | 1.66e-03 | -0.548 | | | |
| Marcks | 2.33e-03 | -0.564 | | | |
| Rpsa | 2.39e-03 | -0.524 | | | |
| Gatad2b | 2.88e-03 | 0.855 | | | |
| Rab11fip5 | 4.50e-03 | 1.008 | | | |
| Selenow | 5.91e-03 | 0.629 | | | |
| Rpl17 | 5.95e-03 | -0.706 | | | |
| BC024386 | 5.96e-03 | 0.821 | | | |
| Nefm | 6.36e-03 | -0.508 | | | |
| Cd44 | 6.42e-03 | 0.594 | | | |
| Pde6d | 1.01e-02 | 0.639 | | | |
| Mbnl1 | 1.16e-02 | -0.554 | | | |
| Sod1 | 1.23e-02 | -0.689 | | | |
| Psip1 | 1.49e-02 | -0.616 | | | |
| Bdnf | 1.68e-02 | 0.794 | | | |
| Scg5 | 1.71e-02 | -1.031 | | | |
| Vcl | 3.26e-02 | 1.235 | | | |
| Abca1 | 3.32e-02 | 1.019 | | | |
| Itga3 | 3.81e-02 | 0.887 | | | |
| Psmb5 | 4.33e-02 | -0.861 | | | |
| Rbfox1 | 4.95e-02 | -0.534 | | | |

PEP2

| **Gene Name** | **adjusted p-value** | **log2FoldChange** | **Gene Name** | **adjusted p-value** | **log2FoldChange** |
|---|---|---|---|---|---|
| Als2 | 1.72e-02 | 1.251 | | | |
| Phkg1 | 2.61e-02 | 1.133 | | | |

NP

| Gene Name | adjusted p-value | log2FoldChange | Gene Name | adjusted p-value | log2FoldChange |
|---|---|---|---|---|---|
| Gcamp6 | 3.29e-16 | -9.064 | | | |
| Ly86 | 9.37e-10 | 0.923 | | | |
| Lin54 | 4.22e-08 | -0.629 | | | |
| Zc3hc1 | 4.79e-08 | -0.929 | | | |
| Tmem233 | 7.43e-08 | 0.512 | | | |
| Cd44 | 2.16e-05 | 0.519 | | | |
| Tmsb10 | 4.43e-05 | 0.792 | | | |
| Cacna2d2 | 5.92e-05 | -2.473 | | | |
| Als2 | 6.33e-05 | 1.065 | | | |
| Magt1 | 1.88e-04 | 1.350 | | | |
| Mme | 2.15e-04 | -0.533 | | | |
| Adamts13 | 2.28e-04 | -1.989 | | | |
| Snrpn | 2.74e-04 | 0.528 | | | |
| Hpgds | 5.16e-04 | 1.036 | | | |
| Scg5 | 7.83e-04 | -2.040 | | | |
| Ccdc6 | 3.35e-03 | 1.788 | | | |
| Golga3 | 2.40e-02 | 1.007 | | | |
| Rcc2 | 3.65e-02 | -1.293 | | | |
| Phkg1 | 4.30e-02 | 1.101 | | | |
| Syngr3 | 4.77e-02 | 0.605 | | | |
| Ankrd24 | 4.84e-02 | 1.352 | | | |

SST

| Gene Name | adjusted p-value | log2FoldChange | Gene Name | adjusted p-value | log2FoldChange |
|---|---|---|---|---|---|
| Stmn4 | 3.60e-08 | 7.848 | | | |
| Gap43 | 7.40e-08 | 3.198 | | | |
| Hspb1 | 6.15e-07 | 2.903 | | | |
| Cdhr1 | 1.31e-04 | 2.224 | | | |
| Flrt3 | 5.45e-04 | 3.169 | | | |
| Gfra3 | 9.82e-04 | 1.881 | | | |
| Cd44 | 1.51e-03 | 2.257 | | | |
| S100a11 | 3.46e-03 | 2.936 | | | |
| Sox11 | 6.32e-03 | 3.668 | | | |
| Sst | 1.06e-02 | -1.172 | | | |
| Npy1r | 1.49e-02 | 5.380 | | | |
| Ly6e | 3.00e-02 | 6.580 | | | |
| Lrrc15 | 4.19e-02 | 5.762 | | | |

| Gene Name | adjusted p-value | log2FoldChange | Gene Name | adjusted p-value | log2FoldChange |
|---|---|---|---|---|---|
| Lynx1 | 4.58e-02 | 2.309 | | | |
| Adora3 | 4.72e-02 | 5.105 | | | |

p-cLTMR2

| Gene Name | adjusted p-value | log2FoldChange | Gene Name | adjusted p-value | log2FoldChange |
|---|---|---|---|---|---|
| No significant DEGs | | | | | |

Injured

| Gene Name | adjusted p-value | log2FoldChange | Gene Name | adjusted p-value | log2FoldChange |
|---|---|---|---|---|---|
| Sprr1a | 1.23e-139 | 4.993 | Jun | 2.93e-40 | 1.497 |
| Stmn2 | 1.21e-111 | 1.209 | Cadm1 | 2.30e-39 | -1.583 |
| Atf3 | 3.40e-106 | 4.242 | Osbpl3 | 6.04e-39 | -1.639 |
| Nts | 1.46e-98 | 3.820 | Cbln2 | 1.26e-38 | 3.640 |
| Flrt3 | 3.02e-91 | 2.857 | Nedd4l | 1.34e-38 | -1.840 |
| Chst2 | 5.38e-86 | 3.114 | Nmb | 1.37e-38 | -3.114 |
| Npy | 2.14e-83 | 5.568 | Gfra1 | 2.94e-38 | 2.392 |
| Nsg1 | 4.95e-79 | 2.465 | Nwd2 | 7.50e-38 | -2.779 |
| Sema6a | 6.59e-79 | 4.102 | Chl1 | 1.01e-37 | 1.957 |
| Sox11 | 3.51e-78 | 2.657 | Car3 | 1.43e-37 | 4.692 |
| Tubb3 | 1.62e-77 | 1.130 | Gap43 | 2.59e-37 | 1.382 |
| Thy1 | 3.68e-70 | 1.384 | Calcb | 3.45e-37 | -2.759 |
| Srxn1 | 1.72e-67 | 2.086 | Cacna2d1 | 6.87e-37 | 1.257 |
| Rasgrf2 | 2.68e-67 | 3.646 | Mcam | 9.43e-37 | 1.545 |
| Calca | 4.42e-67 | -4.434 | Ahnak2 | 1.17e-36 | 1.039 |
| Scn10a | 3.11e-63 | -3.689 | Eif1b | 2.74e-36 | 1.392 |
| Kcnb1 | 9.60e-62 | -2.875 | Abca1 | 9.71e-36 | 1.940 |
| Sez6l | 7.28e-61 | 4.157 | Prkcd | 1.05e-35 | -3.037 |
| Tusc5 | 4.79e-60 | -2.922 | Fxyd2 | 2.52e-35 | -2.634 |
| Pirt | 1.21e-57 | -1.849 | Atp8a1 | 9.39e-35 | -2.003 |
| Scn11a | 2.96e-57 | -4.596 | Tmem233 | 1.11e-34 | -3.584 |
| Tubb6 | 7.90e-53 | 3.228 | Timp3 | 2.25e-34 | 2.682 |
| Rhoq | 1.64e-52 | 1.961 | Rtn4 | 9.18e-34 | 0.891 |
| Tnik | 3.01e-52 | 2.773 | Tubb2a | 2.49e-33 | 0.854 |
| Tuba1b | 5.07e-52 | 0.951 | Ldb2 | 3.05e-33 | -3.081 |
| Stmn4 | 3.17e-51 | 2.541 | Irs2 | 3.46e-33 | 1.749 |
| Fgf13 | 4.34e-51 | -1.879 | Fstl1 | 4.71e-33 | -0.971 |

| Gene Name | adjusted p-value | log2FoldChange | Gene Name | adjusted p-value | log2FoldChange |
|---|---|---|---|---|---|
| Tuba1a | 1.32e-49 | 0.746 | Sncb | 7.47e-33 | 1.756 |
| Gnao1 | 1.92e-49 | -1.144 | Igfbp3 | 7.96e-33 | 3.437 |
| Cckbr | 6.71e-49 | 4.497 | Kbtbd11 | 1.34e-32 | -4.577 |
| Itga7 | 9.27e-49 | 1.903 | Deptor | 3.27e-32 | -2.103 |
| Atp1b3 | 2.42e-48 | -1.935 | Prss12 | 3.28e-32 | 3.053 |
| Slc24a2 | 4.11e-46 | 1.596 | F2rl2 | 1.97e-31 | -3.652 |
| Gpr153 | 1.21e-45 | 2.122 | Cplx1 | 4.61e-31 | 1.159 |
| Gabrg1 | 1.38e-45 | 3.953 | Ppp1r2 | 6.48e-31 | -1.415 |
| Tshz2 | 2.59e-45 | -1.807 | Clu | 1.23e-30 | 1.653 |
| Sparcl1 | 3.68e-45 | 2.214 | Hspb1 | 2.57e-30 | 1.403 |
| Gpr151 | 5.31e-45 | 4.851 | Gnaq | 4.15e-30 | -1.304 |
| Cd9 | 1.04e-44 | -1.767 | Csf1 | 7.13e-30 | 2.961 |
| Cdkn1a | 1.73e-44 | 2.185 | Ctsb | 1.70e-29 | 0.770 |
| Tac1 | 2.74e-44 | -4.142 | Prkce | 2.00e-29 | -1.342 |
| Mal2 | 6.88e-44 | -3.204 | Dgkz | 2.08e-29 | -2.305 |
| Eml1 | 7.12e-44 | -2.397 | Csrp1 | 2.48e-29 | 1.998 |
| Fam171b | 4.41e-42 | 1.486 | Rhoc | 6.89e-29 | 2.089 |
| Nrn1 | 1.15e-41 | -2.574 | Star | 4.07e-28 | 4.974 |
| Ubash3b | 2.28e-41 | -1.530 | Zmat4 | 6.04e-28 | -3.286 |
| Gadd45a | 2.69e-41 | 2.372 | Ptprt | 2.12e-27 | -3.075 |
| Itpr1 | 3.30e-41 | 3.303 | Scg2 | 2.33e-27 | -2.402 |
| Pde11a | 4.16e-41 | -2.798 | Cntn1 | 2.61e-27 | 0.910 |
| Rbms1 | 1.52e-40 | -1.281 | Phf24 | 3.85e-27 | -2.021 |

**Appendix References:**


Ahlmann-Eltze C, Huber W. 2021. Glmgampoi: Fitting gamma-poisson generalized linear models on single cell count data. Bioinformatics. 36(24):5701-5702. https://www.ncbi.nlm.nih.gov/pubmed/33295604. doi:10.1093/bioinformatics/btaa1009.

Alkaslasi MR et al. 2021. Single nucleus rna-sequencing defines unexpected diversity of cholinergic neuron types in the adult mouse spinal cord. Nat Commun. 12(1):2471. https://www.ncbi.nlm.nih.gov/pubmed/33931636. doi:10.1038/s41467-021-22691-2.

Austin PJ, Wu A, Moalem-Taylor G. 2012. Chronic constriction of the sciatic nerve and pain hypersensitivity testing in rats. J Vis Exp. 61). https://www.ncbi.nlm.nih.gov/pubmed/22433911. doi:10.3791/3393.

Bennett GJ, Xie YK. 1988. A peripheral mononeuropathy in rat that produces disorders of pain sensation like those seen in man. Pain. 33(1):87-107. https://www.ncbi.nlm.nih.gov/pubmed/2837713. doi:10.1016/0304-3959(88)90209-6.

Finak G et al. 2015. Mast: A flexible statistical framework for assessing transcriptional changes and characterizing heterogeneity in single-cell rna sequencing data. Genome Biol. 16(278. https://www.ncbi.nlm.nih.gov/pubmed/26653891. doi:10.1186/s13059-015-0844-5.

Garcia-Alonso L, Holland CH, Ibrahim MM, Turei D, Saez-Rodriguez J. 2019. Benchmark and integration of resources for the estimation of human transcription factor activities. Genome Res. 29(8):1363-1375. https://www.ncbi.nlm.nih.gov/pubmed/31340985. doi:10.1101/gr.240663.118.

Hao Y et al. 2024. Dictionary learning for integrative, multimodal and scalable single-cell analysis. Nat Biotechnol. 42(2):293-304. https://www.ncbi.nlm.nih.gov/pubmed/37231261. doi:10.1038/s41587-023-01767-y.

Hu G et al. 2016. Single-cell rna-seq reveals distinct injury responses in different types of drg sensory neurons. Sci Rep. 6(31851. https://www.ncbi.nlm.nih.gov/pubmed/27558660. doi:10.1038/srep31851.

Jin S et al. 2021. Inference and analysis of cell-cell communication using cellchat. Nat Commun. 12(1):1088. https://www.ncbi.nlm.nih.gov/pubmed/33597522. doi:10.1038/s41467-021-21246-9.

Keddie S et al. 2023. Peripherin is a biomarker of axonal damage in peripheral nervous system disease. Brain. 146(11):4562-4573. https://www.ncbi.nlm.nih.gov/pubmed/37435933. doi:10.1093/brain/awad234.

Koike T et al. 2025. Cd34 distribution in c-fiber low threshold mechanoreceptors in the mouse dorsal root ganglion and spinal cord. Brain Res. 1846(149227. https://www.ncbi.nlm.nih.gov/pubmed/39255903. doi:10.1016/j.brainres.2024.149227.

Korczeniewska OA et al. 2024. Time-course progression of whole transcriptome expression changes of trigeminal ganglia compared to dorsal root ganglia in rats exposed to nerve injury. J Pain. 25(1):101-117. https://www.ncbi.nlm.nih.gov/pubmed/37524222. doi:10.1016/j.jpain.2023.07.024.

Korsunsky I et al. 2019. Fast, sensitive and accurate integration of single-cell data with harmony. Nat Methods. 16(12):1289-1296. https://www.ncbi.nlm.nih.gov/pubmed/31740819. doi:10.1038/s41592-019-0619-0.

Ma S et al. 2025. Preventive effect of rage antagonists on distal infraorbital nerve injury-induced pain behaviors of male and female mice. Biochem Pharmacol. 242(Pt 1):117242. https://www.ncbi.nlm.nih.gov/pubmed/40818549. doi:10.1016/j.bcp.2025.117242.

Mulier M et al. 2020. Upregulation of trpm3 in nociceptors innervating inflamed tissue. Elife. 9(https://www.ncbi.nlm.nih.gov/pubmed/32880575. doi:10.7554/eLife.61103.

Nascimento AI et al. 2022. Sensory neurons have an axon initial segment that initiates spontaneous activity in neuropathic pain. Brain. 145(5):1632-1640. https://www.ncbi.nlm.nih.gov/pubmed/35661858. doi:10.1093/brain/awac078.

Oschipok LW, Teh J, McPhail LT, Tetzlaff W. 2008. Expression of semaphorin3c in axotomized rodent facial and rubrospinal neurons. Neurosci Lett. 434(1):113-118. https://www.ncbi.nlm.nih.gov/pubmed/18308469. doi:10.1016/j.neulet.2008.01.048.

Ruparel SB, Akopian AN. 2024. Editorial: Mechanisms of orofacial pain. Front Pain Res (Lausanne). 5(1496188. https://www.ncbi.nlm.nih.gov/pubmed/39618991. doi:10.3389/fpain.2024.1496188.

Shankar Kikkeri N, Nagalli S. Trigeminal neuralgia. In: Statpearls.

Shiers S, Klein RM, Price TJ. 2020. Quantitative differences in neuronal subpopulations between mouse and human dorsal root ganglia demonstrated with rnascope in situ hybridization. Pain. 161(10):2410-2424. https://www.ncbi.nlm.nih.gov/pubmed/32639368. doi:10.1097/j.pain.0000000000001973.

Techameena P, Feng X, Zhang K, Hadjab S. 2024. The single-cell transcriptomic atlas ipain identifies senescence of nociceptors as a therapeutical target for chronic pain treatment. Nat Commun. 15(1):8585. https://www.ncbi.nlm.nih.gov/pubmed/39362841. doi:10.1038/s41467-024-52052-8.

Temmermand R, Barrett JE, Fontana ACK. 2022. Glutamatergic systems in neuropathic pain and emerging non-opioid therapies. Pharmacol Res. 185(106492. https://www.ncbi.nlm.nih.gov/pubmed/36228868. doi:10.1016/j.phrs.2022.106492.

Tilley DM, Cedeno DL, Vetri F, Platt DC, Vallejo R. 2022. Differential target multiplexed spinal cord stimulation programming modulates proteins involved in ion regulation in an animal model of neuropathic pain. Mol Pain. 18(17448069211060181. https://www.ncbi.nlm.nih.gov/pubmed/35048719. doi:10.1177/17448069211060181.

Treutlein EM et al. 2018. The prostaglandin e2 receptor ep3 controls cc-chemokine ligand 2-mediated neuropathic pain induced by mechanical nerve damage. J Biol Chem. 293(25):9685-9695. https://www.ncbi.nlm.nih.gov/pubmed/29752406. doi:10.1074/jbc.RA118.002492.

Vos BP, Strassman AM, Maciewicz RJ. 1994. Behavioral evidence of trigeminal neuropathic pain following chronic constriction injury to the rat's infraorbital nerve. J Neurosci. 14(5 Pt 1):2708-2723. https://www.ncbi.nlm.nih.gov/pubmed/8182437. doi:10.1523/JNEUROSCI.14-05-02708.1994.

Wang H et al. 2025. Suppression of schwann cell ferroptosis mitigates demyelination and trigeminal neuropathic pain in the ion-cci model. Eur J Neurosci. 62(3):e70209. https://www.ncbi.nlm.nih.gov/pubmed/40780703. doi:10.1111/ejn.70209.

Wu T et al. 2021. Clusterprofiler 4.0: A universal enrichment tool for interpreting omics data. Innovation (Camb). 2(3):100141. https://www.ncbi.nlm.nih.gov/pubmed/34557778. doi:10.1016/j.xinn.2021.100141.

Yamanaka H et al. 2007. Alteration of the cell adhesion molecule l1 expression in a specific subset of primary afferent neurons contributes to neuropathic pain. Eur J Neurosci. 25(4):1097-1111. https://www.ncbi.nlm.nih.gov/pubmed/17331206. doi:10.1111/j.1460-9568.2007.05344.x.

Zakrzewska JM. 2013. Multi-dimensionality of chronic pain of the oral cavity and face. J Headache Pain. 14(1):37. https://www.ncbi.nlm.nih.gov/pubmed/23617409. doi:10.1186/1129-2377-14-37.

Zappia L, Oshlack A. 2018. Clustering trees: A visualization for evaluating clusterings at multiple resolutions. Gigascience. 7(7). https://www.ncbi.nlm.nih.gov/pubmed/30010766. doi:10.1093/gigascience/giy083.
